# *In vivo* generation of chimeric antigen receptor T cells using optimally retargeted and functionalized lentiviral vectors with reduced immune clearance

**DOI:** 10.1101/2025.04.23.645810

**Authors:** Kepler S. Mears, Kyrellos Ibrahim, Peter M. Allen, Jordan M. Chinai, Omar I. Avila, Audrey J. Muscato, Sarah K. Lane-Reticker, Alexander Rojas, Nelson H. Knudsen, Chun-Cheih Chao, Kathleen B. Yates, Robert T. Manguso

## Abstract

Despite striking efficacy against hematologic malignancies, the cost and complexity of CAR T manufacturing present significant barriers to broader patient access. Beyond manufacturing challenges, *ex vivo* expansion of T cells may be detrimental for their function and persistence. Thus, delivery of CARs to reprogram host cells *in vivo* would represent a significant advance toward “off-the-shelf” therapy but has been limited by low efficiency, low specificity, and immunogenicity of viral vectors. Here we describe the design of pseudotyped lentiviral vectors (LV) with superior functionality and high target specificity. We show that LV pseudotyped with chimeric envelope glycoproteins from dolphin morbillivirus (DMV) can be engineered to selectively infect human T cells and evade neutralizing antibody responses in measles-vaccinated human serum. We further demonstrate that camelid-derived nanobodies are a superior retargeting domain, overcoming limitations inherent to the use of single chain variable fragment antibodies. Using a chimeric DMV-pseudotyped virus targeting the CD7 receptor, we demonstrate efficient and highly specific infection of T cells both *in vitro* and *in vivo*, generating functional CAR T cells and inducing therapeutic efficacy in a preclinical B cell lymphoma model.

## Introduction

Efficient and selective genetic reprogramming of host cells *in vivo* is a major goal for cell and gene therapy. Replication-incompetent lentiviral-like particles (LVs) have been extensively developed for genetic and protein delivery applications for both research and therapeutic applications^1–3^. LVs are enveloped vectors and will incorporate any transmembrane proteins from the producer cell on their surface. The tropism of LVs can be modified through pseudotyping: heterologously expressing different viral-derived and engineered proteins that are incorporated into the envelope during production^4–7^. Pseudotyping most commonly employs the vesicular stomatitis virus glycoprotein (VSV-G), which binds to the widely expressed low-density lipoprotein receptor (LDLR), providing broad tropism for many cell types and mammalian species. VSV-G pseudotyped LVs are currently used to generate chimeric antigen receptor (CAR) T cells *ex vivo* to treat hematological cancers. However, VSV-G provides minimal cell type selectivity and has been difficult to engineer for specificity, limiting its *in vivo* utility^8,9^.

Specific retargeting of LVs can be achieved through direct engineering of the pseudotype by the addition of a cell-selective targeting domain and has been successfully deployed to selectively transduce lymphocyte subsets^7,8,10–14^. The Measles Virus (MeV) envelope proteins have been demonstrated to be particularly amenable for retargeting LVs to a broad array of targets^15–17^. Measles, and all other paramyxoviruses, utilize a tetrameric hemagglutinin (H) protein for target receptor recognition and a separate trimeric fusion (F) protein to mediate membrane fusion^18,19^ (**Figure 1A**). Receptor recognition by the H protein triggers fusion to the target cell membrane through the F protein, a mechanism that bypasses the need for endocytosis of the viral particle that is a requirement for many viral Envs. By fusing a targeting domain to the H protein, the receptor preference can be altered. Residues in MeV-H can be further mutated to abolish binding to its natural receptors (SLAMF1 and NECTIN4), preventing aberrant transduction on non-target cells. The MeV Env has been demonstrated to provide highly selective targeting to many targets including those that are not endocytosed^8,20,21^. Single chain variable fragments (scFvs) derived from antibodies, DARPins, and cytokines have all been successfully implemented as targeting domains (**Figure 1A, middle**)^9,12,22–24^. Fusion of MeV-H to scFvs and DARPins targeting human CD8 or CD4 have been used to transduce T cells *in vitro,* and to generate functional CAR T cells to treat B cell lymphomas in humanized mouse models^10,11^.

**Figure 1:**
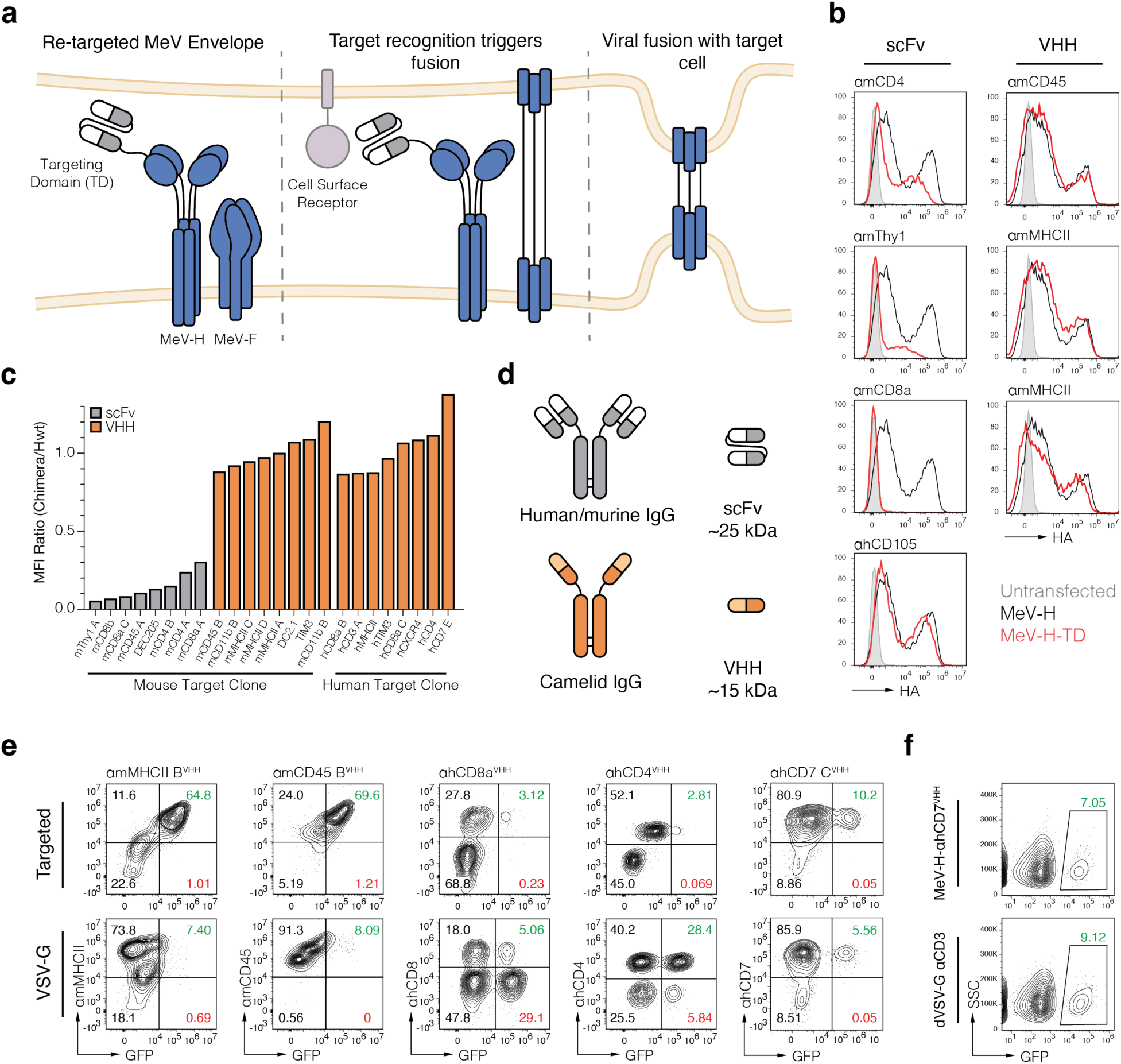
Camelid-derived VHH are a superior targeting domain for MeV-based lentiviral particle retargeting. (A) The hemagglutinin protein of MeV governs target recognition to trigger membrane fusion by proximity;, by adding an exogenous targeting domain the envelope can be re-targeted. (B) Surface expression flow cytometry staining for the MeV-H protein with scFvs and VHHs targeting a variety of relevant murine targets compared to the WT MeV-H protein. (C) MFI ratio of MeV-H-TD vs wt MeV-H for a selection of screened scFvs and VHH. (D) Human antibodies are larger than those from camelids, which lack the light chain and consist of a single heavy domain (VHH). (E) Flow cytometry of primary activated murine splenocytes or human PBMCs transduced with traditional VSVg pseudotyped LV carrying a GFP transgene or VHH re-targeted MeV across several targets. (F) Comparison of anti hCD7 MeV-H-VHH targeted LVs (top) to dVSVG pseudotyped LV with a CD3/28 targeting moiety (bottom).

Engineered MeV pseudotypes for *in vivo* CAR T therapy have been limited by several significant hurdles. First, fusion of scFvs to MeV-H can lead to intermolecular VH-VL pairing and reduced stability, resulting in poor functional titer and requiring screening of several scFv sequences to identify effective targeting domains^25^. Second, patients vaccinated against measles have neutralizing antibodies directed towards MeV-H and MeV-F^26,27^. Here we describe two technical advances to address these challenges: First, we find nanobodies (VHHs) are more successful targeting domains for MeV-H fusions and mediate efficient viral entry. Second, we demonstrate that the ectodomains from the H proteins of related morbilliviruses can be used to create chimeric Env proteins that maintain fusogenic ability, while avoiding humoral immunity, and can be successfully retargeted with VHH to pseudotype LVs. We show that retargeted LVs pseudotyped with a chimeric measles virus/Dolphin Morbillivirus (DMV) Env displaying a anti-hCD7 VHH can evade anti-MeV-H neutralizing antibodies and selectively transduce human T cells *in vitro* and *in vivo* to create CAR T cells. The described methods are an advance towards the generation of efficient and specific off-the-shelf CAR T therapies.

## Results

### Camelid-derived nanobodies are superior targeting domains for measles-virus pseudotyped lentiviral transduction

With the goal of creating targeted LVs for CAR T therapy, we initially designed MeV-H-scFv fusions targeting lineage-defining cell markers for T cells and other immune cells for proof of concept *in vivo* experiments. We obtained commercially available hybridomas with specificity for T cell lineage receptors (CD4, CD8a and Thy1) and sequenced the immunoglobulin complementarity determining region (CDR) sequences to create scFvs (**Extended Data Table 1**)^28^. scFvs were cloned into expression vectors such that the scFv sequence was fused to the extracellular domain of wild-type MeV-H (**Figure 1A, left**). We generated LVs with a GFP transgene pseudotyped with VSV-G or retargeted MeV-H-scFv fusions and evaluated transduction in unmodified, receptor-negative murine MC38 colon cancer cells, or MC38 cells engineered to overexpress target matched receptors (Thy1.2, CD4, or CD8a). As a positive control for re-targeting, we generated LV pseudotyped with a human CD105-targeted MeV envelope previously described (MeV-H-αhCD105^scFv^), with a corresponding hCD105 overexpressing MC38 cell line^17^. MeV does not bind mouse NECT4 or Slamf1 naturally, so we did not receptor-inactivate MeV-H enabling us to utilize MC38 cells overexpressing human NECT4 as a positive control. WT MeV pseudotyped LV provided moderate transduction of MC38 cells expressing the native receptor NECTIN4 (**Extended Data Figure 1A**).

All LVs pseudotyped with in-house MeV-H-scFvs demonstrated minimal or no transduction of cognate receptor expressing MC38 cells expressing, while the published MeV-H-hCD105^scFv^ did result in some measurable transduction (**Extended Data Figure 1A**). Most MeV-H-scFvs were able to transduce the native NECTIN4 indicating the engineered envelopes were still fusion competent. We concentrated LVs pseudotyped with our best performing MeV-H-αmThy1.2A^scFv^ which provided a slight increase in transduction of mThy1.2 MC38 cells (**Extended Data Figure 1B**). As overexpression cell lines do not strictly recapitulate the native protein, we attempted transduction of primary mouse splenocytes with endogenous Thy1.2 expression with lentivirus pseudotyped with the MeV-H-αmThy1.2A^scFv^ and did observe specific transduction (**Extended Data Figure 1C**), however the signal was unstable over the subsequent days. Our results suggest that improper receptor expression, and function of the retargeted envelope, do not explain the low transduction rates we observed using MeV-H-scFv retargeted lentivirus.

We hypothesized the poor transduction efficiencies could be due to low surface expression of the MeV-H-scFv in the HEK293T packaging cells as LV pseudotyping relies on abundant surface expression of viral glycoproteins on the packaging cell line for incorporation into virus particles. We transfected HEK293T cells with HA-tagged MeV-H-scFv expression vectors targeting CD4, Thy1, and CD8a, and compared cell surface expression of the fusions to surface expression of WT MeV-H (**Figure 1B**). Surface expression of MeV-H-scFv fusions was consistently lower than MeV-H alone or the previously reported MeV-H-αhCD105^scFv^ (**Figure 1C, Extended Data Figure 2A**)^17^. Moreover, the level of surface expression across HA-tagged MeV-H fusions correlated with transduction rates of hNECT4 MC38 cells (**Extended Data Figure 2B**), suggesting that viral retargeting was impeded by poor surface expression of the engineered envelope proteins.

scFvs are prone to aggregation due to the hydrophobic domains of VH and VL which facilitate their association in natural antibody tertiary structure^29^. However, in the context of an scFv, these regions can facilitate self-aggregation and mispairing^30^. To address whether inherent stability of scFvs impacted surface expression we purified His-tagged soluble scFvs from HEK293 cells. We observed scFvs that exhibited higher functional titer and/or surface expression also produced higher yields by SDS-PAGE gels (**Extended Data Figure 2C**), suggesting that intrinsic properties associated with scFv expression and/or stability may impact their potential as targeting domains. However, while the Thy1.2A^scFv^ produced purified yields comparable to hCD105^scFv^, the MeV-H-αThy1.2A^scFv^ fusion exhibited low surface expression and functional titer compared to MeV-H-αhCD105^scFv^, indicating additional factors contribute to impaired MeV-H-scFv expression. All purified 6His-tagged scFvs bound their cognate targets on engineered MC38s overexpressing cells *in vitro* (**Extended Data Figure 2D**), leading us to conclude the fusion of the scFv to MeV-H was the driver of low transduction rather than the binding ability of an scFv. To determine whether this phenomenon was generalizable, we evaluated a wider selection of scFvs targeting immune lineage markers including CD45, Thy1, CD4, CD8a, and CD8b and compared surface expression to WT MeV-H (**Figure 1C**, grey bars; **Extended Data Table 1**). Across all scFv sequences tested, we were unable to obtain surface expression comparable to WT MeV-H or the hCD105 scFv fusion (**Figure 1B**, left column; **Figure 1C**, gray bars). Since MeV-H assembles into tetrameric complexes (**Figure 1A**)^18^ and the hydrophobic regions of the heavy and light antibody chains within scFvs are prone to aggregation, we hypothesized that the close proximity of multiple scFvs was enhancing aggregation and reducing surface expression (**Extended Data Figure 2E**)^30,31^. As it has been reported that the success of retargeted LVs is highly dependent on the scFv used^25^, we sought alternative targeting domains.

Camelid-derived single-domain antibodies (nanobodies or VHHs) are smaller and more soluble than scFvs as they lack a light chain and consequently also lack the hydrophobic domains required for heavy and light chain assembly in human and murine antibodies (**Figure 1D**). We identified published VHHs targeting murine and human immune lineage markers (**Extended Data Table 1**) and cloned a subset into HA-tagged MeV-H expression vectors, then evaluated their surface expression in 293T producer cells. We found much higher surface expression of MeV-H-VHH on the producer cell surface compared to MeV-H-scFvs, at levels comparable to wild-type MeV-H for almost all VHH clones surveyed (**Figure 1B**, right column; **Figure 1C**, orange bars; **Extended Data Figure 2F**). LVs pseudotyped with MeV-H-VHH fusions targeting murine MHC class II or CD45 were generated and applied *in vitro* to activated mouse splenocytes. When compared to transduction using VSV-G-pseudotyped LV, the MeV-H-VHH constructs drove comparable or improved transduction efficiencies with increased target cell selectivity (**Figure 1E).** We then generated similar LVs targeting human CD4, CD8a, or CD7, and applied them to activated human PBMCs which resulted in similar high selectivity when compared to VSV-G, however less efficient transduction. These data suggest that VHH retargeting domains showed improved surface expression and functional titer over scFvs for MeV-H pseudotyped LV.

Recently, several methods of LV targeting have exploited a receptor blind vesicular stomatitis virus glycoprotein (VSV-G) that maintains fusogenic function, referred to as dead VSV-G (dVSV-G) for retargeting. When dVSV-G is expressed in combination with a separate surface protein for receptor targeting, specific delivery can be achieved ^32–35^. To benchmark our method to these approaches, we compared transduction of activated primary human T cells with LV pseudotyped with MeV-H-VHH targeting CD7 or LV pseudotyped with dVSV-G and the costimulatory pair of a membrane-bound scFv targeting CD3 and human CD80, and observed similar efficiencies (**Figure 1F**)^33^.

However, the use of dVSV-G may suffer from a decrease in specificity due to the requirement of low pH via endocytosis to trigger membrane fusion^36^, as non-specific interactions can result in endocytosis. Viral targeting using pH dependent Env has been reported to promote such off-target activity^37^. An endocytosis requirement also limits the potential receptors used for targeting. *In vitro* transduction of MC38 cells with LVs pseudotyped with dVSV-G demonstrated that dead VSV-G still drove detectable transduction activity (**Extended Data Figure 3A-B**), suggesting that targeted delivery strategies relying on dVSV-G pseudotyping maintain a risk of off-target transduction. For *in vivo* therapeutic applications, the MeV-H method represents a superior method for true selectivity and VHH a superior targeting domain ^8,38^.

### Alternative morbillivirus envelope pseudotyping enables evasion of measles-directed antibodies

Broad measles vaccination rates represent a significant potential obstacle to LV retargeting with MeV-H-VHH pseudotyping, as antibodies directed against MeV envelope proteins can neutralize infection, including transduction of MeV pseudotyped LVs ^39–41^. We collected sera from a cohort of 21 patients, aged 12 to 75 (median age of 52), that had balanced representation of males and females (**Extended Data Table 2**) and quantified the measles-specific IgG titers of donor samples by ELISA. As expected, the majority were seropositive aside from a single donor, confirming the broad presence of measles targeted antibodies (**Figure 2A**). A subset of donors representing high, medium and low titer was selected for further analysis (**Figure 2A**, colored bars). We developed a serum binding assay to measure serum immunoglobulin (IgG) binding to HEK293T cells expressing HA-tagged MeV-H and -F envelope proteins. HEK293T cells expressing HA-tagged MeV-H or MeV-F were incubated with heat-denatured patient serum before staining with anti-HA and anti-IgG antibodies and analyzing by flow cytometry (**Figure 2B**). Unsurprisingly serum-derived measles-directed antibodies bound surface-expressed MeV envelope glycoproteins in a serum dose- and anti-measles antibody titer-dependent manner (**Figure 2C**), potentially inhibiting the function of the envelope proteins.

**Figure 2:**
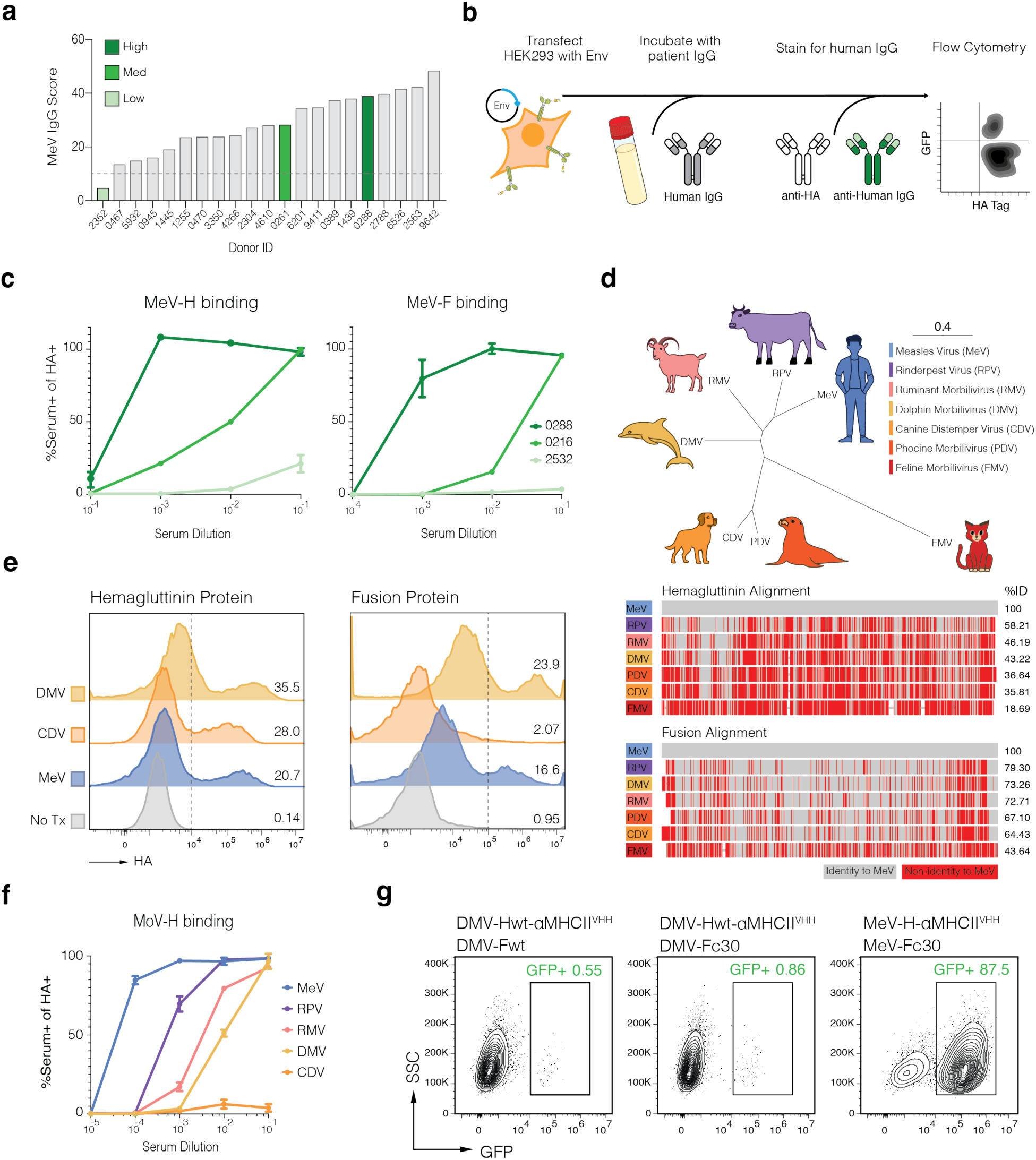
Pseudotyping with alternative morbillivirus glycoproteins can evade measles-directed antibodies. (A) Serum reactivity to measles from a cohort of self-reported MMR patients. A subset of low, medium and high reactivity (color bars) was selected for experiments. (B) Flow cytometry assay to determine MeV envelope binding of patient serum. Envelope expressing producer cells are incubated with patient serum, followed by staining with an anti-human IgG to determine serum Ab binding and an HA antibody to measure envelope expression. (C) Results from the assay in (B) for producer cells expressing MeV-H and MeV-F for the subset of patient samples. (D) Genetic distance for the MoV-H proteins in the morbillivirus family (top). Alignment of MoV envelopes to MeV shows low overall sequence similarity for the H protein. (E) Expression of DMV-αMHCII^VHH^, CDV-αMHCII^VHH^ and MeV-αMHCII^VHH^ envelope proteins in HEK293 producer cells. (F) Serum antibody binding to producer cells expressing a subset of MoV-H cells that demonstrated high surface expression. (G) LVs pseudotyped with MHCII-targeted DMV-H demonstrate minimal transduction. Through truncation of the F protein a slight increase was observed but significantly less than measles.

The mechanism for receptor recognition and envelope fusion is conserved among all morbilliviruses (MoV) and the greater paramyxovirus family^20^. However the surface glycoproteins have low overall sequence similarity, particularly in their extracellular domains, while still maintaining structural homology from AlphaFold3 predictions (**Figure 2D, Extended Data Figure 4A-B**)^20,42^. Measles-neutralizing antibodies primarily target the extracellular domains of H and F proteins, as such we hypothesized that alternative MoV glycoproteins could be similarly engineered for re-targeting but evade neutralization from preexisting immunity against MeV^43–45^, as has been accomplished with Nipah Virus for LVs^46^ and CDV for oncolytic virus therapy^27^, and modified for MeV ^41^. We expressed HA-tagged retargeted hemagglutinin proteins from several morbillivirus species and evaluated surface expression on HEK293T cells compared to a well-expressed MeV-H-αmMHCII^VHH^. Each retargeted MoV-H fusion protein was successfully expressed on the surface of HEK293T producer cells at levels comparable to MeV-H, except for Phocine mobillivirus (PDV)-H **(Figure 2E, Extended Data Figure 4C**). We attempted to measure wild-type unmodified PDV-H surface expression but were unable to detect any in HEK293T cells (data not shown). In a serum binding assay we observed higher binding of measles-directed antibodies to H proteins of Rinderpest (RPV) and Small ruminant virus (RMV), and reduced binding to the less homologous Canine distemper virus (CDV)- and Dolphin morbillivirus (DMV)-H proteins (**Figure 2F**). As prior work has demonstrated that measles neutralizing antibodies primarily target MeV-H^43–45^, we focused our efforts on optimizing CDV and DMV glycoproteins for further engineering and in vivo CAR delivery.

DMV-F exhibited high levels of surface expression on HEK293T cells while CDV-F was barely detectable (**Figure 2E**, right). Given the large reduction in anti-measles antibody binding to CDV-H, we attempted to optimize CDV-F expression. We replaced the unusually long signal peptide of CDV-F with shorter alternative signal peptides^47,48^ and tested previously reported truncations^49^, but were unable to drive higher CDV-F expression (**Extended Data Figure 5A**). DMV-H also exhibited resistance to measles-directed antibody binding and both DMV-H and DMV-F were well expressed in HEK293T, thus we opted to focus on DMV for LV pseudotyping and VHH retargeting. Previous studies found that functional pseudotyping of LVs with MoV required truncation of the cytoplasmic tail of MoV-F^15,50^. We designed DMV-H fused to a VHH targeting mouse MHC-II and co-expressed the construct with full-length or truncated DMV-F to generate LV and compare transduction of target cells to MeV-H-VHH. While we detected low transduction of MHCII+ A20 leukemia cells with DMV-pseudotyped LV, the functional titer was much lower than that of MeV pseudotypes and further engineering would be required to enhance transduction efficiency (**Figure 2G, Extended Data Figure 5B**).

### Chimerized DMV/MeV glycoproteins improve functional titer while maintaining immune evasion

The structural similarities between MoV glycoproteins have been leveraged to create functionally chimeric H hybrid proteins swapping the domains between MoV^51^. MoV-H proteins play a role in triggering membrane fusion through interactions with MoV-F, In addition to receptor recognition. Regions in the globular head (aa244-250) and in the stalk (aa84-105) of MeV-H have been shown to interact with MeV-F^52^. Functional hybrids of MeV/CDV glycoproteins have been created that preserve the key interactions between species-matched H and F domains have been successfully pseudotyped, in addition to resisting neutralization by measles-immune serum^51^. We hypothesized a similar strategy could produce chimeric proteins that preserve the functional MeV-H core while decreasing immune recognition with the ectodomains of DMV or CDV (**Figure 3A**). Chimeric MeV/DMV-H and MeV/CDV-H were generated with the HA tagged αMHCII^VHH^ and were well expressed on the surface of producer cells (**Extended Data Figure 6A)**. Furthermore, both H protein chimeras produced affected pseudotyped LVs with MeV-F, that demonstrated comparable if not superior functional titer when applied to A20 cells (**Figure 3B**).

**Figure 3:**
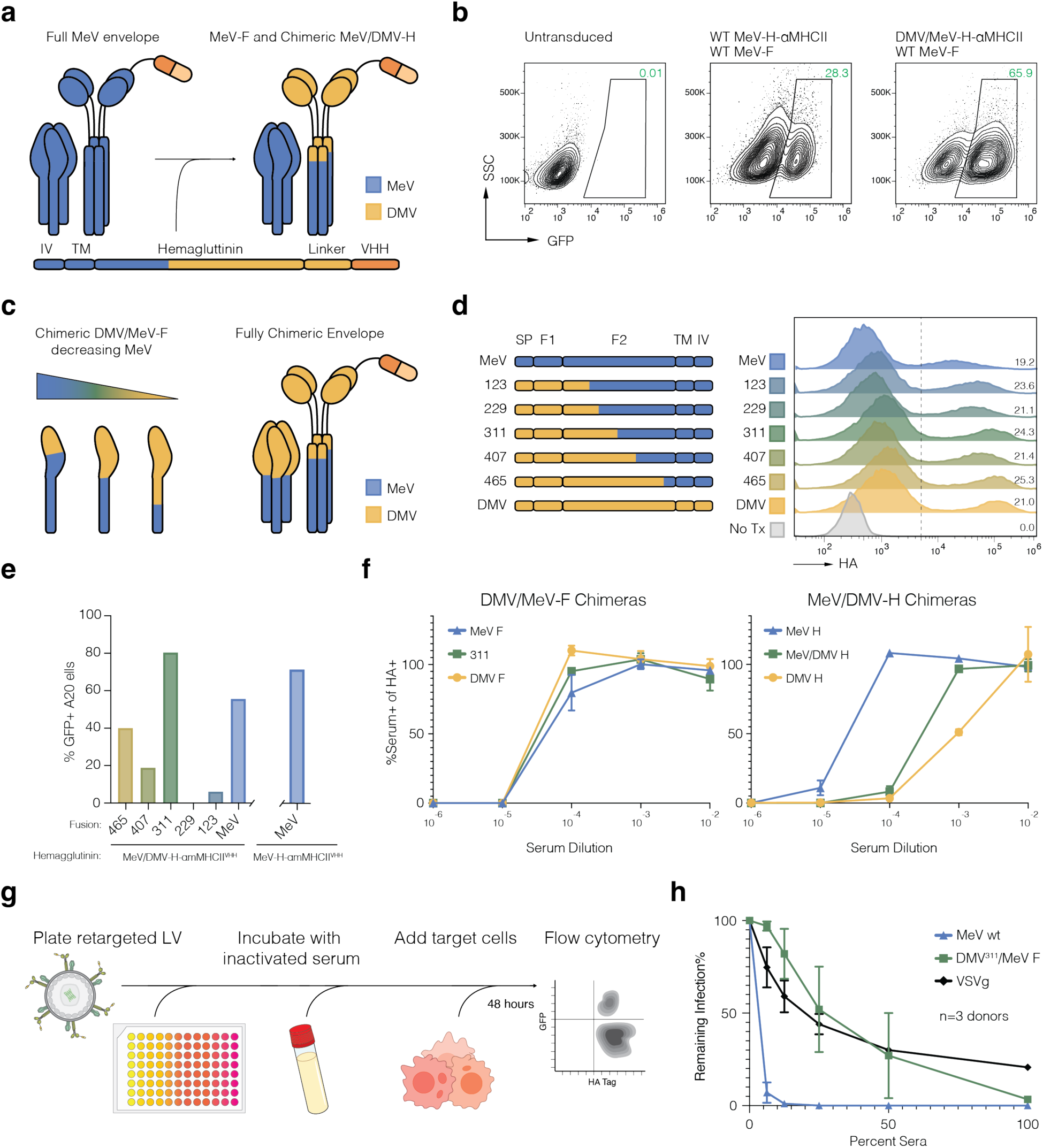
Lentivirus pseudotyped with MeV/DMV chimeric envelope proteins are not neutralized in measles-vaccinated sera. (A) Design of chimeric Hemmagluttinin protein. Residues 1-N are MeV while N-End are DMV. (B) Chimeric MeV/DMV-H produces infectious LVs with MeV-F comparable to that of standard MeV-H/F LVs. (C) Design of complementary chimeric fusion proteins. As the MoV-F proteins are type I transmembrane proteins and thus the signal peptide is that of the extravirion MoV. (D) Surface expression of tested chimeric MeV/DMV-F proteins. (E) Transduction of chimeric MeV/DMV-F protein LVs with Chimeric MeV/DMV-H protein (left) and standard MeV-H (right). Chimeric MeV/DMV-H and DMV^311^/Mev-F produce titers comparable to that of fully MeV pseudotyped LVs. (F) Patient serum binding to chimeric envelope proteins utilizing the assay described in 2B. (G) Flow cytometry assay to determine neutralization of viral particle infectivity. Re-targeted LV is plated in a series of dilutions of patient donor serum. Target cells are then added, and the plate is incubated for 48 hours before analysis by flow cytometry. (H) Quantification of neutralized infection comparing LV pseudotyped with full MeV (MeV-H-αMHCII^VHH^ + MeV-F), chimeric MeV/DMV (MeV/DMV-H-αMHCII^VHH^; DMV^3^^11^/Mev-F), or VSV-G envelopes (n=3, patient IDs 6526, 6201, 9411).

Neutralizing responses to measles are predominantly directed toward MeV-H with some reports suggesting MeV-F is also targeted^44,45^. We detected some serum antibody binding to most MoV-Fs with the functional titer of wild-type DMV-pseudotyped lentivirus being too low to accurately test neutralization (**Figure 2G**). We reasoned that by generating chimeric DMV/MeV-F proteins, we could further reduce potential neutralization while maintaining high viral titers (**Figure 3C**). MoV-F proteins are type I transmembrane proteins so to replace the ectodomain requires substitution of the N terminus and signal peptide. We designed and confirmed surface expression of several chimeras DMV and MeV Fusion constructs with increasing contributions of DMV-derived ectodomain (**Figure 3D**). We generated LVs pseudotyped with each chimera construct paired with MeV/DMV-H-αMHCII^VHH^ or MeV-H-αMHCII^VHH^ and assessed transduction of A20 cells. Only the DMV^311^/MeV-F construct exhibited comparable transduction efficiency of target cells compared to MeV-pseudotyped LV, confirming the chimeric proteins retained critical interactions and fusogenic capacity (**Figure 3E**). Parallel attempts to functionalize chimeric CDV/MeV-F, including the signal peptide modifications and additional rational engineering strategies, failed to generate well-expressed or functional CDV/MeV-F chimeras **(Extended Data Figure 6B-C)**.

The MeV-DMV-H chimera had a large reduction in serum binding while theDMV^311^/MeV-F demonstrated no change (**Figure 3F**). However not all antibody responses are neutralizing, so we performed serum neutralization assays (as previously described) with the MeV/DMV-H-αMHCII^VHH^;DMV^311^/MeV-F-pseudotyped LVs applied to A20 cells and compared to VSV-G-LVs and MeV-αMHCII^VHH^-LVs (**Figure 3G**). Across serum three donors we observed a significant increase in average transduction of the dual-chimeric approach compared to MeV-αMHCII^VHH^-LV, and rates comparable to the VSV-G-LV condition (**Figure 3H)**. Despite reported low levels of VSV-G-specific antibodies in humans^53^, we observed a dose-dependent decrease in transduction with the VSV-G-LV, suggesting non-specific factors in human serum limit *in vitro* transduction^54^ (**Figure 3H**, black). Collectively, this data demonstrates chimerization successfully evaded measles-specific antibody neutralization. With a successful fully chimeric immune resistance and selective re-targeted pseudotyping strategy in hand, we set out to evaluate feasibility of *in vivo* targeting for CAR T generation using this approach.

### Retargeted LV enables CAR T generation and killing in mixed *in vitro* cultures

*In vivo* CAR T cell generation poses a challenge as resting T cells are extremely resistant to lentiviral transduction^55–57^. During conventional *ex vivo* CAR T cell manufacturing, T cells are activated and expanded *in vitro*, which increases expression of LDLR, the receptor for VSV-G, increasing the transducability of the cells^58,59^. T cell activation also induces effector differentiation and increases cytotoxic functions, enhancing the therapeutic potential of the CAR T cells when reintroduced to the patient^60^. Activation of T cells in both research and clinical contexts is typically accomplished with reagents derived from antibodies targeting CD3 and CD28 to deliver TCR stimulation and costimulation, respectively, as well as supplemental cytokines^61^. For *in vivo* generation of CAR T we set out to incorporate co-activation signals to drive T cell activation and expansion in our re-targeted LVs. We engineered producer HEK293T cells to express both a membrane-tethered scFv targeting CD3 (αCD3) and human CD80, the natural ligand for the T cell costimulatory receptor CD28. These cell surface molecules would then be incorporated into virions, as has been previously described, and activate target T cells^33^. As CD19 CAR T products composed of mixed CD8+ and CD4+ cells are associated with superior and more durable responses^38,62^, we elected to use a pan T cell targeting construct MeV/DMV-H-αCD7^VHH^ for CAR delivery and included a GFP reporter.

We first tested the activation of primary T cells co-cultured with unmodified or αCD3/CD80 HEK293T cells in vitro, and observed significant upregulation of T cell activation markers CD25 and CD69 in the αCD3/CD80 condition (**Extended Data Figure 7**). We generated CAR-GFP LV in unmodified HEK293T or αCD3/CD80 HEK293T cell lines, with either VSV-G and MeV/DMV-H-αCD7^VHH^ pseudotypes and tested their transduction and activation of primary T cells *in vitro* (**Figure 4A**). We tested both unstimulated human primary T cells and pre-stimmulated T cells using standard αCD3/αCD28 bead-based activation, and measured activation by expression of CD69 and transduction by GFP fluorescence. As expected, T cells preactivated with beads showed high levels of activation markers and GFP expression regardless of packaging cell line and pseudotype, though LV carrying αCD3/CD80 drove further increases in activation and GFP+ cells (**Figure 4B**, top). Transduction of non-activated T cells with αCD3/CD80 HEK293 produced LVs induced a marked increase in CD69 and GFP expression compared to standard HEK293 (**Figure 4B**, bottom). These data support that LVs enhanced with αCD3/CD80 can successfully drive activation of target T cells and enhance transduction efficiency.

**Figure 4:**
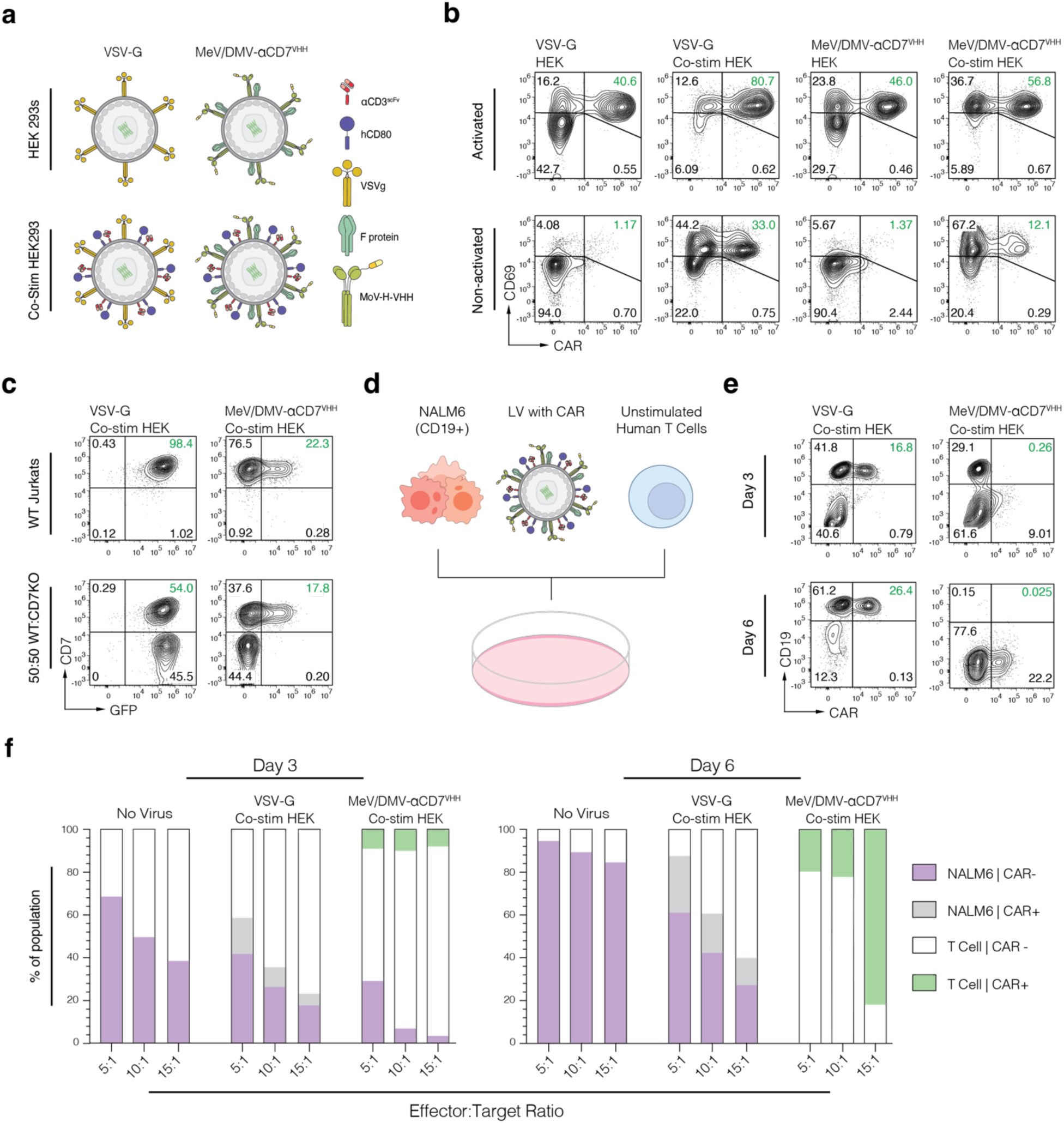
Retargeted and stimulatory LV enables CAR T generation and killing in mixed in vitro cultures. (A) Stimulatory functionalization of pseudotyped lentiviral vectors with chimeric MoV targeted envelopes. Co-stim HEK293s have been engineered to express a surface anchored αCD3 scFv and CD80 to enable activation of T cells. (B) Comparison of standard and co-stim functionalized VSV-G and chimeric MeV/DMV αCD7 pseudotypes on pre-activated and unactivated human T cells. Co-stim LVs successfully activate unactivated T cells as measured by CD69 expression and result in greater transduction. (C) Specificity analysis of finalized TaSEL vectors on CD7 knock-out Jurkat cells. (D) *In vitro* CAR-T generation and cancer killing assay. Unactivated T cells (effector) and NALM6 (target) cells are mixed at varying ratios with LVs containing a CAR-GFP transgene. (E) Example data from (D) comparing VSVG and TaSEL pseudotypes. (F) Quantification at day 3 and 6 post mix of D with VSV-G and TaSEL pseudotypes at varying effector to target ratios.

To confirm that αCD3/CD80 enhancement did not impact the specificity of the chimeric retargeting approach we used MeV/DMV-H-αCD7^VHH^ LVs to transduce a mixed population of wildtype (CD7+) and CRISPR knock out (CD7-) Jurkat T cells. MeV/DMV-H-αCD7^VHH^ LV only transduced CD7+ cells, with minimal GFP signal detected in Jurkats lacking expression of CD7 while VSV-G-LVs transduced both populations equally (**Figure 4C**). In an *in vitro* co-culture assay we mixed unstimulated human primary T cells with CD19+ NALM6 B cell lymphoma cells at varying effector to target ratios (**Figure 4D**), then we added αCD3/CD80 enhanced LVs pseudotyped with VSV-G or MeV/DMV-H-αCD7^VHH^ and monitored T cell activation, CAR+, and tumor killing by flow cytometry. By Day 3 a significant proportion of the CD19+ NALM6 cancer cells were CAR+ in the VSV-G-LV condition, while the MeV/DMV-H-αCD7^VHH^ LV only transduced T cells (**Figure 4E**). This effect grew more pronounced by day 6, which was also marked by a relative expansion of CAR+ T cells and a reduction in total NALM6 cancer cells in the MeV/DMV-H-αCD7^VHH^ condition (**Figure 4E**). Indeed, as the ratio of effector to target cells increased in MeV/DMV-H-αCD7^VHH^ wells, we observed a significant reduction in total recovered NALM6 cells at the earlier time point, and all conditions eliminated target CD19+ cells by day 6 (**Figure 4F**). In the VSV-G-LV condition by contrast only CAR+ cancer cells were detected in co-culture, and we were unable to detect CAR T generation or cancer-directed cytotoxicity. Surface enhanced and retargeted LV can specifically and successfully generate CAR T cells in a heterogeneous context without the need for additional T cell stimulation.

### *In Vivo* administration of retargeted LV generates effective CAR T response

With successful *in vitro* experiments we moved on to *in vivo* models using an LV composed of MeV/DMV-H-αCD7^VHH^, DMV^311^/MeV-F, and αCD3/CD80, now referred to as Targeted and Stimulatory Engineered Lentivirus (TaSEL) (**Figure 4A**, bottom right). We used a xenograft mouse model to evaluate the capacity of TaSEL vectors to generate human CAR T cells *in vivo* and compared anti-tumor efficacy to conventional *ex vivo* generated CAR T cells. Luciferase-expressing NAML6 B cell leukemia cells were injected into immunodeficient NOD.Cg-*Prkdc^scid^Il2rg^tm1Wjl^*/SzJ (NSG) mice at day -8 to establish B-ALL tumors. On day -1, a subset of mice then received intravenous (i.v.) injections of human T cells, of which one cohort also received a single injection of TaSEL delivering αCD19 CAR-GFP and the other received no virus. As a control for the expected efficacy of CAR T cell therapy in this model, we included a group of tumor-bearing mice treated with a dose of conventional αCD19 CAR-GFP T cells made ex vivo (**Figure 5A)**. Throughout the experiment, mice were imaged twice weekly to monitor tumor burden, and peripheral blood was collected weekly via submandibular bleeds to assess CAR T production.

**Figure 5:**
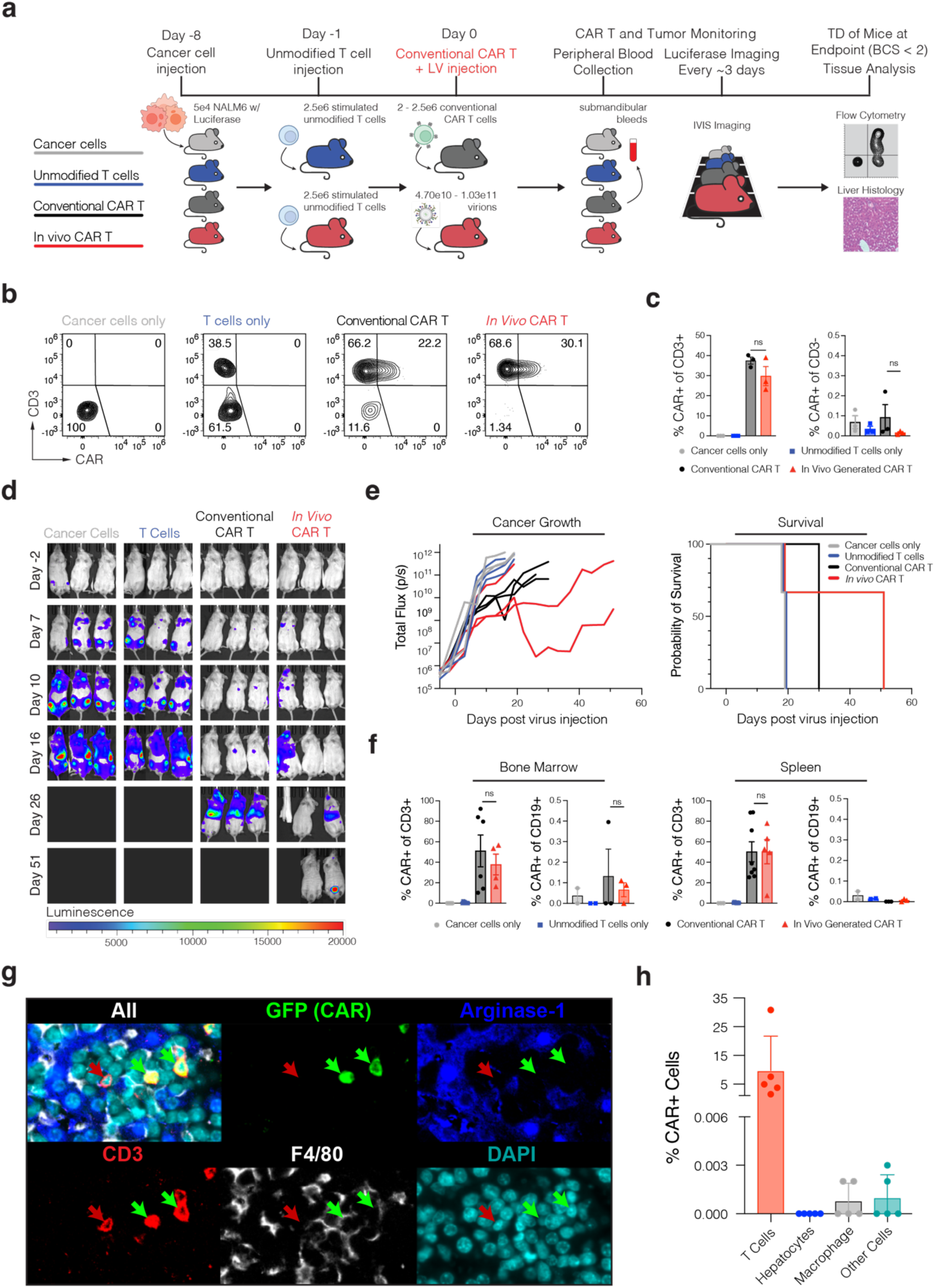
*In vivo* administration of TaSEL generates CAR T cells and controls NALM6 tumors. (A) Experimental scheme for *in vivo* CAR T cell generation and evaluation. Cohorts of mice all received NALM6 leukemia cells expressing luciferase. Control mice either received no T cells or unmodified human pan T cells. Treatment mice received unmodified T cells chased by TaSEL the following day (*In Vivo* CAR) or *ex vivo* generated CAR T cells (Conventional CAR). Analysis consisted of luciferase imaging for tumor quantification, per. (B) CAR T cell content of peripheral blood collected on D6 post virus injection via submandibular bleeds. X-axis is CAR+ (FITC), y-axis is CD3+ (APC) (C) Quantification of CAR+ frequencies in target T cells (CD3+) and off-target cells (CD3-). (D) Representative IVIS imaging at each indicated timepoint following TaSEL administration. Mice not shown were sacrificed according to Body Condition Score guidelines and tissues were collected and analyzed. Bioluminescence overlays represent current tumor burden for a given mouse (E) Quantification of individual mouse tumor burden in radiance (photons/second) over time and probability of survival for each mouse group over time. (F) Quantification of transduction frequencies of cell types of interest in spleen and bone marrow. CD19+ = NALM6 cells, CD3+ = T cells, CAR+ = CAR T cells. Data represents mean ± SEM (all groups n = 3, used one-way ANOVA) (G) Multiplex immunofluorescence staining of liver sections from mice with *In Vivo* CAR. Large tissue sections were stained, imaged, and analyzed. Cell segmentation, and classification was performed followed by assessment of GFP staining intensity. Breakdown of GFP-positive cells (CAR+) within each cell-type is shown. Groups were compared with a Mann-Whitney statistical test.

At day 6 post injection the transferred human T cells had engrafted into mice and TaSEL had successfully transduced target T cells to generate CAR+ T cells *in vivo* at frequencies comparable to that of conventional CAR T cell transfer (**Figure 5B**). Additionally, no GFP fluorescence was detected in off-target cell populations by flow, indicating highly selective transduction by TaSEL (**Figure 5C).** To understand the functionality of the CAR T response, we also monitored tumor burden with non-invasive luciferase imaging over time and observed that untreated animals (NALM6 only, T cells only) rapidly succumbed to disease (**Figure 5D-E**). Mice that had received conventional CAR T cells showed modest control and slowing of tumor growth, although none survived beyond 30 days. In the cohort of mice that received T cells and a single dose of TaSEL, two of three mice showed striking and prolonged control of tumor growth and survived long enough that they began showing signs of xenographic graft-versus-host disease (x-GVHD), a common limiting factor in humanized mouse models (**Figure 5D-E**).

We analyzed tissue collected in sacrificed animals for evidence of on- and off-target infection events. Flow cytometry of bone marrow and spleen cells revealed comparable engrafted GFP+ CAR T cell frequencies at endpoint in animals that received either conventional CAR T or *in vivo* administration of TaSEL (**Figure 5F**). Analysis of non-T cell populations (CD3-) in collected blood at endpoint failed to detect off-target transduction in TaSEL-treated mice (**Extended Data Table 3**). CAR-GFP signal was also undetected in off-target cells in the bone marrow or spleen, corroborating our earlier findings in peripheral blood (**Figure 5F**). To further interrogate off-target effects we performed multiplexed immunofluorescence analysis of liver tissue sections to determine whether LV-delivered GFP could be detected in cells other than T cells (αCD3) or Kupffer cells/macrophages (αF4/80), which are known to sequester lentiviral particles in the liver^63,64^(**Figure 5G, Extended Data Figure 8**). The GFP+(CAR+) cells identified in these sections were over 99.7% CD3-positive. Across multiple broad tissue sections, no GFP-positive hepatocytes were observed. Rare off-target (CD3-negative) GFP-positive cells were identified at a frequency less than 1/6,700th of that observed within the CD3-positive T cell compartment. These events occurred at such a low frequency that we cannot exclude them as being attributed to fluorescent staining artifacts which are prone to occur in large-tissue multiplex immunofluorescence studies such as this. Thus, *in vivo* delivery of TaSEL can selectively and efficiently generate CAR T with comparable or improved anti-tumor function, relative to conventional manufactured CAR T.

## Discussion

Here we demonstrate that TaSEL is a significant advance for *in vivo* CAR T generation that drives efficient and specific cell type transduction and evades broad vaccine-induced immunity and neutralization. To optimize this system for the *in vivo* delivery of chimeric antigen receptors to T cells, we made two major advances: First, we show that the use of camelid-derived heavy chain-only (VHH) antibodies dramatically improves the function of morbillivirus envelope glycoprotein pseudotyping as a strategy for redirecting virus specificity. Second, we generate a chimeric envelope where the extracellular region is derived from dolphin morbillivirus (DMV), which has low amino acid identity with measles virus and substantially evades neutralization from anti-measles antibodies. We further functionalized the retargeted viral particles by incorporating αCD3 and CD80 to activate T cells upon viral encounter to ensure T cells are competent for transduction. These strategies combined enable the selective and efficient delivery of CARs to T cells in an in vivo model, which shows efficacy that is at least comparable to conventional *ex vivo* generated CAR T cells.

Lentivirus can be pseudotyped with viral envelope proteins from several families of viruses^4–6,32^, and receptor engineering to accomplish cell type-selective retargeting has been demonstrated using envelope glycoproteins from several viruses including Sindbis virus envelope ^65–68^ and a receptor binding mutated “blind” version of VSV-G^33–35^. However, these exhibit mediocre specificity, likely because membrane fusion requires a pH-dependent trigger from endocytosis^36^, which can also result from nonspecific interactions with the host cells, independent of receptor binding^37^. The technologies that have proven to be the most selective are those that employ envelope glycoproteins from the paramyxovirus family, including Nipah virus (NiV) and measles virus^37,46^. Paramyxovirus envelope glycoproteins are multiprotein complexes where the receptor recognition performed the tetrametic H protein, and fusion by the trimeric F protein. Fusion of the envelope complex is triggered by receptor recognition at the cell surface, and is independent of pH or endocytosis^20^. This is an optimal mechanism for receptor retargeting as it eliminates unwanted off-target transduction from the inevitable envelope-independent endocytosis of viral particles^37^.

While the envelopes of MeV and NiV are amenable to engineering both viruses are human pathogens. Immunity to MeV is widespread in the human population due to worldwide vaccination efforts. Neutralizing antibodies induced by vaccination have been found directed against the envelope H protein and possibly against the F protein of measles^43,69^. While NiV infection is currently rare and immunity in humans not widespread, NiV has a documented history of outbreaks^70^. NiV infection is life-threatening and the World Health Organization has identified Nipah virus as a potential cause of future epidemics and vaccine development efforts are underway^70,71^, which would preclude the use of Nipah virus envelope for therapeutic lentiviral engineering.

In this work, we have demonstrated that chimeric envelope proteins between different morbillivirus’ can pseudotype lentiviral particles and be engineered for cell specific transduction. These viruses are not known to infect humans so future widespread immunity against them is unlikely. Their envelope glycoproteins share significant structural homology with measles virus but have low amino acid identity, and thus the ability for anti-measles antibodies to neutralize transduction via these envelopes is limited. We show here that even in donors with a high anti-measles antibody titer, binding of s chimeric MeV/DMV pseudotyped lentivirus is reduced at least 100-fold and neutralization of transduction is equivalent to a VSV-G control. As the conserved epitopes for anti-measles neutralizing antibody binding have been well-studied^69^, future work to alter amino acids in these key areas could even further reduce antibody binding. Thus, our method of using dolphin morbillivirus envelope enables specific and neutralization-resistant lentiviral pseudotyping that is not threatened by future public health efforts.

The use of camelid-derived single-domain antibodies as the targeting domain is a key advance that enables the high efficiency we observe with TaSEL. Previous work has mainly implemented single chain variable fragment (scFv) antibodies to redirect measles or NiV pseudotyped lentiviral vectors. While scFvs have produced successful re-targeted LVs, optimization is often required with testing many potential scFvs, suggesting that scFvs are generally inefficient for re-targeting^8,25^. We hypothesize that the mechanism for inefficiency is self-aggregation and mispairing via the hydrophobic regions that natively facilitate Vh and Vl binding^30^.

The intrinsic tendency for self-aggregation of scFvs is well documented ^72^ and could be exacerbated by the tetrameric H protein complex on the surface leading to poor stability^18^. We observed universally low expression of our designed MeV-H-scFv fusion, supporting the potential aggregation. Other binding domains such as DARPins have been used^24^, although no systematic effort has been reported and the success rate for DARPins is unclear. Furthermore DARPins are used infrequently and require screening large display libraries, limiting potential candidates. VHHs on the other hand are increasingly popular and as such many more binders are available. Beyond this, the ability to immunize camelids or the recently developed nanomouse against any target of interest makes the technology more accessible ^73^.

Finally, we demonstrate that our DMV-H protein retargeting system can be combined with lentiviral surface engineering to functionalize viral particles. By engineering viral packaging cells to overexpress an αCD3 scFv and CD80, these molecules can be incorporated into viral particles to enable activation of T cells on contact^33^. Efficient LV transduction of T cells requires T cell activation and proliferation, likely due to retroviral transduction generally being inefficient in non-dividing cells and downregulation of viral restriction factors^74,75^. The majority of circulating T cells in human blood and secondary lymphoid organs have either an effector memory (EM) or central memory (CM) phenotype and are not activated and rapidly proliferating. The inability to efficiently transduce these populations would severely restrict the “accessible” population of T cells in a patient, potentially limiting the efficiency of the product. Additionally, recent evidence suggests that CAR T cells generated from CM T cells have superior efficacy and persistence^76^, thus transducing these populations of T cells *in vivo* would be a priority for an off-the-shelf CAR T cell product. Surface functionalization is a prime method for further engineering of LVs.

TaSEL provides a significant advantage compared to other current methods of LV re-targeting for *in vivo* use. There are a few limitations of TaSEL, namely in production and further immune reduction. Generally, the MoV based pseudotyping methods result in lower functional titers than VSV-G. The low titers require a large amount of LV to be produced for *in vivo* experiments and they are difficult to concentrate. Optimizations for production and increases in functional titer would be beneficial to creating an off-the-shelf therapy. For reduced immune clearance we focused on DMV for chimerization, however CDV likely provides a further advantage. While we were unable to generate a usable CDV-F construct, the potential for a further reduction in avoidance on neutralizing antibodies is possible. Importantly, we still demonstrate a significant reduction *in vitro* where neutralization is easy to observe, the *in vivo* context may not require as significant reduction to achieve measurable effects.

In summary, our TaSEL technology overcomes a series of unique challenges for *in vivo* T cell-focused gene therapy and may represent a path forward to generate off-the-shelf T cell therapeutics for patients. The system could be used for the delivery of any therapeutic payload that can be packaged into a lentiviral vector and applied to any therapeutic area where engineered T cells are currently being evaluated, including hematologic malignancy, solid tumor malignancy, and autoimmune disease. Given the high unmet need in these therapeutic areas and the ongoing challenges of manufacturing engineered autologous T cell products for all patients who could benefit from them, off-the-shelf technologies such as TaSEL should be a major focus for the biotechnology industry.

## Acknowledgements

We thank Branka Horvat and the Measles Virus Biobank for the generous gift of the CL55 and Y503 antibodies, INSERM, France, Measles virus Biobank https://ciri.ens-lyon.fr/teams/IbIV/measles-biobank. We thank Nathan Pirete and the Broad Institute Flow Cytometry Core for training and assistance in flow cytometry experiments and the Broad Institute vivarium staff for the assistance with mouse studies. We would like to thank Professor Fritz Melcher from the German Rheumatology Research Center Leibniz-Institute for providing the FGK4.5 hybridoma, Dr. Hasaya Akiba for providing the RMT3-23 hybridoma, and Dr. Yoji Murata for providing the MY-1 and Miap301 hybridomas.

## Author information

These authors contributed equally: Kepler S. Mears, Kyrellos Ibrahim, and Peter M. Allen These authors contributed equally: Jordan M. Chinai, Omar I. Avila and Audrey J. Muscato

## Ethics Declarations and competing interests

This manuscript was supported in part by funding from Calico Life Sciences, LLC. R.T.M., K.B.Y., K.S.M., K.I., and P.M.A. are authors on a patent related to the technology described in this manuscript. R.T.M. has received speaking or consulting fees from Bristol Myers Squibb, Gilead Sciences, Kumquat Biosciences, Immunai Therapeutics, and BioNTech and has equity ownership in OncoRev, LLC and Jumble Therapeutics.

## Author contributions

Conceptualization, K.S.M., K.B.Y., and R.T.M.; formal analysis, K.S.M., P.M.A, K.I., J.M.C., O.I.A., A.J.M, S.K.L-R., A.R. and C.C.; investigation, K.S.M., P.M.A, K.I., J.M.C., O.I.A., A.J.M, S.K.L-R., A.R., N.H.K. and C.C.; resources, K.S.M., P.M.A, K.I., J.M.C., O.I.A., A.J.M, S.K.L-R., A.R., N.H.K. and C.C.; writing – original draft, K.S.M., P.M.A, K.I., J.M.C., K.B.Y., and R.T.M.; writing – review and editing, K.S.M., P.M.A, K.I., K.B.Y., J.M.C., and R.T.M.; visualization K.S.M., P.M.A, K.I., J.M.C.; supervision, , K.S.M., K.B.Y., and R.T.M.

## Methods

### Mouse experiments

Immunodeficient NOD.Cg-*Prkdc^scid^Il2rg^tm1Wjl^*/SzJ (NSG) mice were raised in-house. NALM6 cells, human pan T cells, and TaSEL were all administered via tail vein injections using a rodent restrainer. Tumor cells were prepared by 3 washes in HBSS (Gibco/ThermoFisher 14065056) as follows: pellet by centrifugation at 350×g for 5 minutes, decant supernatant, then resuspend at 5e5 - 8e5 cells/mL. Tumor cells were counted with a Countess II (ThermoFisher) and diluted to a final concentration of 5e5 cells/mL HBSS. Mice were administered 100 µL (5e4 tumor cells) via tail vein injection 8 days before TaSEL injections. Human pan T cells were similarly prepared with 500×g centrifugation. T cells were counted with a Countess II and verified by hand with a hemacytometer, then diluted to a final concentration of 2.5e7 cells/mL. Mice were administered 100 µL (2.5e6 T cells) via tail vein injection 1 day before TaSEL injections. CAR T cells were generated as described for *In vitro CAR T cell activation and killing assay* and CAR expression rates were quantified by flow cytometry. Cell preparation was adjusted to administer 2.5e6 CAR-positive T cells per 100 µL injection. Finally, 100 µL TaSEL was administered on Day 0 via tail vein injection.

### Peripheral bleeds

Submandibular blood was collected by puncturing the vein slightly behind the mandible and below the ear canal with a 4mm animal lancet. Blood was collected in heparin tubes and mice were monitored for recovery. In the event a submandibular bleed could not be initiated, blood was collected from the tail vein using a lancet.

### IVIS imaging

Mice were injected with D-luciferin (PerkinElmer, XenoLight) at 150ug/g body weight, prepared following manufacturer’s instructions. After a 15 minute incubation, mice were anesthetized with 1% isoflurane. Mice were then transferred to a Perkin Elmer IVIS Spectrum CT and luciferase signals were obtained by bioluminescence imaging. Mice were imaged on average every 3 days during tumor challenge experiments.

### Endpoint

When endpoints were reached in accordance with Body Condition Scoring, mice were sacrificed CO_2_ and terminal bleeds. Blood, spleen, liver, and lung samples were collected for analysis. Single-cell suspensions of spleen were prepared by pressing the organ through 70 µm filters with the back of a 1 mL syringe. Splenocytes were isolated by filtering collected blood through a 70 μm filter, followed by lysis of erythrocytes (ACK Lysing Buffer, Gibco). Lung and liver samples were prepared as described in *Multiplex Immunofluorescence Staining, Imaging, and Analysis*.

#### Vector design

MoV envelope proteins were purchased as gene blocks from GenScript and expressed from a CMV promoter from plasmid pCG-HcΔ18 (addgene 84817). Lentiviral reporter transfer vectors were generated from pLX311-Cas9 (addgene 118018) with Cas9 swapped with GFP. CAR lentiviral transfer vectors were pSLCAR-CD19-BBx (addgene 135992).

Hybridomas were sequenced by GenScript to determine the VH and VL domains. Single chain variable fragments were designed by linking the VH and VL domains with a flexible linker (G_4_S)_3_ then codon optimized for human expression and gene blocks ordered from GenScript. VHH sequences were similarly codon optimized and ordered. MoV-H-scFv/VHH fusions were cloned into the pCG backbone.

Sequence for the co-stimulatory αCD3 (OKT3 clone)-P2A-hCD80 (UniProt sequence) were purchased as a gene block from GenScript and Gateway assembled into a lentiviral transgene vector pLEX-307 (addgene 41392) to be stably expressed from an EF1a promoter.

#### Lentiviral Preparations

For LVs retargeted with morbillivirus envelope proteins, LentiX-293T cells were transfected with morbillivirus H and F envelope plasmids, packaging plasmid psPAX2 (Addgene: 12260) and transfer plasmid at a 5:3:24:32 ratio, with 2𝜇g plasmids DNA transfected per 1e6 cell seeded. For VSV-G LV, Lenti-X 293T cells were transfected with VSV-G envelope (pMD2.G, Addgene:12259), psPAX2 and transfer plasmid at a ratio of 1:3:4. The media was changed 8 hours post transfection. Supernatant collected 48 hours post media change was purified by 0.45 um SFCA filter membrane. For in vitro assays, particles were concentrated 100-fold by incubation with Lenti-X™ Concentrator (Takara Cat. #631232) overnight before being spun down at 1500 x g for 45 minutes. For *in vivo* assays, particles were concurrently spun in 30mL increments in an ultracentrifuge at 24,000 rpm for 2hrs. Resulting pellets were supplemented with PBS and left to resuspend overnight at 4°C. Concentrated stocks were aliquoted and stored at -80°C. Optimization of large scale production, concentration, and characterization was guided by previously published protocols ^77^.

#### Vector Copy Number

Lentiviral preparations were quantified for vector copy number using the Lenti-X™ qRT-PCR Titration Kit (Takara Bio Cat: #631235). In brief, during lentiviral preparation, aliquots were taken directly after viral supernatant was collected and filtered. RNA was isolated from each sample as per the manufacturer’s instructions and then frozen. On day of titering, samples were thawed and prepared on ice, on a lab bench thoroughly cleaned with RNaseZap™ RNase Decontamination Wipes (ThermoFisher, AM9788). Samples were diluted and plated as per the Lenti-X™ qRT-PCR Titration Kit instructions and then analyzed by qRT-PCR using a Thermo Fisher QStudio 6 FLX real-Time PCR System.

HIV p24 content was quantified using the PerkinElmer HIV p24 AlphaLISA kit (no longer available from PerkinElmer). Briefly, aliquots of unconcentrated and concentrated viral supernatant were diluted and plated per manufacturer’s instructions. AlphaLISA acceptor beads and biotinylated anti-HIV p24 antibodies were incubated at room temperature for 1 hour. Streptavidin donor beads were prepared and added to the samples in a dark BL2+ room (<100 lumens) and incubated in the dark at room temperature for 30 minutes. Plate was subsequently read on a PerkinElmer Envision Plate Reader and analyzed using the manufacturer’s AlphaLISA analysis protocol for GraphPad Prism.

#### Cell culture

LentiX-293T cells were cultured in Dulbecco’s modified Eagle’s medium (Gibco) supplemented with 10% fetal bovine serum (FBS), Gentamicin, Penicillin and Streptomycin. HEK293T cells were sub-cultured three times a week at ratios of either 1:10 or 1:20. A20s and NALM6 cells were both cultured in RPMI with similar supplementation. These cells were sub-cultured 3 times a week at a ratio of 1:10. Primary human Pan T cells from Stemcell Technology were cultured in RPMI supplemented with 10% FBS along with Gentamicin, Pen-Strep, 1x NEAA(Life Technologies), 10mM HEPES (Thermo Scientific), 1mM Sodium pyruvate, 55uM 2-Mercaptoethanol (Thermo).

#### Serum envelope protein binding assay

LentiX HEK293T cells were transfected as previously described with 1ug of plasmid encoding the target envelope protein. 16 hours later, human serum samples of known MeV IgG titer were thawed and heat inactivated for 20 minutes at 56°C. Following heat inactivation, stepwise 10-fold dilutions were made of the serum samples with pre-warmed DMEM (formulated as previously described). The transfected cells were then lifted and re-plated in serum dilutions in triplicate. Cells were left to incubate on ice for 1 hour. Following incubation, cells were stained with fluorescent antibodies against Human IgG(APC) (Biolegend, clone: M1310G05) and HA-tag (BioLegend, clone: 16B12) using standard indirect flow cytometry staining protocol. Cells were then analyzed by flow cytometry using a Beckman CytoFLEX LX.

#### *In vitro* CAR T cell activation and killing assay

Re-targeted LV was prepared as previously described, in brief: HEK293s, either unmodified or stably expressing co-stimulatory αCD3-hCD80, were transfected with env plasmids encoding VSV-G or MeV/DMV-H-aCD7^VHH^; DMV/MeV-F^311^ as well as plasmids encoding a anti-CD19-CAR-GFP transgene. Donor Pan T cells were cultured and stimulated with Dynabeads™ Human T-Activator CD3/CD28 beads (Gibco™ 11131D) for 24 hours. Dynabeads were then removed from solution via magnets, and Pan T cells were washed following Dynabead manufacturer’s instructions. CD19+ NALM6 cells and activated Pan-T cells were then plated in various effector-target ratios and 10uL of 100-fold concentrated re-targeted LV was added to corresponding wells. 3 days and 6 days post-plating, aliquots from each well were taken, stained with aCD19 (Biolegend HIB19) antibody to determine NALM6 populations and analyzed by flow cytometry.

#### Surface expression

For surface expression 1E6 number of HEK293s were seeded and the next day 1ug of envelope plasmid was transfected with BLANK. For HA tagged proteins 24 after transfections cells were stained with anti HA antibody (BioLegend, clone: 16B12) for 1 hour. For surface of untagged MeV-H and MeV-F 10ug/mL of primary mouse anti-MeV-H (CL55) and mouse anti-MeV-F (Y503) were incubated for 2 hours at 4°C followed by a wash with cold MACS buffers then incubated with anti-mouse IgG (Biolegend, clone: Poly4053) antibody for 1 hour.

#### Serum Neutralization Assay

Serum donor samples were received from the Mass General Brigham BioBank and stored at -80°C. Samples were quantified for human IgG against MeV using a MeV IgG ELISA kit (Abcam: ab108750). Seropositivity was determined using a cut-off control well following the manufacturer’s instructions.

Re-targeted LV was prepared as previously described, in brief: unmodified HEK293T producer cells were transfected with env plasmids encoding different env protein variants as well as a transgene plasmid encoding aCD19-CAR-GFP. Supernatant was collected as previously described. Human serum samples of known MeV IgG titer were thawed and heat inactivated for 20 minutes at 56°C. Following heat inactivation, stepwise 10-fold dilutions were made of the serum samples with pre-warmed DMEM (formulated as previously described) in a 96well plate. Retargeted LV was then plated into the serum dilutions and left to incubate on ice for 1 hour. Following incubation, A20 lymphoma cells were plated in the wells. 48 hours later aliquots of each well were collected and analyzed by flow cytometry. “Remaining transduction” was calculated by dividing the %GFP+ cells for a well of a given LV and serum dilution by the %GFP+ cells for that LV in the control well containing no serum.

#### Multiplex Immunofluorescence Staining, Imaging, and Analysis

After mice were sacrificed, tissues were harvested and fixed in neutral buffered formalin for 20 hours. Tissus were then embedded in paraffin, and sectioned at 4-5 μm. Tissues were deparaffinized and hydrated by baking at 65°C for 15 minutes followed by a xylene and an ethanol gradient. Antigen retrieval was performed with a citrate-based buffer (Vector Labs) for 45 minutes at 100°C. Following antigen retrieval, cyclic immunofluorescence staining and imaging was performed using a Lunaphore Comet with primary antibodies targeting human CD3 (CD3-12, Abcam) mouse F4/80 (D2S9R, Cell Signalling), mouse Arginase-1(D4E3M, Cell Signaling), and GFP (600-101-215, Rockland). Secondary staining was performed with Donkey Anti-Rabbit IgG AlexaFluor 647+, Donkey Anti-Rabbit IgG AlexaFluor 555+, or Donkey anti-Goat IgG AlexaFluor 647+ (Life Technologies). Background subtraction was performed with Horizon Viewer (Lunaphore). Corrected images were then analyzed using Qupath^78^.

Areas of tissue folding or large tissue artifacts were excluded from analysis. On analyzable tissue, cell segmentation was performed using Instaseg utilizing signal outputs from all fluorescent parameters except for GFP^79^. Cell classifiers were then trained to classify segmented cells based upon their staining for DAPI, CD3, F4/80, and Arginase-1 (CD3+,F4/80-, Arginase-1-; CD3+,F4/80+, Arginase-1-; CD3-,F4/80+,Arginase-1-; CD3-,F4/80-,Arginase-1+; CD3-, F4/80-,Arginase-1-, Debris/RBCs, Annuclear cells) with an average of 130 annotation per sample. Annotations were made blinded to GFP signal. A Random trees classifier was utilized with consideration of all available fluorescent and geometric parameters except for those directly related to GFP signal. Model classification was manually reviewed to ensure accuracy.

GFP-positivity was defined using a threshold of up to 3.5 standard deviations above the mean background fluorescence of the analyzed area of the image (as this would yield < 5 false positive cells per 150,000 cell section due to background alone). Following thresholding, manual review was of all cells in all categories above the threshold. Mis-classified cells were reclassified appropriately. Cells with GFP staining that were wholly discordant with cellular staining or not associated with true cells were excluded as staining/imaging artifacts. CD3+, F4/80+, Arginase-1-cells (likely representing macrophage-lymphocyte pairs or phagocytosed T cells) were excluded from this analysis. Annotations with cell area <12 μm2 were excluded from final analysis as these were primarily cell fragments and debris.

