## Extended data figures for "*In vivo* generation of chimeric antigen receptor T cells using optimally retargeted and functionalized lentiviral vectors with reduced immune clearance"

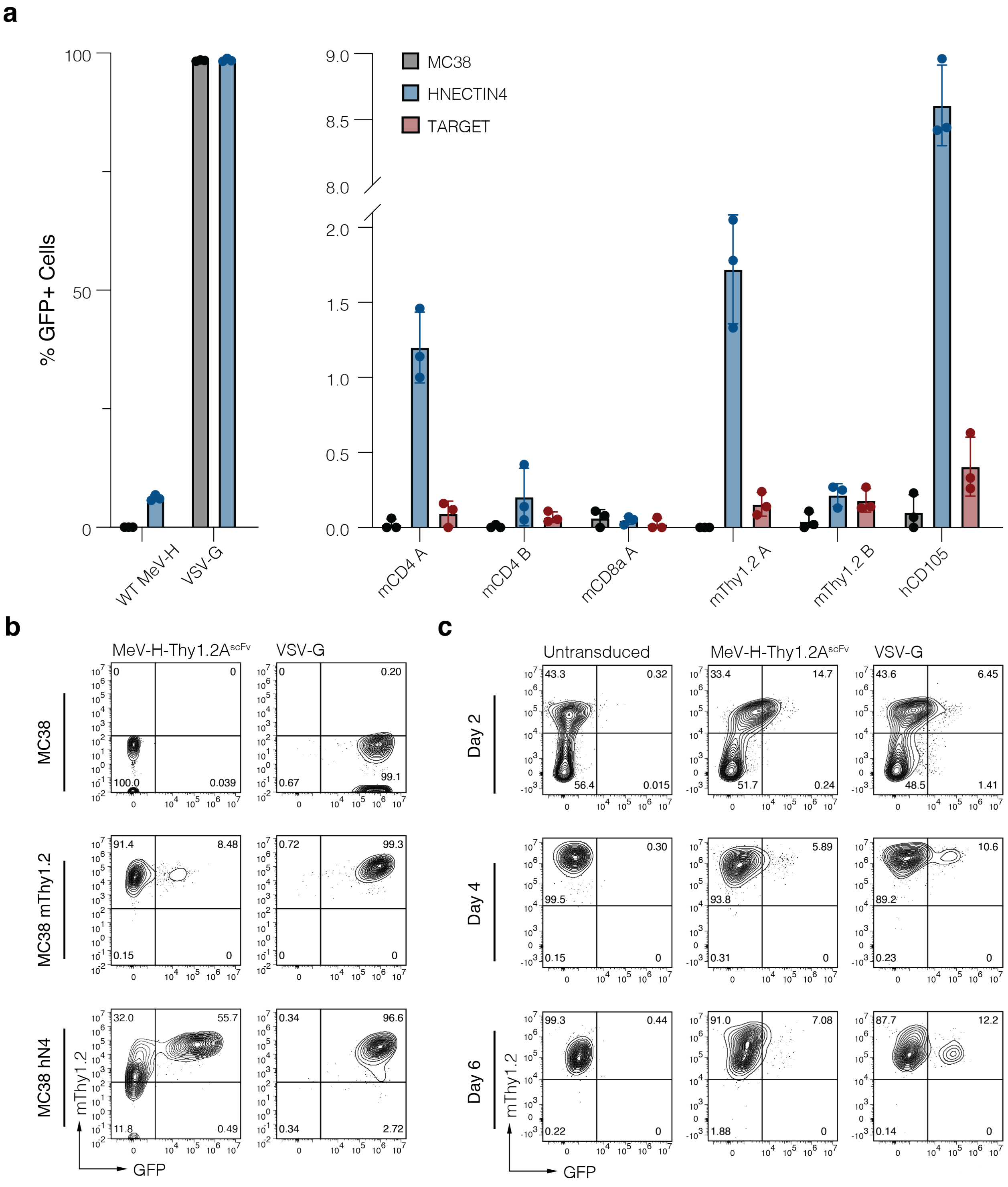
a
% GFP+ Cells
100
50
0
WT MeV-H
VSV-G
MC38
HNECTIN4
TARGET
mCD4 A
mCD4 B
mCD8a A
mThy1.2 A
mThy1.2 B
hCD105
b
MeV-H-Thy1.2AscFv
VSV-G
MC38
MC38 mThy1.2
MC38 hN4
mThy1.2
GFP
c
Untransduced
MeV-H-Thy1.2AscFv
VSV-G
Day 2
Day 4
Day 6
mThy1.2
GFP

Extended Data Figure 1: scFvs are poor targeting domains for retargeted MeV pseudotyped LVs.

(A) MeV-H-scFv pseudotypes were generated from hybridoma sequencing. MeV-H-scFv were applied to WT MC38 (grey), NECTIN4 expressing MC38s (blue) and scFv target matched expressing MC38 (red). (B) The most promising targeted pseudotype from (A) MeV-H-Thy1.2A$^{scFv}$ , was concentrated with Lenti-X then applied to WT MC38, NECTIN4 MC38 expressing and Thy1.2 MC38 cells and transduction analyzed with flow cytometry. (C) Primary activated whole mouse splenocytes were transduced with VSV-G or MeV-H-Thy1.2A$^{scFv}$ pseudotyped LVs and transduction measured by flow cytometry at days 2, 4 and 6 post application of LV.

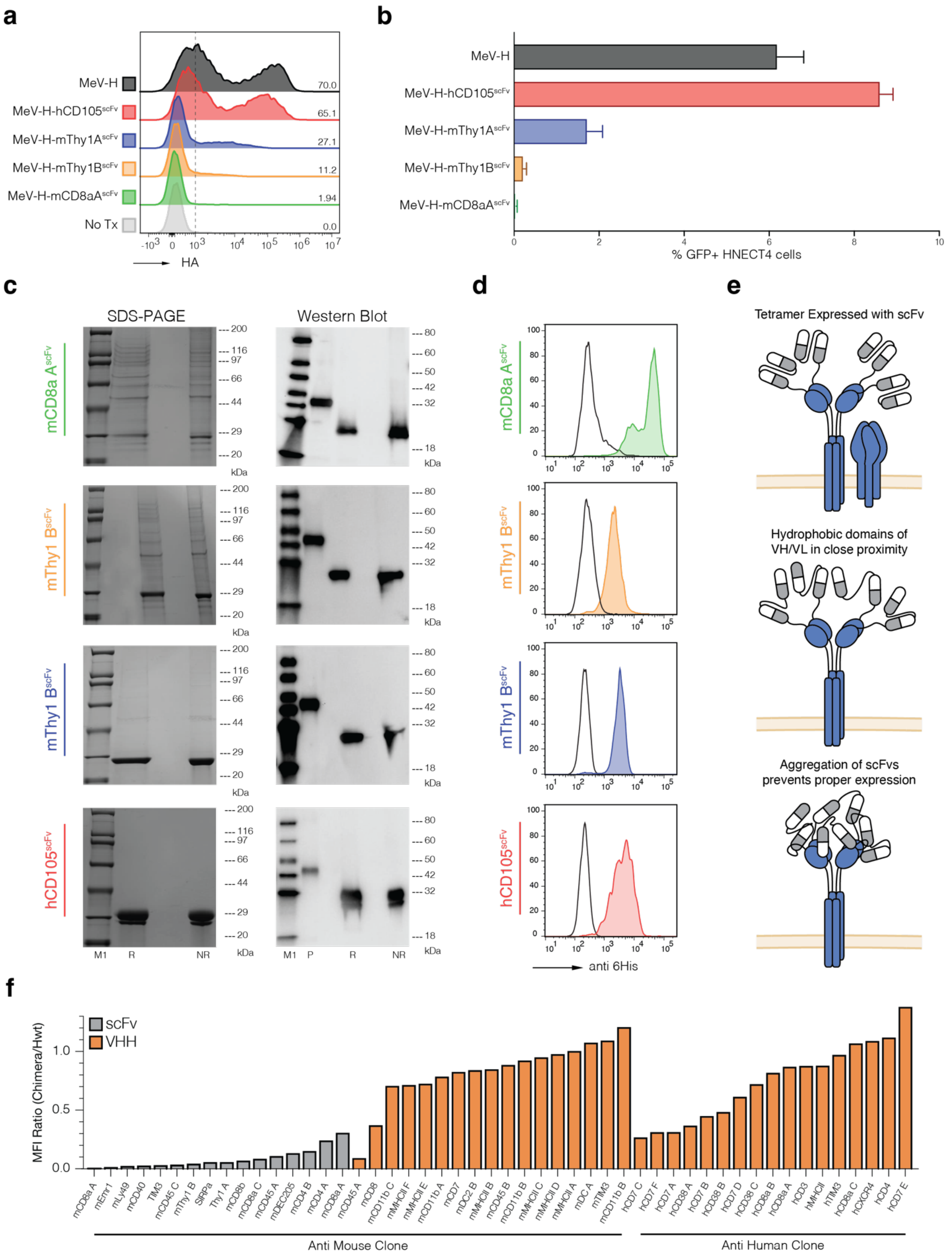
a
b
c
d
e
f
MeV-H
MeV-H-hCD105scFv
MeV-H-mThy1AscFv
MeV-H-mThy1BscFv
MeV-H-mCD8aAscFv
No Tx
HA
% GFP+ HNECT4 cells
SDS-PAGE
Western Blot
anti 6His
Tetramer Expressed with scFv
Hydrophobic domains of VH/VL in close proximity
Aggregation of scFvs prevents proper expression
MFI Ratio (Chimera/Hwt)
scFv
VHH
Anti Mouse Clone
Anti Human Clone

Extended Data Figure 2: Surface expression of Env-scFv fusion is critical for functional pseudotyping

(A) Surface expression of MeV-H-scFv on HEK239s. (B) Transduction of NECTIN4 expressing MC38s treated with LVs pseudotyped with MeV-H proteins from (A). (C) SDS-PAGE and western blot of soluble scFvs purified from HEK293s with a 6His tag Ni+ bead affinity chromatography. (D) Flow cytometry of MC38 expressing cognate targets for the scFvs in (C) stained using the purified soluble scFvs as a primary and an α6His antibody as secondary. E) Proposed mechanism for the ineffective and poor surface expression of scFvs in the context of MeV-H-scFvs. (F) MeV-H-scFv:WT MeV-H MFI ratio of all scFvs and VHH tested in this paper.

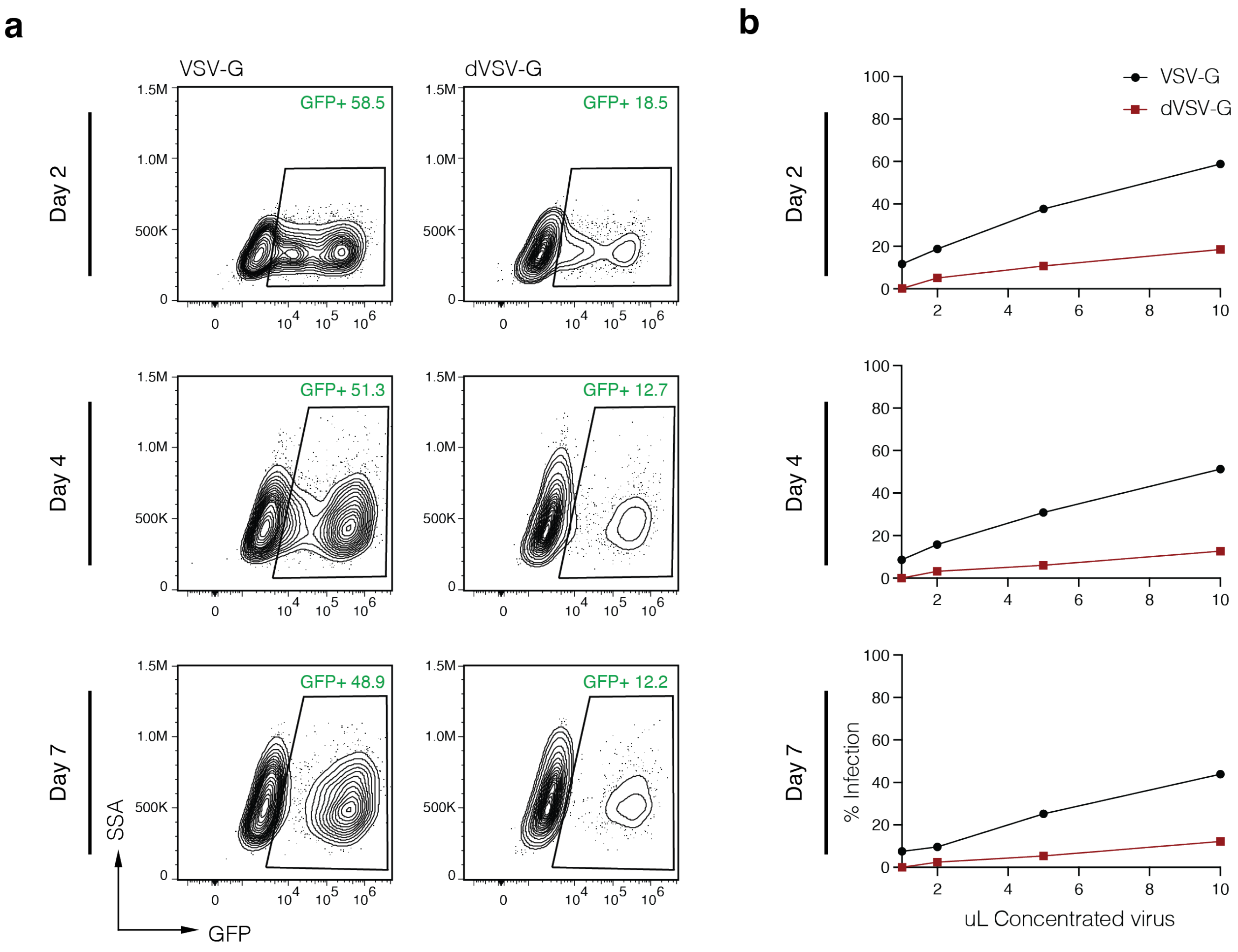


Extended Data Figure 3: dVSVG has substantial off target activity in vitro.

(A) Flow cytometry plots of MC38 cells treated with VSV-G and dVSV-G pseudotyped LVs carrying a GFP transgene on MC38 cells at day 2, 4 and 7 post treatment. (B) Quantified transduction of VSV-G and dVSV-G from (A) across different volumes of LVs.

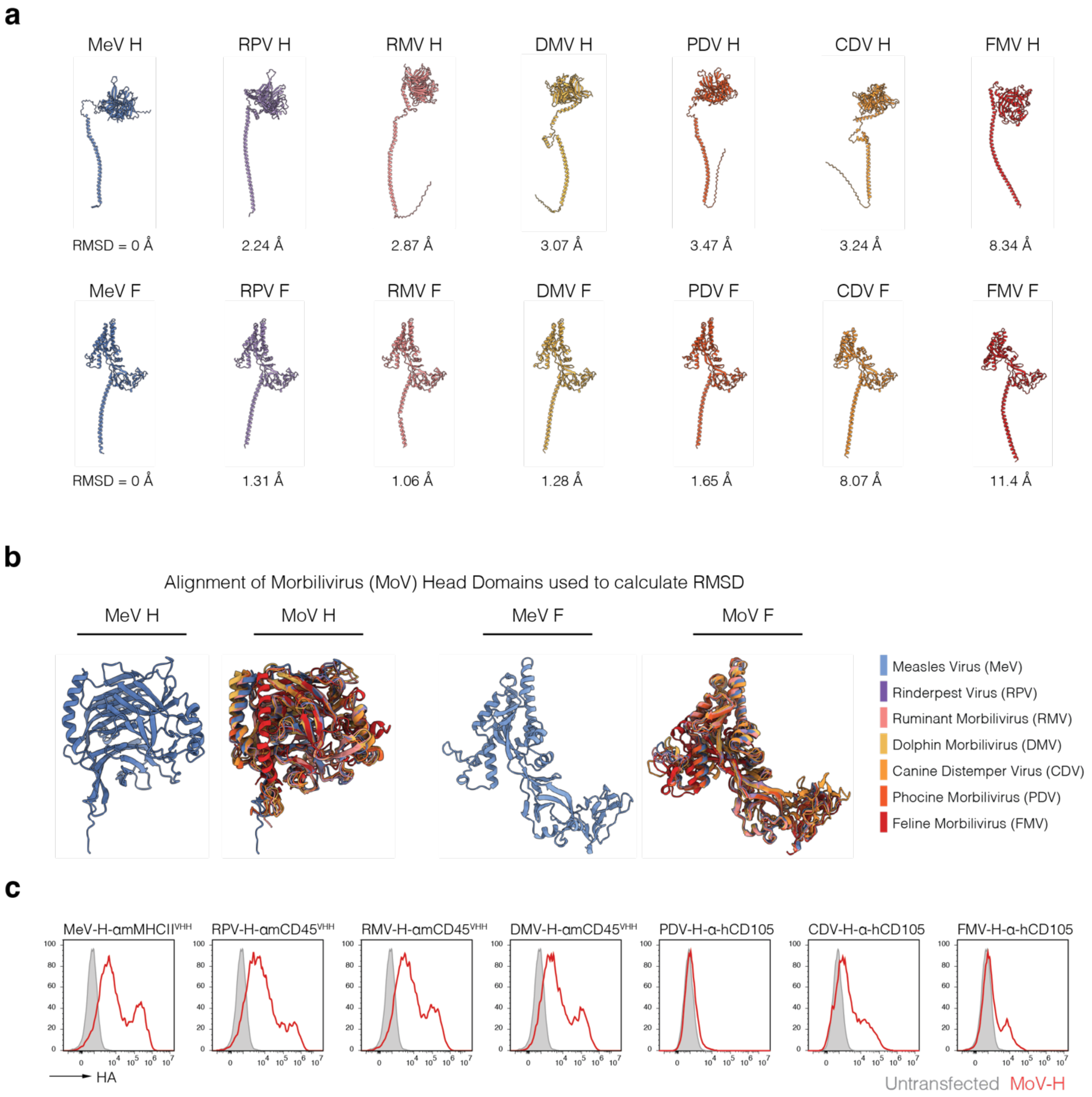


Extended Data Figure 4: MoV envelope proteins are structurally similar

(A) AlphaFold3 predictions of MoV H and F complexes (showing a single protomer) with root mean squared deviation (RMSD) from the MeV prediction. Reported RMSD excludes all of the transmembrane helix. (B) Alignment of MoV-H and MoV-F extravirion domains used to calculate RMSD. (C) Surface expression data of MoV-H (grey, untransfected; red, MoV-H).

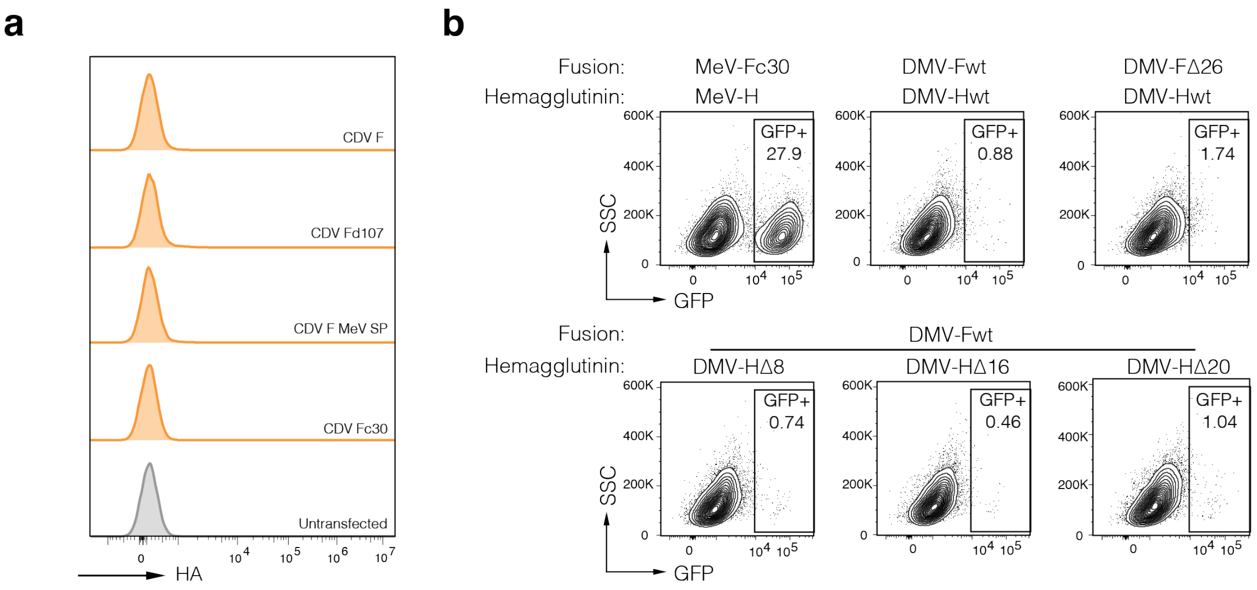


Extended Data Figure 5: Attempts to optimize wt CDV/DMV envelopes are unsuccessful

(A) Surface expression of engineered CDV variants assessed by flow cytometry, (CDV Fd107 - a 107aa truncation to the endogenous signal peptide, MeV SP - Measles Fusion Protein Signal Peptide, CDV-Fc30 that is truncated 30aa from the amino-terminus). (B) Transduction of variants of DMV H and F proteins with varying truncations, each virus had an H protein fused to the aMHCII-VHH and the cells used were A20 Leukemia cells. Analyzed by Flow cytometry.

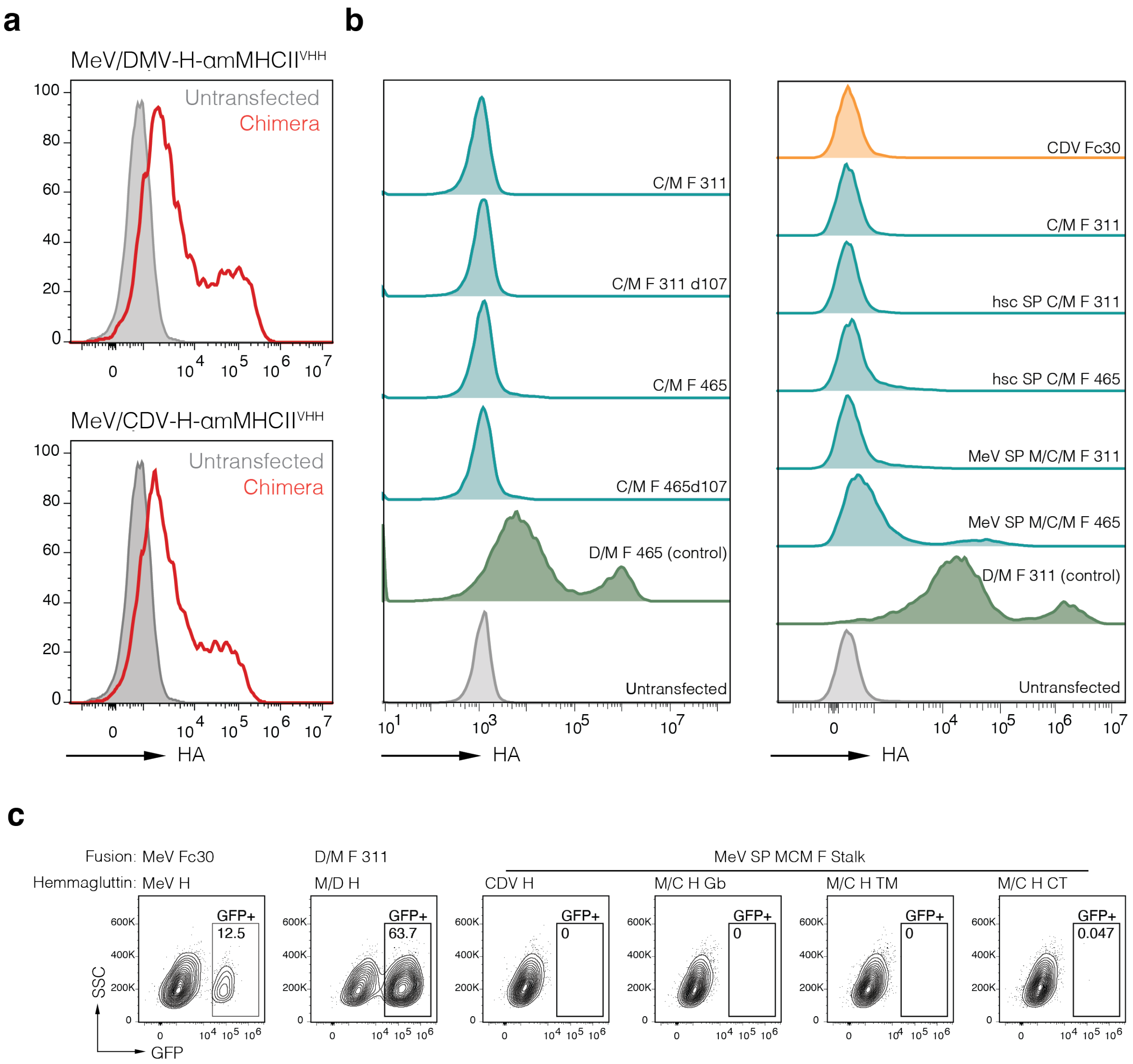


Extended Data Figure 6: Surface expression of DMV/CDV optimization is unsuccessful

(A) Surface expression of chimeric H proteins. (B) Surface expression of engineered variants of CDV/MeV-F chimeras. (C/M #: CDV/MeV chimeric splice site residue number; d107: a 107aa truncation to the endogenous signal peptide; hsc SP: heterologous signal peptide; MeV SP M/C/M F: variant using MeV SP and the first 19 residues from MeV F) (C) Transduction attempts with underlined variant in (B) with variants of the M/C H chimeras.

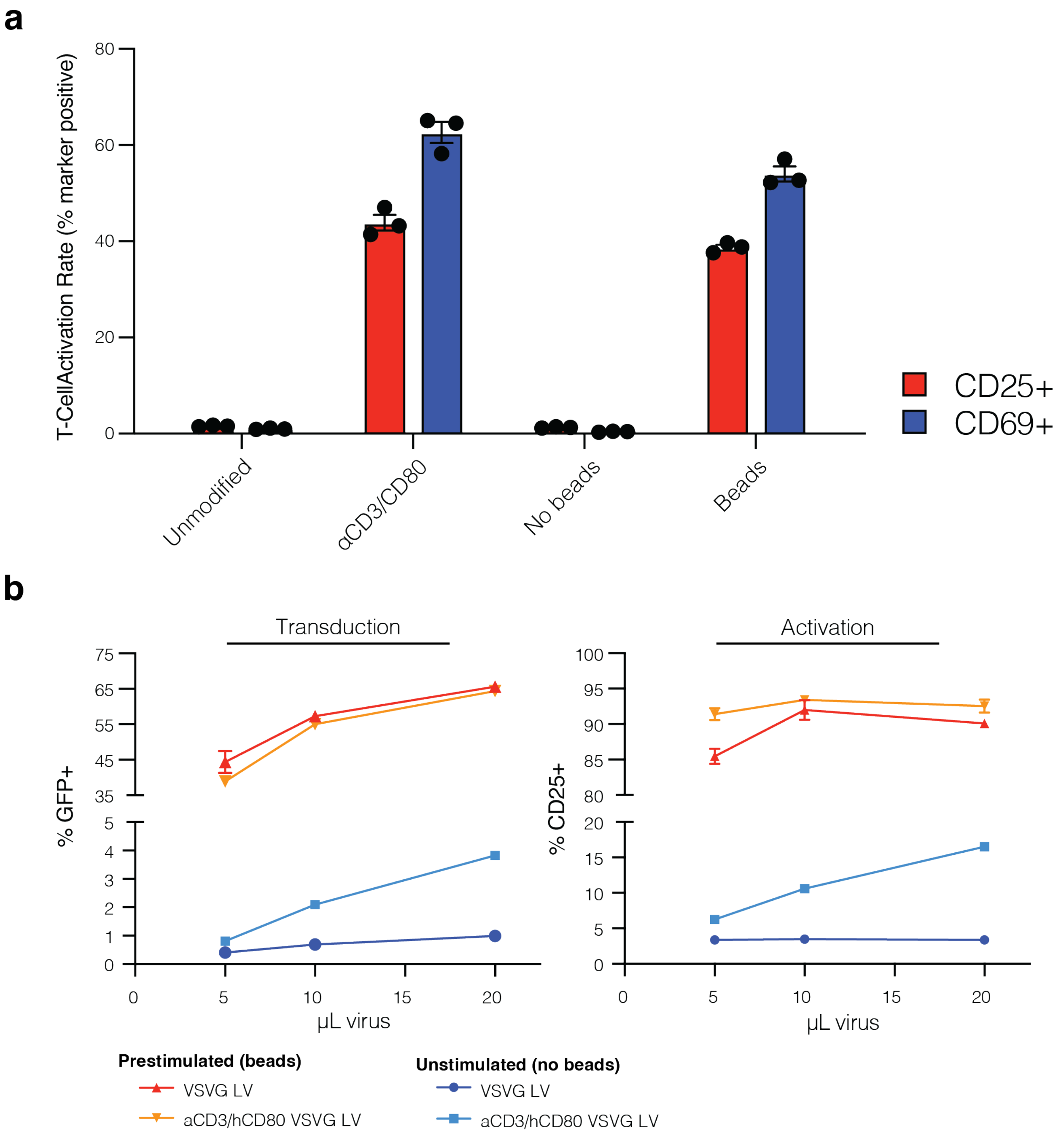


Extended Data Figure 7

A) Coculture of human pan T cells with unmodified HEK cells or HEK cells engineered to surface express αCD3 and CD80. B) LVs produced from the costim HEK cells activate and transduce unstimulated T cells at higher rates than LVs produced from unmodified HEK cells.

**a**

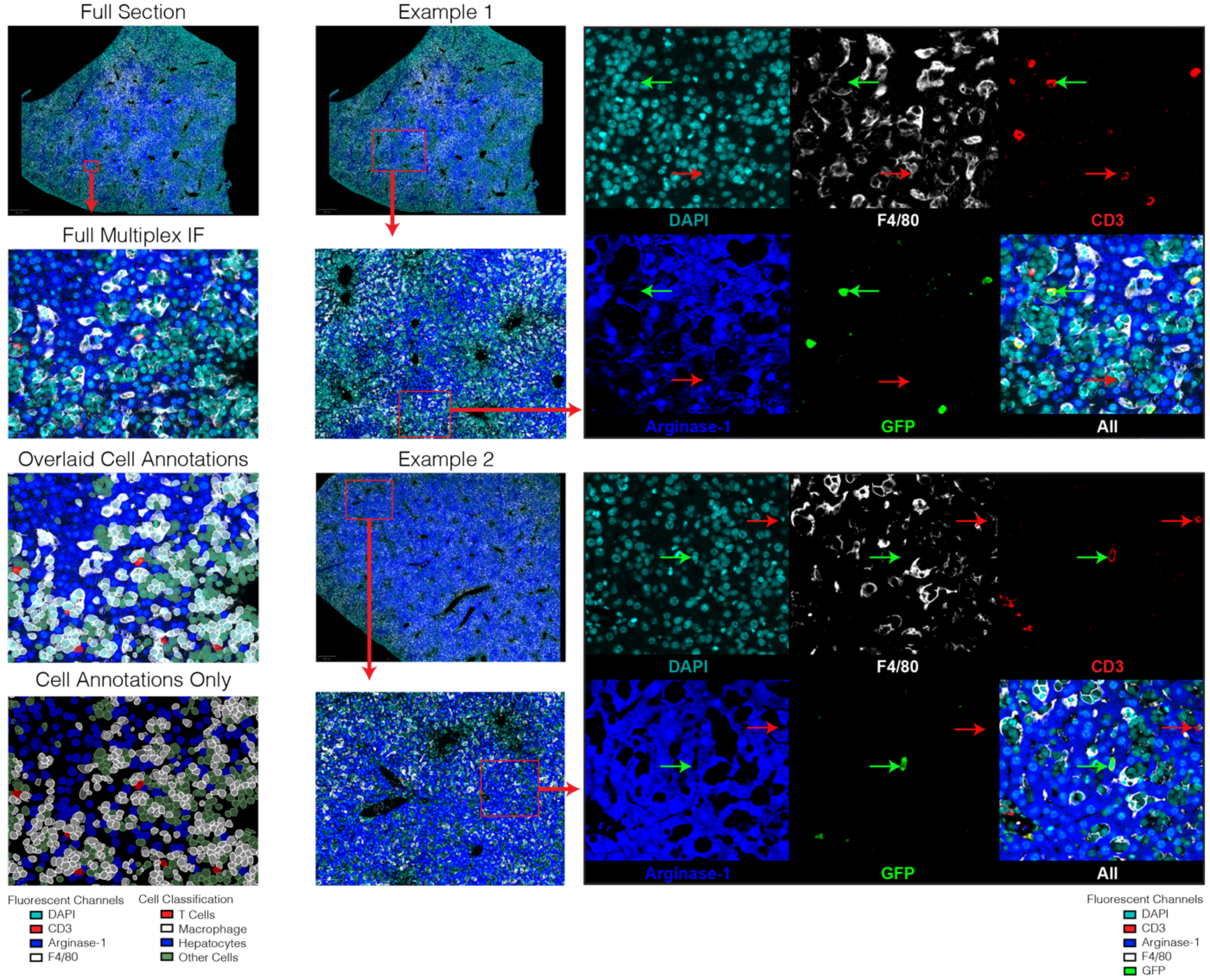


**c**

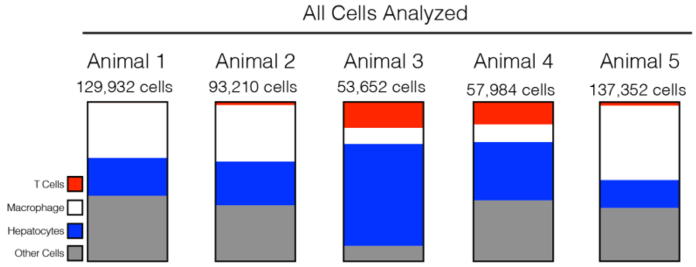


**d**

Composition of GFP +/- subsets

| | Animal 1 | | | Animal 2 | | | Animal 3 | | | Animal 4 | | | Animal 5 | | |
|---|---|---|---|---|---|---|---|---|---|---|---|---|---|---|---|
| | GFP- | GFP+ | % GFP+ | GFP- | GFP+ | % GFP+ | GFP- | GFP+ | % GFP+ | GFP- | GFP+ | % GFP+ | GFP- | GFP+ | % GFP+ |
| T Cells | 764 | 58 | 7.592% | 1785 | 67 | 3.754% | 8603 | 127 | 1.476% | 7757 | 383 | 4.937% | 2370 | 731 | 30.844% |
| Macrophage | 44682 | 1 | 0.002% | 32977 | 0 | 0.000% | 5379 | 0 | 0.000% | 6435 | 0 | 0.000% | 63981 | 1 | 0.002% |
| Hepatocytes | 30720 | 0 | 0.000% | 25404 | 0 | 0.000% | 34164 | 0 | 0.000% | 21109 | 0 | 0.000% | 23985 | 0 | 0.000% |
| Other Cells | 53706 | 1 | 0.002% | 32976 | 1 | 0.003% | 5379 | 0 | 0.000% | 22300 | 0 | 0.000% | 46284 | 0 | 0.000% |

**e**

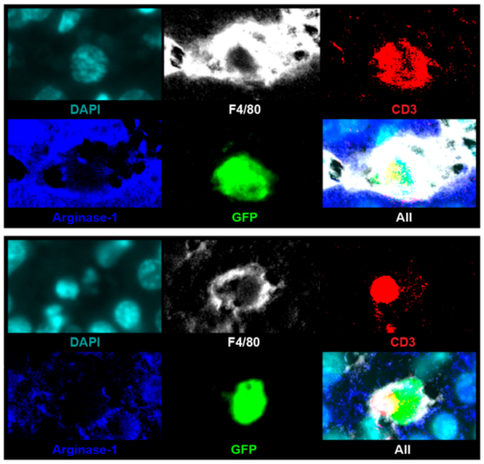

### Extended Data Figure 8

Representative multiplex immunofluorescence images of *in vivo* liver samples for the determination of transduction specificity. A) Representative example of full-section tissue staining and imaging of *in vivo* liver tissue (far left) with magnification (middle left) and cell segmentation and results of cellular classification model both with (middle right) and without (far right) overlay of IF imaging: Macrophage (F4/80+, CD3-, Arginase-1-), T Cells (F4/80-, CD3+, Arginase-1-), Hepatocytes (F4/80-, CD3-, Arginase-1+), or other cells (F4/80-, CD3-, Arginase-1-). B) Representative examples demonstrating specificity of GFP expression to T cells within animal livers. Green arrows indicate transduced T cells, while red arrows indicate non-transduced T cells. C) Results of cellular classification across whole-section multiplex IF tissue sections. D) Numeric breakdown of absolute number of cells classified as GFP+/- for each cell type in each sample. E) Example of Macrophage/T-Cell complexes that were excluded from analysis.

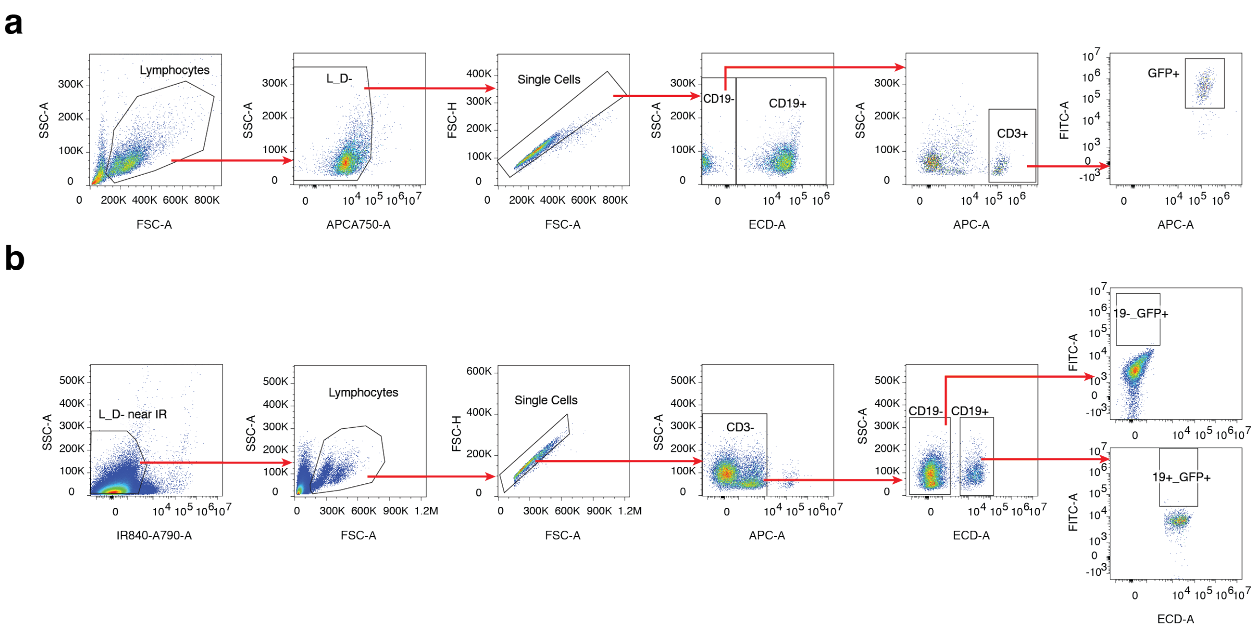


### Extended Data Figure 9

Gating strategies for flow cytometry analysis in figure 5. A) Global analysis of peripheral bleeds and tissue samples at endpoint with channel near IR live/dead (APCA750), hCD19 (ECD), CD3 (APC), GFP/CAR (FITC). B) Investigation of off-target transduction in blood collected at endpoint with channels IR live/dead (IR840-A790), hCD19 (ECD), CD3 (APC), GFP/CAR (FITC) (Extended Data Table 3)

### Extended Data Table 1

scFv and VHH sequences.

### Extended Data Table 2

Donor patient cohort information, MeV IgG score determined by MeV IgG ELISA kit following quantification instructions from manufacturer (Abcam: ab108750) .

### Extended Data Table 3

Off target counts.

### Extended Data Table 4

Plasmids generated in this study.
