## Extended data table 1 for "*In vivo* generation of chimeric antigen receptor T cells using optimally retargeted and functionalized lentiviral vectors with reduced immune clearance"

| | Clone name | Original Source Name | type | target | Source | Sequence |
|---|---|---|---|---|---|---|
| 1 | mCD45 A scFv | YW 62.3.20 | scFv | mCD45 | Culture Collections | QVQLKESGPGLVQPSQTLSLTCTVSGFSLTSYTVHWVRQPTGKGLEWMGIIWTDGSTDYNSALKSRLRISRDTSKSQVFLKMNSPQTEDMATYYCARGPRYRYKGYFDYWGQGVMVTVSSGGGGSGGGGSGGGGSNTVMTQSPTSMFISVGDRVTMNCKASQTVGPNVDWYQQKTGQSPKLLIYGASNRYTGVPDRFTGSGSGTDFTLTISNIQAEDLAVYYCLQYNYYPYTFGAGTKLALKGGGSGGGSYPYDVPDYA |
| 2 | mCD45 B scFv | MB23G2 | scFv | mCD45 | ATCC | MNFSNTLVFLLFLLKGILCEVQLVESGGGLVQPGRSLKLSCLASGFSFSNYGMNWIRQAPGKGLEWVATISSSSSHIYYADTVKGRFTISRENAKNTLYLQMTSLRSEDTALYYCASLTGNWGQGVMVTVSSGGGGSGGGGSGGGGSDIVMTQGALPNPVPSGESASITCHSSKSLLHSNGKTFLSWYLQRPGQSPQLLIYWMSTRASGVSDRFSGSGSGTDFTLKISSVEAEDVGVYYCQQFLEYPLTFGSGTKLEIKGGGSGGGSHHHHHH |
| 3 | mCD45 C scFv | MB4B4 | scFv | mCD45 | ATCC | EVKLAESGGGLVQPGNSLTLSCVASGFTFSHYGMHWIRQAPKKGLEWIAMIYHDSNKMYYADTVKGRFTISRDNSKNTLNLEMNSLRSEDTAMYHCAASMRYPLFDYWGQGVMVTVSSGGGGSGGGGSGGGGSDVQMTQSPSNLAASPGESVSINCKASKSISKYLAWYQQKPEKANKLLIFSGSTLHSGTPSRFSGSGSGTDFTLTIRDLEPEDFGLYYCQHYSEYPLTFGSGTKLEIKGGGSGGGSYPYDVPDYA |
| 4 | mCD45 D scFv | M1/89.18.7.HK | scFv | mCD45 | ATCC | QVQLKESGPGLVKPSLTLSLTCTVSGFSLNSYGVIWVRQPPGKGLEWLGVKWGYGNTNYNSALKSRLNINRDTSKSQVFLKMDNVQTEDTAMYFCARSRFNYGGPLDYWGQGVMVTVSSGGGGSGGGGSGGGGSQAVLTQPNSVSTSLGSTVKLSCTLSSGNIENNYVHWYQQYEGRSPTTMIYNDDKRPDGVPDRFSGSIDSSSNSAFLTINNVEIEDEAIYFCHSYVSSIIFGGGTKLTVLGGGSGGGSYPYDVPDYA |
| 5 | mCD45 E scFv | M1/9.3.4.HL.2 | scFv | mCD45 | ATCC | QVQLKESGPGLVKPSLTLSLTCTVSGFSLNSYGVIWVRQPPGKGLEWLGVKWGYGNTNYNSALKSRLNINRDTSKSQVFLKMDNVQTEDTAMYFCARSRFNYGGPLDYWGQGVMVTVSSGGGGSGGGGSGGGGSQAVLTQPNSVSTSLGSTVKLSCTLSSGNIENNYVHWYQQYEGRSPTTMIYNDDKRPDGVPDRFSGSIDSSSNSAFLTINNVEIEDEAIYFCHSYVSSIIFGGGTKLTVL |
| 6 | mCD45 F scFv | 14.8 | scFv | mCD45 RA | Culture Collections | QVQLKESGPGLVQPSQTLSLTCTVSGLSLASNSVSWIRQPPGKGLEWMGVIWSNGGTDYNSAIKSRLSISRDTSKSQVFLKMNSLQTEDTAMYFCATFYYSSAYWGQGTLVTVSSGGGGSGGGGSGGGGSDIVMTQAPLSVSVTPGESASISCRSSKSLLHSKGFTYVYWYLQKPGKSPQLLISRMSNLASGIPDRFSGSGSETDFTLKISRVEAEDVGIYYCGQLLENPWTFGGGTKLELK |
| 7 | mCD8b A scFv | YTS 156.7.7 | scFv | mCD8b | Culture Collections | EVKLLESGGGLVQPGGSLRLSCAASGFTFNDFYMNWIRQPPGQAPEWLGVIRNKGNGYTTEVNTSVKGRFTISRDNTQNILYLQMNSLRAEDTAIYYCARGGPYYYSGDDAPYWGQGVMVTVSSGGGGSGGGGSGGGGSDIQMTQSPSFLSTSLGNSITI |

| | | | | | | |
|---|---|---|---|---|---|---|
| | | | | | | TCHASQNIKGWLAWYQQKSGNAPQLLIYKASSLQSGVPSR FSGSGSGTDYIFTISNLQPEDIATYYCQHYQSFPWTFGGGT KLELKGGGSGGGSYPYDVPDYA |
| 8 | mCD8a A scFv | 53.6-72 | scFv | mCD8a (mLyt-2) | ATCC | EVQLVESGGGLVQPGRSLKLSCVASGFTFSDYGMNWIRQ APGKGLEWVAYISSSSGTLYYADTLKGRFTISRDNAKSTLY LQLSSLRSEDTALYYCVRHRYKYYGDNFDSGPFDYWGQG VMVTVSSGGGGSGGGGSGGGGSDTVLTQSPALAVSPGE RVTISCRASESVSTRMHWYQQKPGQQPKLLIYGASNLESG VPARFSASGSGTDFTLTIDPVEATDTATYFCQQSWHDPRT FGGGTKLELK |
| 9 | mCD8a B scFv | YTS 105.18.10 | scFv | mCD8a | Culture Collections | EVKLVESGGGLVQPGRSLKLSCAASGFNFNDYWMGWVR QAPGKGLEWIGEINKDSSTINYTPSLKDKFTISRDNAQNTLY LQMSKLGSEDTAIYYCARARGMMVLIIPHYFDYWGQGVMV TVSSGGGGSGGGGSGGGGSDTVLTQSPALAMSPGERITI SCRASESVSTRMHWYQQKPGQQPKLLIYGASNLESGVPA RFSGSGSGTDFTLTIDPVEANDTATYFCQQSWYDPWTFG GGTKLELKGGGSGGGSYPYDVPDYA |
| 10 | mCD8a C scFv | YTS 169.4.2.1 | scFv | mCD8a (mLyt-2) | Culture Collections | MAVLVLLLCLVTFPSCALSQVQLKESGPGLVQPSQTLSLTC TVSGFSLTSNSVHWVRQPPGKGLEWMGGIWGDGDTDYN SALKSRLSISRDTSKNQVFLKMNSLQTDDTAIYFCTPLIGSW YFDFWGPGTMVTASSGGGGSGGGGSGGGGSNIVMTQSP KSMSISVGDRVTMNCKASQNVRNNIAWYQQKPGQSPKLLI YYASYRYTGVPDRFTGDGFGTDFTLAINSVQAEDAAFYYC QRIYNSPYTFGAGTKLELIGGGSGGGSYPYDVPDYA |
| 11 | mThy1 A scFv | YTS 154.7.7.10 | scFv | mThy-1 | Culture Collections | MAVLVLLLCLLTSPSCVLSQVQLKESGPGLVKPSETLSLTC TVSGFSFTSYHISWVRQPPGKGLEWMGVIWGDGSTAYNS VFKSRLSISRDTSKSQVFLKMSSLKTEDTATYYCARDRDW ELGNWFAYWGQGTLVTVSSGGGGSGGGGSGGGGSDIQ MTQSPPSLSASLGDKVTITCQASQNINKYIAWYQQKPGKAP RQLIYYTSILVSGTPSRFSGSGSGRDYSFSISNVESEDIASY YCLQYDNLWTFGGGTKLELKGGGSGGGS |
| 12 | mThy1 B scFv | 30-H12 | scFv | mThy-1.2 | ATCC | MGLSWVFLVALLKGIQCEVQVVETGGGLVQPGSSLKLSCE SSGFTFSSTWMNWVRQAPGKGLEWVALVKDKYSNYEAN YAESVKGRFIISRDDSKNRVYLQMNTLRDQDTATYHCTRS RGYKNWYFDFWGPGTMVTVSSGGGGSGGGGSGGGGSD IQMTQSPPSLSASLGDKVTITCQASQNINNYIAWYQQKPGK APGLLILYTSTLVSGTPSRFSGSGSGRDYSFSISNVESEDIA SYYCLQYDNSPRTFGGGTKLELKGGGSGGGSHHHHHH |
| 13 | mThy1 C scFv | YBM 29.2.1 | scFv | mThy-1 | Culture Collections | MAVRVLLLCLVTFPSCALSQVQLMESGPGLVQPSETLSLTC TVSGFSLTTYNVHWVRQPPGKGLEWMGVMWSGGSTDYN SALKSRLSISRDTSKNQVLLKMNSLQSEDTTTYYCARAFYY YDGSYYHDYFDYWGQGVMVTVSSGGGGSGGGGSGGGG SEIVLTQSPTTMAASPGEKVTLTCHASSSVSYMHWYQQRS GASPKVWIYETSKLASGVPNRFSGSGSGTSYSLTINSMATE DAATYYCQQGSSYPLTFGSGTKLEIKGGGSGGGSYPYDVP DYA |

| | | | | | | |
|---|---|---|---|---|---|---|
| 14 | mCD4 A scFv | YTS 177.9.6.1 | scFv | mCD4 | Culture Collections | MAVLVLLLCLVTFPSCALSQVQLKESGPGLVQPSQTLSLTC<br>TVSGFSLTSNSVHWVRQPPGKGLEWMGGIWGDGSTDYN<br>SALKSRLSISRDTSKSQVFLKMNSLQTDDTAIYFCTRYYNS<br>YIYEGWYFDFWGPGTMVTVSSGGGGSGGGGSGGGGSDI<br>QMTQSPSSMSASLGDTVTINCLASQDIGNYLSWYQQKPGK<br>APKVMIYGATNLEDGVPSRFSGSRSGSDYSLTINSLGYDDE<br>GIYHCHQYYEYPLTFGSGTKLEIKGGGSGGGSYPYDVPDY<br>A |
| 15 | mCD4 B scFv | YTA 3.1.2 | scFv | mCD4 | Culture Collections | MQWLWTLLFLMVVAQSTQAQIQLVQSGPELKKPGESVKIS<br>CKASGYTFTNYGMYWVKQAPGQGLQYMGWINTETGKPTY<br>ADDFKGRFVFFLETSASTAYLQINNLKNEDMATYFCARPSR<br>YYDGPNWFGYWGQGTLVTVSSGGGGSGGGGSGGGGSD<br>IQMTQSPHSLSASLGETVSIECLASEGISNYLAWYQQKPGK<br>SPQLLIYYASSLQDGVPSRFSGSGSGTHYSLKISNMQPEDE<br>GIYYCQQAYKFPLTFGSGTKLEIKGGGSGGGSHHHHHH |
| 16 | mCD4 C scFv | YTS 191.1.1.2 | scFv | mCD4 | Culture Collections | EVQLVESGGGLVQPGRSLKLSCAASGFTFSDYNMAWVRQ<br>APKKGLEWVATIDYDGSTTYYRDSVKGRFTISRDNTKNTLF<br>LQMDSLKSEDTATYYCAKQAYYDGFYHVDYFAYWGQGVV<br>VTVSSGGGGSGGGGSGGGGSDTVLTQSPALAVSLGQRVT<br>ISCKASESVSSSRYSYMHWYQQKPGQQPKLLIYHASNLES<br>GVPARFSGSGSGTDFTLNIDPVEADDIATYFCQQSWNDPP<br>TFGGGTELELKGGGSGGGSYPYDVPDYA |
| 17 | mCD4 D scFv | GK1.5 | scFv | mCD4 | ATCC | MKCSWIILFLMALTTGVNSEVQLQQDGAELGKPGTSVKLS<br>CKVSDYNIRRTYMHWVNQRPGKGLEWIGRIDPANGNTIYG<br>EKFKSKATLTADTSSNTAYMQLSQLKSDDTAIYFCAIGVQY<br>LDYWGQGVMVTVSSGGGGSGGGGSGGGGSDTVLTQSP<br>ALAVSPGERVTISCRATESVSTLIHWFQQRPGQQPKLLIYLT<br>SHLDSGVPARFSGSGSGTDFTLTIDPVEADDTATYYCQQT<br>WNDPWTFGGGTKLELKGGGSGGGSHHHHHH |
| 18 | mNK1.1 A scFv | PK136 | scFv | mNK1.1 | ATCC | EVQLLQSGAELVRPGASVKLSCTASGFNIKDDYMHWVKQR<br>PEQGLEWIGRIDPANINTKCAPKFQDKATITADTSSNTAYLQ<br>LSSLTSEDTAVYYCARSPYYPSYWGQGTLVTVSAGGGGS<br>GGGGSGGGGSDVLMTQTPLSLPVSLGDQASISCRSSQSIV<br>HSNGNTYLEWYLQKPGQSPKLLIYKVSNRFSGVPDRFSGS<br>GSGTDFTLKISRVEAEDLGVYYCFQGSHVPRTFGGGTKLEI<br>KGGGSGGGSYPYDVPDYA |
| 19 | mTIM3 | RMT3-23 | scFv | mTIM-3 | 1 | EVQLVESGGGLVQPGRSLKLSCAASGFTFSDFYMAWVRQ<br>APKKGLEWVASISYEGSSTYYGDSVMGRFTISRDNAKSTL<br>YLQMNSLRSEDTATYYCARQREANWEDWGQGVMVTVSS<br>GGGGSGGGGSGGGGSDIQMTQSPASLSASLGETVTIQCR<br>ASEDIYSGLAWFQQKPGKSPQLLIYGASSLQDGVPSRFSG<br>SGSGTQYSLKISSMQTEDEGVYFCQQGLKYPPTFGSGTKL<br>EIKGGGSGGGSYPYDVPDYA |
| 20 | hCD3 scFv | OKT3 | scFv | hCD3 | 2 | MGWSCIILFLVATATGVHSQVQLQQSGAELARPGASVKMS<br>CKASGYTFTRYTMHWVKQRPGQGLEWIGYINPSRGYTNY<br>NQKFKDKATLTTDKSSSTAYMQLSSLTSEDSAVYYCARYY |

| | | | | | | |
|---|---|---|---|---|---|---|
| | | | | | | DDHYCLDYWGQGTTLTVSSGGGGSGGGGSGGGGSDIVL TQSPAIMSASPGEKVTMTCSASSSVSYMNWYQQKSGTSP KRWIYDTSKLASGVPAHFRGSGSGTSYSLTISGMEAEDAA TYYCQQWSSNPFTFGSGTKLEINRGSEQKLISEEDLNAVG QDTQEVIVVPHSLPFKVVVISAILALVVLTIISLIILIMLWQKKP R |
| 21 | SIRPa | MY-1 | scFv | SIRPa | 3 | EVQLVESGGGLVQPGRSLKLSCAASGFTFSNSDMAWVRQ APTKGLEWVASITPSGGNTYYRDSVKGRFSVSRDNAKSSL YLQMDSLRSEDSATYFCARLDGYGGYPFFAYWGQGTLVT VSSGGGGSGGGGSGGGGSDTVLAQSPALAVSPGERVSIS CGASESVSNIIHWYQQRPGQQPKLLIYGASSLESGVPARFS GSGSGTDFTLNIDPVEADDTATYFCQQSWNDPLTFGSGTK LEIKGGGSGGGSYPYDVPDYA |
| 22 | mCD47 | Miap301 | scFv | mCD47 | 4 | MGWICIIFLVATATGVHSQVKLLQSGAALVKPGASVKMSCQ ASGYSFTDYWVTWVKQSHGQSLEWIGEIYPSNTVTNFNDN FKGKATLTVDKSTSTAYMELSRLTSEDSAIYYCTRLGNSGY RVGWFLYWGQGTLVTVISGGGGSGGGGSGGGGSDTVLT QSPALAVSPGERVTISCKASESVSSHMHWYQQKPGQQPK LLIYKASNLASGVPARFSGSGSGTDFTLTIDPVEADDTATYF CQQGWNGPFTFGAGTRLELKGGGSGGGSHHHHHH |
| 23 | mCD40 A | FGK4.5 | scFv | mCD40 | 5 | EVQLVESDGGLVQPGRSLKLPCAASGFTFSDYYMAWVRQ APTKGLEWVASISYDGSSTYYRDSVKGRFTISRDNAKSTLY LQMDSLRSEDTATYYCGRHSSYFDYWGQGVMVTVSSGG GGSGGGGSGGGGSDTVLTQSPALAVSPGERVTISCRASD SVSTLMHWYQQKPGQQPKLLIYLASHLESGVPARFSGSGS GTDFTLTIDPVEADDTATYYCQQSWNDPWTFGGGTKLELK GGGSGGGSYPYDVPDYA |
| 24 | Emr1 | F4/80 | scFv | mF4/80 | ATCC | MRVQIQFWGLLLLWTSGIQCDVQMTQSPYNLVASPGESVS INCKASKSISKYLAWYQQKPGKANKLLIYEGSTLQSGIPSRF SGSGSGTDFTLTIRSLEPEDFGLYYCQQHNEYPLTFGSGT KLEIKGGGGSGGGGSGGGGSQVQLQQSGAELVKPGTSVK LSCKASGYTFTRNHMNWVKQTTGQGLEWIGRINPGTGGT SYNVNFKGKATLTVDESSSTAFMQLSSLTPEDSAVYYCAR GDSYWYFDFWGPGTMVTVSSGGGSGGGSYPYDVPDYA |
| 25 | DEC205 | DEC-205 | scFv | mDEC-205 | ATCC | EVKLLESGGGLVQPGGSLRLSCAASGFTFNDFYMNWIRQP PGQAPEWLGVIRNKGNGYTTEVNTSVKGRFTISRDNTQNIL YLQMNSLRAEDTAIYYCARGGPYYYSGDDAPYWGQGVMV TVSSGGGGSGGGGSGGGGSDIQMTQSPSFLSTSLGNSITI TCHASQNIKGWLAWYQQKSGNAPQLLIYKASSLQSGVPSR FSGSGSGTDYIFTISNLQPEDIATYYCQHYQSFPWTFGGGT KLELKGGGSGGGSYPYDVPDYA |
| 26 | MuLVa | 83a25 | scFv | MuLV Ampho | ATCC | EVQLVESGGGLVQPGRSMKLSCAASGFTFSSFPMAWVRQ APTKGLEWVATISTSGGSTYYRDSVKGRFTISRDNVKSTLY LQMNSLRSEDTATYYCTRISTMMVLYWYFDFWGPGTMVT VSSGGGGSGGGGSGGGGSDIQMTQSPASLSASLGETVSI ECLASEDISNHLAWYQQKSGKSPQLLIYAASRLQDGVPSR |

| | | | | | | |
|---|---|---|---|---|---|---|
| | | | | | | FSGSGSGTRYSLKISGMQAEDEADYFCQQSTKYPYTFGAGTKLELKGGGSGGGSHHHHHH* |
| 27 | Ly49 | 4D11 | scFv | Ly49 | ATCC | EVQLVDSGGGLAQPGRSMRLSCAASGFTFSNYYMAWVRQAPTKGLEWVASISTGGSNTYYRDSVKGRFTISRDNVNSILSLQMDSLRSEDTATYYCTRQGRSGSYYLDYWGQGVMVTVSSGGGGSGGGGSGGGGSDIQMTQSPASLSASLGETVSIECLAGEGISNDLAWYHQKSGNSPQLLIYAASRLEDGVPSRFSGSGSGTRFSLKISGMQPEDEADYFCQQTFKYPFTFGSGTKLEIK |
| 28 | ahCD105 | ahCD105 | scFv | hCD105 | 6 | MAEVQLLESGGGLVQPGGSLRLSCAASGFTFSSYAMSWVRQAPGKGLEWVSAIYGSDGDTTYADSVKGRFTISRDNSKNTLYLQMNSLRAEDTAVYYCARVFYTAGFDYWGQGTLVTVSSGSDSNAGHASAGNTSDIELTQSPSSLSASVGDRVTITCRASQSISSSLNWYQQKPGKAPKLLIYAASSLQSGVPSRFSGSGSGTDFTLTISSLQPEDFATYYCQQAPAKPPTFGQGTKLEIKR |
| 29 | mTIM3 VHH | mH2 | VHH | mTIM-3 | 7 | QVQLVESGGGLVQAGGSLRLSCAASGFTFDDYAIGWFRQAPGKEREGVSCISSSDGSTYYTDSVKGRFTISSDNAKNTVYLQMNSLKPEDTAVYYCAADTTFFGCSLNRDYDYWGQGTQVTVSS |
| 30 | mDC VHH A | DC2.1 | VHH | mouse dendritic cells | 8 | QVQLQESGGGLVPPGGSLRLSCAASGFTFSSYGMSWVRQAPGKGLEWVSSISAYDTTSYADSVKGRFTISRDNAENTLYLQMSSLKPEDTAVYYCARSSYPMTYPSLRLSDYRGQGTQVTVSS |
| 31 | mDC VHH B | DC1.8 | VHH | mouse dendritic cells | 8 | QVQLQESGGGLVPPGGSLRLSCAASGFTFSSYGMSWVRQAPGKGLEWVSSISAYDTTSYADSVKGRFTISRDNAENTLYLQMSSLKPEDTAVYYCARSSYPMTYPSLRLSDYRGQGTQVTVSS |
| 32 | mCD45 VHH A | G7 | VHH | mCD45 | 9 | MAQVQLVESGGGLVHPGDSLRLSCAASGSVFNSATMGWYRQSPGSQRELVATIVVGTPTYADSVKGRFTISRDNAKNIVYLQMNSLKPEDTAVYYCNYRATYTSGYSRDYWGQGTQVTVS |
| 33 | mCD45 VHH B | 32 | VHH | mCD45 | 9 | MAQVQLVQSGGGLVQPGGSLRLSCAASGRAFNSAAMGWYRQAPGSQRELVASISAGTASYADAVKGRFTISRDYAKNIIYLQMNSLKPDDTAVYFCNYRTTYTSGYSEDYWGQGTQVTVS |
| 34 | mCD11b VHH A | V36 | VHH | mCD11b | 9 | MAQVQLVESGGGQVRAGGALRLACEVSGRTFSEDTVAWFRQLPGSAREFVAAISGSDDWNYYSDSVKGRFTISIDRAKTTAYLQMNSLKSEDTAVYYCAARLFGLPNIPGSYSDWGQGTQVTVS |
| 35 | mCD11b VHH B | VHHCD11b | VHH | mCD11b | 10 | QVQLQESGGGLVQAGGSHNLSCTASGITFSSLAMGWFRQTPGKEREFVANIMRSGSSVFYADSVRGRFTISRDNAKNTAHLQMNSLKPEDTAVYFCAATRGAWPAEYWGQGTQVTVSSGGLPETGGHHHHHH |

| | | | | | | |
|---|---|---|---|---|---|---|
| 36 | mCD11b VHH C | 81 | VHH | mCD11b | 9 | MAQVTLVESGGGLVPSGGSLRLSCAVSGRTSIDYAMAWFRQVPGKEREFVAGVDRGGDVTDYADFLKGRFTVSRDDAKETMNLKMNNLQPEDTAIYYCAVTNTIPTMRIPLTFIGYDHWGQGTQVTVS |
| 37 | mCD11b VHH D | 51 | VHH | mCD11b | 9 | MAQVQLQESGGGLVQAGASLTLSCAASGRNVEAYAMGWFRQAPGKEREFVATVEWSDGRPIYADSVKGRFTISEDTAKNTVDLLMNSLKPEDTAVYYCAARYSTGYYSASEYDYWGQGTQVTVS |
| 38 | mMHCII A | VHH7 | VHH | mMHCII | 7 | QVQLQESGGGLVQAGDSLRLSCAASGRTFSRGVMGWFRRAPGKEREFVAIFSGSSWSGRSTYYSDSVKGRFTISRDNAKNTVYLQMNGLKPEDTAVYYCAAGYPEAYSAYGRESTYDYWGQGTQVTVSSGGLPETGGHHHHHH |
| 39 | mMHCII B | N11 | VHH | mMHCII | 9 | MAQVQLVQSGGGLVQPGGSLGLSCAASGNIGSRDNMGWYRQAPGKQREWVATISGYGIATYRDSVKGRFTVAKDTAKNIVSLQMNYLTTEDTAVYYCYAYAVDSRNIFWSQGTQVTVS |
| 40 | mMHCII C | DC1 | VHH | mMHCII | 11 | QVQLQESGGGLVQAGGSLRLSCAASGRTFSRSAMGWFRQAPGKEREFVATISWSGGVTYYADSVKGRFTISRDNAKNTVYLQMNGLKPEDTAVYYCAAGYPEAYSAYGRESTYDYWGQGTQVTVSSGGLPETGGHHHHHH |
| 41 | mMHCII D | DC8 | VHH | mMHCII | 11 | QVQLQESGGGLVQPGGSLRLSCTASGFTFSTYYMSWVRKAPGKGPEWVSVMNSSGGDTRYADFVKGRFTISRDNAKNTLYLQMNSLKPEDTALYYCAQGRSDIYPTFTRGQGTQVTVSSGSLPETGGHHHHHH |
| 42 | mMHCII E | DC14 | VHH | mMHCII | 11 | QVQLQESGGGLVQAGGSLRLSCAASGRTFGYALGWFRQAPGKEREFVTAINWSGGRLGYADSVKGRFTISKDNYKNTLYLQMNGLKPEDTAVYYCAAGRGSTRSDDYDYWGQGTQVTVSSGSLPETGGHHHHHH |
| 43 | mMHCII F | DC15 | VHH | mMHCII | 11 | QVQLQESGGGLVQPGASLRLSCVSAFSLDHYSVAWFRQAPGKEREGVSCHSRSNKATDYADSVKGRFTISRDSAKRTVYLQMNNLKPEDTAVYYCAVKQWGMCTGGVWGRSQYDYWGQGTQVTVSSGSLPETGGHHHHHH |
| 44 | mMHCII G | 49 | VHH | mMHCII | 9 | MAQVQLVESGGGLVKPGGSLRLSCVVQGTLISTKAMGWYRQAPGKQREVVATITTTNTPGYVDPVKGRFTISRDNAKNTVYLQMDSLKSEDTAVYYCNLLSWRSDYNWGQGTQVTVS |
| 45 | mCD8 | VHHCD8 | VHH | mCD8 | 10 | QVQLQESGGGLVQAGGSLRLSCAASGFTFDDYAIGWFRQAPGKEREGVSCINNSDGSGYVADSVKGRFTISSDNAKNTVYLQMNSLKPEDTAVYYCAAAGDGGCYYGLDYWGKGTQVTVSS |
| 46 | hMHCII A | VHH4 | VHH | hMHCII | 12 | QVQLQESGGGLVQAGGSLRLSCAASGSTLSSYGMGWYRQAPGKQREVVATISATGSISYADSVKGRFTISRDSAKNTMYLQLNSLTPEDTAVYYCNTIYRSTLYWGQGTQVTVSS |
| 47 | hMHCII B | VHHDC13 | VHH | mMHCII | 13 | QVQLQESGGGLVQAGGSHNLSCTASGITFSSLAMGWFRQTPGKEREFVANIMRSGSSVFYADSVRGRFTISRDNAKNTA |

| | | | | | | |
|---|---|---|---|---|---|---|
| | | | | | | HLQMNSLKPEDTAVYFCAATRGAWPAEYWGQGTQVTVSS GGLPETGGHHHHHH |
| 48 | hTIM-3 | hH6 | VHH | hTIM-3 | 7 | QVQLVESGGGMVQPGDSLRLSCVASGRTGSSYIIGWFRQ APGKEREFVARVSPSGGTRDYADSVKGRFTVSRDNAKNT VYLQMDRLKPEDTAIYTCAAAGGKWTADSGEYNYWGQGT QVTVSS |
| 49 | hCTLA-4 | H11 | VHH | hCTLA-4 | 7 | QVQLQESGGGLAQPGGSLRLSCAASGSTISSVAVGWYRQ TPGNQREWVATSSTSSTTATYADSVKGRFTISRDNAKNTIY LQMNSLKPEDTAVYYCKTGLTNWGQGTQVTVSS |
| 50 | hCD8a A | R3HCD27 | VHH | hCD8a | 14 | QVQLQESGGGSVQPGGSLRLSCAASGFTFDDYAMSWVR QVPGKGLEWVSTINWNGGSAEYAEPVKGRFTISRDNAKNT VYLQMNSLKLEDTAVYYCAKDADLVWYNLSTGQGTQVTVS S |
| 51 | hCD8a B | R3HCD129 | VHH | hCD8a | 14 | QVQLQESGGGLVQPGGSLRLSCAASGFTFDDYAMSWVR QVPGKGLEWVSTINWNGGSAEYAEPVKGRFTISRDNAKNT VYLQMNSLKLEDTAVYYCAKDADLVWYNLRTGQGTQVTV SS |
| 52 | hCD8a C | R2HCD26 | VHH | hCD8a | 14 | QVQLQESGGGLVQAGGSLRLSCAASGFTFDDYAIGWFRQ APGKEREGVSCIRVSDGSTYYADPVKGRFTISSDNAKNTV YLQMNSLKPEDAAVYYCAAGSLYTCVQSIVVVPARPYYDM DYWGKGTQVTVSS |
| 53 | hCD4 | 03F11 | VHH | hCD4 | 15 | EVQLVESGGGSVQPGGSLTLSCGTSGRTFNVMGWFRQA PGKEREFVAAVRWSSTGIYYTQYADSVKSRFTISRDNAKN TVYLEMNSLKPEDTAVYYCAADTYNSNPARWDGYDFRGQ GTQVTVSS |
| 54 | hCXCR4 | 281F12 | VHH | hCXCR4 | 15 | EVQLVESGGGLVQAGDSLRLSCAASGRAFSRYAMGWFRQ APGKEREFVAAIGWGPSKTNYADSVKGRFTISRDNAKNTV YLQMNTLKPEDTAVYSCAAKFVNTDSTWSRSEMYTYWGQ GTQVTVSS |
| 55 | hCD38 A | MU551 | VHH | hCD38 | 16 | DVQLQESGGGLVQPGHSLRLSCVGSGSRFDNYAMGWFR QAPGKEREFVAAISWSSGTTRYLDTVKGRFTISRDNSKSTV YLQMNSLRAEDTAVYYCAARYQPRYYDSGDMDGYEYDN WGQGTQVTVSS |
| 56 | hCD38 B | MU375 | VHH | hCD38 | 16 | DVQLQESGGGSVQPGGSLTLSCTASGLLFRLASMGWYRQ APGKERELIATITVGGKTNYKDSVQGRFIITRDNTGDNTKST VTLQMNRLRAEDTAVYYCNTASPAVGADTWGQGTRVTVS S |
| 57 | hCD38 C | MU1053 | VHH | hCD38 | 16 | DVQLQESGGGLVQPGGSLRLSCTGSGRTFRNYPMAWFR QAPGKEREFVAGITWVGASTLYADFAKGRFTISRDNSKNT VYLQMNSLRAEDTAVYSCAAGRGIVAGRIPAEYADWGQGT QVTVSS |
| 58 | hCD7 A | VHH10 | VHH | hCD7 | 17 | MDVQLQESGGGSVQAGGSLRLSCAASGYTHSSYCMAWF RQAPGREREGVASIDSDGTTSYADSVKGRFTISQDNAKNT |

| | | | | | | |
|---|---|---|---|---|---|---|
| | | | | | | LYLQMNSLKPEDTAMYYCAARFGPMGCVDLSTLSFGHWGQGTQVTVSIT |
| 59 | hCD7 B | VHH6 | VHH | hCD7 | 17 | MDVQLQESGGGLVQAGGSLRLSCAVSGYPYSSYCMGWFRQAPGKEREGVAAIDSDGRTRYADSVKGRFTISQDNAKNTLYLQMNRMKPEDTAMYYCAARFGPMGCVDLSTLSFGHWGQGTQVTVSIT |
| 60 | hCD7 C | Humanized VHH10 | VHH | hCD7 | 17 | MDVQLQESGGGSVQPGGSLRLSCAASGYTHSSYCMAWFRQAPGREREGVASIDSDGTTSYADSVKGRFTISQDNSKNTLYLQMNSLRAEDTAMYYCAARFGPMGCVDLSTLSFGHWGQGTQVTVSIT |
| 61 | hCD7 D | Humanized VHH6 | VHH | hCD7 | 17 | MDVQLQESGGGLVQPGGSLRLSCAVSGYPYSSYCMGWFRQAPGKEREGVAAIDSDGRTRYADSVKGRFTISQDNSKNTLYLQMNRMRAEDTAMYYCAARFGPMGCVDLSTLSFGHWGQGTQVTVSIT |
| 62 | hCD7 E | Humanized Scaffold VHH10 | VHH | hCD7 | 17 | EVQLQASGGGFVQPGGSLRLSCAASGYTHSSYCMGWFRQAPGKEREFVSSIDSDGTTSYYADSVKGRFTISRDNSKNTVYLQMNSLRAEDTATYYCAARFGPMGCVDLSTLSFGHYWGQGTQVTVSS |
| 63 | hCD7 F | Humanized Scaffold VHH6 | VHH | hCD7 | 17 | QVQLQESGGGLVQPGGSLRLSCAASGYPYSSYCMGWYRQAPGKEREGVsAIDSDGRTRYYADSVKGRFTISRDNSKNTVYLQMNSLRAEDTAVYYCAARFGPMGCVDLSTLSFGHWGQGTQVTVSS |
