## Extended data table 2 for "*In vivo* generation of chimeric antigen receptor T cells using optimally retargeted and functionalized lentiviral vectors with reduced immune clearance"

| Donor ID | Sex | Age | MeV IgG score |
|---|---|---|---|
| 2352 | F | 67 | 4.56 |
| 0467 | M | 46 | 13.36 |
| 5932 | F | 62 | 14.79 |
| 0945 | F | 47 | 15.91 |
| 1445 | M | 52 | 18.93 |
| 1255 | F | 32 | 23.44 |
| 0470 | M | 54 | 23.70 |
| 3350 | F | 65 | 23.71 |
| 4266 | M | 53 | 24.17 |
| 2304 | M | 45 | 27.00 |
| 4610 | M | 43 | 27.99 |
| 0261 | F | 50 | 28.10 |
| 6201 | M | 36 | 34.40 |
| 9411 | F | 75 | 34.58 |
| 0389 | M | 82 | 37.34 |
| 1439 | F | 73 | 37.83 |
| 0288 | F | 49 | 38.74 |
| 2788 | M | 72 | 39.59 |
| 6526 | F | 13 | 41.48 |
| 2563 | M | 12 | 42.15 |
| 9642 | M | 62 | 48.28 |
