## Extended data table 3 for "*In vivo* generation of chimeric antigen receptor T cells using optimally retargeted and functionalized lentiviral vectors with reduced immune clearance"

| Condition | CD3-/CD45+ | CD3-/CD45+/CAR+ | CD3-/CD45- | CD3-/CD45-/CAR+ | CD3+ | CD3+/CAR+ |
|---|---|---|---|---|---|---|
| Nalm6 | 75.0 ± 43.1 | 0.3 ± 0.3 | 11266.3 ± 6614.2 | 2.0 ± 1.0 | 0.0 ± 0.0 | 0.0 ± 0.0 |
| Unmodified T cells | 77.7 ± 24.7 | 1.7 ± 1.2 | 19063.3 ± 4341.9 | 2.3 ± 0.9 | 42.0 ± 18.5 | 0.0 ± 0.0 |
| CAR T | 87.3 ± 20.1 | 3.0 ± 1.5 | 12327.0 ± 1261.3 | 2.7 ± 1.8 | 176.7 ± 28.3 | 61.3 ± 10.3 |
| In Vivo CAR T | 103.3 ± 46.0 | 0.3 ± 0.3 | 20850.0 ± 11697.0 | 1.7 ± 1.2 | 2126.0 ± 1330.8 | 519.7 ± 268.4 |
