## Extended data table 4 for "*In vivo* generation of chimeric antigen receptor T cells using optimally retargeted and functionalized lentiviral vectors with reduced immune clearance"

| Plasmid | Description | Sequence |
|---|---|---|
| pCG-MeV-F | Expression of WT measles virus Fusion protein with 30AA intravirion truncation | gaattcgagctcgccccgttacataacttacggtaaatggcccgcctggctgaccgcccaacgacccccgcccattgacgtcaataatgacgtatgttcccatagtaacgccaatagggactttccattgacgtcaatgggtggagtatttacggtaaactgcccacttggcagtacatcaagtgtatcatatgccaagtacgcccctattgacgtcaatgacgg<br>taaatggcccgcctggcattatgcccagtacatgaccttatgggactttcctacttggcagtacatctacgtattagtcatcgctattaccatggtgatgcggttttggcagtacatcaatgggcgtggatagcggtttgactcacggggatttccaagtctccaccccattgacgtcaatgggagtttgttttggcaccaaaatcaacgggactttccaaaatgtcgtaac<br>aactccgccccattgacgcaaatgggcggtaggcgtgtacggtgggaggtctatataagcagagctcgtttagtgaaccgtcagatcgcctggagacgccatccacgctgttttgacctccatagaagacaccgggaccgatccagcctccggggggatcgatcccccgatcctgagaacttcagggtgagtttggggacccttgattgttctttctttttcgctat<br>tgtaaaattcatgttatatggagggggcaaagttttcagggtgttgtttagaatgggaagatgtcccttgtatcaccatggaccctcatgataattttgtttctttcactttctactctgttgacaaccattgtctcctcttattttcttttcattttctgtaactttttcgttaaactttagcttgcatttgtaacgaattttttaaattcacttttgtttatttgtcagattgtaagtactttctctaatcac<br>ttttttttcaaggcaatcagggtatattatattgtacttcagcacagttttagagaacaattgttataattaaatgataaggtagaatatttctgcatataaattctggctggcgtggaaatattcttattggtagaaacaactacatcctggtcatcatcctgcctttctctttatggttacaatgatatacactgtttgagatgaggataaaatactctgagtccaaaccgggccc<br>ctctgctaaccatgttcatgccttcttctttttcctacagctcctgggcaacgtgctggttattgtgctgtctcatcattttggcaaagaattgtaatacgactcactatagggcgaattcggatcccccggtagttaattaaaaattagggtgcaagatcatcgataacaagtttgtacaaaaaagcaggcttcgccaccatgggcctgaaggtgaacgtgagcgccatct<br>tcatggccgtgctgctgaccctgcagacccccaacaggccagatccactggggcaatctgtctaagatcggagtggtgggaatcggatccgcctcttataaagtgatgacaaggagctcccaccagagcctggtcatcaagctgatgcctaacatcaccctgctgaacaattgtacacgcgtggagatcgccgagtaccggagactgctgaggaccgtgctg<br>gagccaatcagggacgccctgaacgccatgacacagaatatcaggccagtgcagagcgtggcatctagcaggcgccacaagagatttgccggcgtggtgctggcaggagccgccctgggagtggcaaccgccgcccagatcacagccggcatcgccctgcaccagtccatgctgaactctcaggccatcgataatctgagggcctccctggagac<br>aacaaaccaggcaatcgaggcaatcaggcaggcaggacaggagatgatcctggccgtgcagggcgtgcaggactatatcaacaatgagctgatcccttctatgaatcagctgagctgcgatctgatcggccagaagctgggcctgaagctgctgcggtactataccgagatcctgtccctgttcggaccttctctgcgggaccccatcagcgccgagatc<br>tccatccaggccctgtcctatgcactgggaggcgacatcaacaaggtgctggagaagctgggctactctggcggcgatctgctgggcatcctggagagccggggcatcaaggccagaatcacccacgtggatacagagtcttacttcatcgtgctgagcatcgcctaccccacactgtccgagatcaagggcgtgatcgtgcacaggctggagggcgtga<br>gctataatatcggctcccaggagtggtacaccacagtgcctaagtatgtggccacccagggctacctgatctctaacttcgacgagtcctcttgtacctttatgccagagggcacagtgtgctcccagaatgccctgtacccaatgtctcccctgctgcaggagtgtctgcggggctctaccaagagctgcgccagaacactggtgagcggctccttcggcaatc<br>ggttcatcctgtcccagggcaacctgatcgccaattgtgcctctatcctgtgcaagtgctataccacaggcaccatcatcaaccaggaccccgataagatcctgacatacatcgcagcagaccactgccctgtggtggaagtgaatggcgtgaccatccaagtgggcagccggagatacccagacgccgtgtatctgcacaggatcgatctgggaccacct<br>atctccctggagcgcctggatgtgggcacaaacctgggcaatgccatcgccaagctggaggacgccaaggagctgctggagagctccgatcagatcctgaggagcatgaagggcctgtctagcacctccatcgtgtacatcctgatcgccgtgtgcctgggaggcctgatcggaatccccgccctgatctgctgttgcaggggccgctgagacccagcttt<br>cttgtacaaagtggtactagtgtgaaatagacatcagaattaagaaaaacgtagggtccaagtggttccccgttatggactcgctatctgtcaaccagatcttataccctgaagttcacctagatagcccgatagttaccaataagatagtagccatcctggagtatgctcgactgtttaaacctgcaggcatgcaagctgatctttttccctctgccaaaaattatgg<br>ggacatcatgaagccccttgagcatctgacttctggctaataaaggaaatttattttcattgcaatagtgtgttggaattttttgtgtctctcactcggaaggacatatgggagggcaaatcatttaaaacatcagaatgagtatttggtttagagtttggcaacatatgcccatatgctggctgccatgaacaaaggttggctataaagaggtcatcagtatatgaaaca<br>gccccctgctgtccattccttattccatagaaaagccttgacttgaggttagattttttttatattttgttttgtgttatttttttctttaacatccctaaaattttccttacatgttttactagccagatttttcctcctctcctgactactcccagtcatagctgtccctcttctcttatggagatccctcgaggagcttttttgcaaaagcctaggcctccaaaaaagcctcttcactacttctgg<br>aatagctcagaggccgaggcggcctcggcctctgcataaataaaaaaaattagtcagccatggggcggagaatgggcggaactgggcggagttaggggcgggatgggcggagttaggggcgggactatggttgctgactaattgctgcattaatgaatcggccaacgcgcggggagaggcggtttgcgtattgggcgctcttccgcttcctcgctcactg<br>actcgctgcgctcggtcgttcggctgcggcgagcggtatcagctcactcaaaggcggtaatacggttatccacagaatcaggggataacgcaggaaagaacatgtgagcaaaaggccagcaaaaggccaggaaccgtaaaaaggccgcgttgctggcgtttttccataggctccgcccccctgacgagcatcacaaaaatcgacgctcaagtcagag<br>gtggcgaaacccgacaggactataaagataccaggcgtttccccctggaagctccctcgtgcgctctcctgttccgaccctgccgcttaccggatacctgtccgcctttctcccttcgggaagcgtggcgctttctcatagctcacgctgtaggtatctcagttcggtgtaggtcgttcgctccaagctgggctgtgtgcacgaaccccccgttcagcccgaccgctg<br>cgccttatccggtaactatcgtcttgagtccaacccggtaagacacgacttatcgccactggcagcagccactggtaacaggattagcagagcgaggtatgtaggcggtgctacagagttcttgaagtggtggcctaactacggctacactagaagaacagtatttggtatctgcgctctgctgaagccagttaccttcggaaaaagagttggtagctcttgatcc<br>ggcaaacaaaccaccgctggtagcggtggtttttttgtttgcaagcagcagattacgcgcagaaaaaaaggatctcaagaagatcctttgatcttttctacggggtctgacgctcagtggaacgaaaactcacgttaagggattttggtcatgagattatcaaaaaggatcttcacctagatccttttaaattaaaaatgaagttttaaatcaatctaaagtatatatga<br>gtaaacttggtctgacagttaccaatgcttaatcagtgaggcacctatctcagcgatctgtctatttcgttcatccatagttgcctgactccccgtcgtgtagataactacgatacgggagggcttaccatctggccccagtgctgcaatgataccgcgagacccacgctcaccggctccagatttatcagcaataaaccagccagccggaagggccgagcgcag<br>aagtggtcctgcaactttatccgcctccatccagtctattaattgttgccgggaagctagagtaagtagttcgccagttaatagtttgcgcaacgttgttgccattgctacaggcatcgtggtgtcacgctcgtcgtttggtatggcttcattcagctccggttcccaacgatcaaggcgagttacatgatcccccatgttgtgcaaaaaagcggttagctccttcggtcctc<br>cgatcgttgtcagaagtaagttggccgcagtgttatcactcatggttatggcagcactgcataattctcttactgtcatgccatccgtaagatgcttttctgtgactggtgagtactcaaccaagtcattctgagaatagtgtatgcggcgaccgagttgctcttgcccggcgtcaatacgggataataccgcgccacatagcagaactttaaaagtgctcatcattggaa<br>aacgttcttcggggcgaaaactctcaaggatcttaccgctgttgagatccagttcgatgtaacccactcgtgcacccaactgatcttcagcatcttttactttcaccagcgtttctgggtgagcaaaaacaggaaggcaaaatgccgcaaaaaagggaataagggcgacacggaaatgttgaatactcatactcttcctttttcaatattattgaagcatttatcagg<br>gttattgtctcatgagcggatacatatttgaatgtatttagaaaaataaacaaataggggttccgcgcacatttccccgaaaagtgccacctgacgtctaagaaaccattattatcatgacattaacctataaaaataggcgtatcacgaggccctttcgtctcgcgcgtttcggtgatgacggtgaaaacctctgacacatgcagctcccggagacggtcacagctt<br>gtctgtaagcggatgccgggagcagacaagcccgtcagggcgcgtcagcgggtgttggcgggtgtcggggctggcttaactatgcggcatcagagcagattgtactgagagtgcaccattcgacgctctcccttatgcgactcctgcattaggaagcagcccagtagtaggttgaggccgttgagcaccgccgccgcaaggaatggtctggcttatcgaaa<br>ttaatacgactcactatagggagaccg |
| pCG-MeV-H | Expression of WT measles virus Hemagluttinin protein with 18AA intravirion truncation and N481A mutation | gaattcgagctcgccccgttacataacttacggtaaatggcccgcctggctgaccgcccaacgacccccgcccattgacgtcaataatgacgtatgttcccatagtaacgccaatagggactttccattgacgtcaatgggtggagtatttacggtaaactgcccacttggcagtacatcaagtgtatcatatgccaagtacgcccctattgacgtcaatgacgg<br>taaatggcccgcctggcattatgcccagtacatgaccttatgggactttcctacttggcagtacatctacgtattagtcatcgctattaccatggtgatgcggttttggcagtacatcaatgggcgtggatagcggtttgactcacggggatttccaagtctccaccccattgacgtcaatgggagtttgttttggcaccaaaatcaacgggactttccaaaatgtcgtaac<br>aactccgccccattgacgcaaatgggcggtaggcgtgtacggtgggaggtctatataagcagagctcgtttagtgaaccgtcagatcgcctggagacgccatccacgctgttttgacctccatagaagacaccgggaccgatccagcctccggggggatcgatcccccgatcctgagaacttcagggtgagtttggggacccttgattgttctttctttttcgctat<br>tgtaaaattcatgttatatggagggggcaaagttttcagggtgttgtttagaatgggaagatgtcccttgtatcaccatggaccctcatgataattttgtttctttcactttctactctgttgacaaccattgtctcctcttattttcttttcattttctgtaactttttcgttaaactttagcttgcatttgtaacgaattttttaaattcacttttgtttatttgtcagattgtaagtactttctctaatcac<br>ttttttttcaaggcaatcagggtatattatattgtacttcagcacagttttagagaacaattgttataattaaatgataaggtagaatatttctgcatataaattctggctggcgtggaaatattcttattggtagaaacaactacatcctggtcatcatcctgcctttctctttatggttacaatgatatacactgtttgagatgaggataaaatactctgagtccaaaccgggccc<br>ctctgctaaccatgttcatgccttcttctttttcctacagctcctgggcaacgtgctggttattgtgctgtctcatcattttggcaaagaattgtaatacgactcactatagggcgaattcggatcccccggtagttaattaaaaattagggtgcaagatcatcgataacaagtttgtacaaaaaagcaggcttcgccaccatgggctctaggatcgtgatcaaccgcgagc<br>acctgatgatcgacaggccatacgtgctgctggccgtgctgttcgtgatgtttctgagcctgatcggcctgctggcaatcgcaggaatcaggctgcacagagccgccatctataccgccgagatccacaagagcctgtccaccaacctggacgtgacaaattccatcgagcaccaggtgaaggatgtgctgacaccactgttcaagatcatcggcgacgaa<br>gtgggcctgaggaccccacagcggttcaccgacctggtgaagttcatcagcgataagatcaagtttctgaaccctgatcgggagtacgacttcagagatctgacctggtgcatcaatcccccctgagcggatcaagctggactacgatcagtattgtgccgacgtggccgccgaggagctgatgaacgccctggtgaattccaccctgctggaggccagagc<br>cacaaaccagtttctggccgtgagcaagggcaattgctccggcccaaccacaatccggggccagttctctaacatgtctctgagcctgctggatctgtacctgagcagaggctataatgtgagctccatcgtgaccatgacaagccagggcatgtacggcggcacctatctggtgggcaagcccaacctgtctagcaagggctccgagctgtctcagctgag<br>catgcacagggtgtttgaagtgggcgtgatcaggaacccaggcctgggagccccgtgtttcacatgaccaattacttcgagcagcccgtgagcaacgatttttctaattgcatggtggccctgggagagctgaagttcgccgccctgtgccacagggaggacagcatcaccatcccatatcagggctctggcaagggcgtgagcttccagctggtgaagct<br>gggcgtgtggaagtcccccaccgacatgagatcctgggtgcctctgtctacagacgatccagtgatcgatcggctgtacctgtcctctcacagaggcgtgatcgccgacaatcaggcaaagtgggcagtgccaaccacaaggaccgacgataagctgagaatggagacatgctttcagcaggcctgtaagggcaagaaccaggccctgtgcgagaatc<br>cagagtgggcacctctgaaggataacagaatccctagctacggcgtgctgagcgtgaatctgtccctgaccgtggagctgaagatcaagatcgccagcggcttcggcccactgatcacacacggctccggcatggacctgtacaagaccaaccacaacaacgtgtactggctgacaatcccacccatgaagaacctggccctgggcgtgatcaatacc<br>ctggagtggatccccaggtttaaggtgtcccctgccctgttcaccgtgccaatcaaggaggccggcgaggattgtcacgccccaacatacctgcccgccgaggtggacggcgatgtgaagctgagctccaacctggtcatcctgcctggccaggacctgcagtacgtgctggccacctatgatacaagccgcgtggagcacgccgtggtgtactacgtgta<br>ctcccctctcggagcttctcctacttttatcctttcagactgccaatcaagggcgtgcctatcgagctgcaggtggagtgctttacctgggataagaagctgtggtgcaggcacttctgcgtgctggcagactctgagagcggaggacacatcacccactccggaatggtgggcatgggcgtgtcttgtaccgtgacacgcgaggacggcacatgagacccag<br>ctttcttgtacaaagtggtggctgctagtgaaccaatcacatgatgtcacccagacatcaggcatacccactagtgtgaaatagacatcagaattaagaaaaacgtagggtccaagtggttccccgttatggactcgctatctgtcaaccagatcttataccctgaagttcacctagatagcccgatagttaccaataagatagtagccatcctggagtatgctcga<br>ctgtttaaacctgcaggcatgcaagctgatctttttccctctgccaaaaattatggggacatcatgaagccccttgagcatctgacttctggctaataaaggaaatttattttcattgcaatagtgtgttggaattttttgtgtctctcactcggaaggacatatgggagggcaaatcatttaaaacatcagaatgagtatttggtttagagtttggcaacatatgcccatatgct<br>ggctgccatgaacaaaggttggctataaagaggtcatcagtatatgaaacagccccctgctgtccattccttattccatagaaaagccttgacttgaggttagattttttttatattttgttttgtgttatttttttctttaacatccctaaaattttccttacatgttttactagccagatttttcctcctctcctgactactcccagtcatagctgtccctcttctcttatggagatccctcga<br>ggagctttttgcaaaagcctaggcctccaaaaaagcctcttcactacttctggaatagctcagaggccgaggcggcctcggcctctgcataaataaaaaaaattagtcagccatggggcggagaatgggcggaactgggcggagttaggggcgggatgggcggagttaggggcgggactatggttgctgactaattgctgcattaatgaatcggccaacg<br>cgcggggagaggcggtttgcgtattgggcgctcttccgcttcctcgctcactgactcgctgcgctcggtcgttcggctgcggcgagcggtatcagctcactcaaaggcggtaatacggttatccacagaatcaggggataacgcaggaaagaacatgtgagcaaaaggccagcaaaaggccaggaaccgtaaaaaggccgcgttgctggcgtttttccat<br>aggctccgcccccctgacgagcatcacaaaaatcgacgctcaagtcagaggtggcgaaacccgacaggactataaagataccaggcgtttccccctggaagctccctcgtgcgctctcctgttccgaccctgccgcttaccggatacctgtccgcctttctcccttcgggaagcgtggcgctttctcatagctcacgctgtaggtatctcagttcggtgtaggtcg<br>ttcgctccaagctgggctgtgtgcacgaaccccccgttcagcccgaccgctgcgccttatccggtaactatcgtcttgagtccaacccggtaagacacgacttatcgccactggcagcagccactggtaacaggattagcagagcgaggtatgtaggcggtgctacagagttcttgaagtggtggcctaactacggctacactagaagaacagtatttggtatc<br>tgcgctctgctgaagccagttaccttcggaaaaagagttggtagctcttgatccggcaaacaaaccaccgctggtagcggtggtttttttgtttgcaagcagcagattacgcgcagaaaaaaaggatctcaagaagatcctttgatcttttctacggggtctgacgctcagtggaacgaaaactcacgttaagggattttggtcatgagattatcaaaaaggatcttc<br>acctagatccttttaaattaaaaatgaagttttaaatcaatctaaagtatatatgagtaaacttggtctgacagttaccaatgcttaatcagtgaggcacctatctcagcgatctgtctatttcgttcatccatagttgcctgactccccgtcgtgtagataactacgatacgggagggcttaccatctggccccagtgctgcaatgataccgcgagacccacgctcaccg<br>gctccagatttatcagcaataaaccagccagccggaagggccgagcgcagaagtggtcctgcaactttatccgcctccatccagtctattaattgttgccgggaagctagagtaagtagttcgccagttaatagtttgcgcaacgttgttgccattgctacaggcatcgtggtgtcacgctcgtcgtttggtatggcttcattcagctccggttcccaacgatcaaggc<br>gagttacatgatcccccatgttgtgcaaaaaagcggttagctccttcggtcctccgatcgttgtcagaagtaagttggccgcagtgttatcactcatggttatggcagcactgcataattctcttactgtcatgccatccgtaagatgcttttctgtgactggtgagtactcaaccaagtcattctgagaatagtgtatgcggcgaccgagttgctcttgcccggcgtcaata<br>cgggataataccgcgccacatagcagaactttaaaagtgctcatcattggaaaacgttcttcggggcgaaaactctcaaggatcttaccgctgttgagatccagttcgatgtaacccactcgtgcacccaactgatcttcagcatcttttactttcaccagcgtttctgggtgagcaaaaacaggaaggcaaaatgccgcaaaaaagggaataagggcgacac<br>ggaaatgttgaatactcatactcttcctttttcaatattattgaagcatttatcagggttattgtctcatgagcggatacatatttgaatgtatttagaaaaataaacaaataggggttccgcgcacatttccccgaaaagtgccacctgacgtctaagaaaccattattatcatgacattaacctataaaaataggcgtatcacgaggccctttcgtctcgcgcgtttcggtg<br>atgacggtgaaaacctctgacacatgcagctcccggagacggtcacagcttgtctgtaagcggatgccgggagcagacaagcccgtcagggcgcgtcagcgggtgttggcgggtgtcggggctggcttaactatgcggcatcagagcagattgtactgagagtgcaccattcgacgctctcccttatgcgactcctgcattaggaagcagcccagtagtag<br>gttgaggccgttgagcaccgccgccgcaaggaatggtctggcttatcgaaattaatacgactcactatagggagaccg |

| | | |
|---|---|---|
| pCG-MeV-H-hCD105scFv-6His | Expression of pCG-MeV-H with the human CD105 targeting scFv and 6His tag at the C-terminus | gaattcgagctcgccccgttacataacttacggtaaatggcccgcctggctgaccgcccaacgacccccgcccattgacgtcaataatgacgtatgttcccatagtaacgccaatagggactttccattgacgtcaatgggtggagtatttacggtaaactgcccacttggcagtacatcaagtgtatcatatgccaagtacgccccctattgacgtcaatgacgg taaatggcccgcctggcattatgcccagtacatgaccttatgggactttcctacttggcagtacatctacgtattagtcatcgctattaccatggtgatgcggttttggcagtacatcaatgggcgtggatagcggtttgactcacggggatttccaagtctccaccccattgacgtcaatgggagtttgttttggcaccaaaatcaacgggactttccaaaatgtcgtaac aactccgccccattgacgcaaatgggcggtaggcgtgtacggtgggaggtctatataagcagagctcgtttagtgaaccgtcagatcgcctggagacgccatccacgctgttttgacctccatagaagacaccgggaccgatccagcctccggggggatcgatcccccgatcctgagaacttcagggtgagtttggggacccttgattgttctttctttttcgctat tgtaaaattcatgttatatggagggggcaaagttttcagggtgttgtttagaatgggaagatgtcccttgtatcaccatggaccctcatgataattttgtttctttcactttctactctgttgacaaccattgtctcctcttattttcttttcattttctgtaactttttcgttaaactttagcttgcatttgtaacgaatttttaaattcacttttgtttatttgtcagattgtaagtactttctctaatcac tttttttcaaggcaatcagggtatattatattgtacttcagcacagttttagagaacaattgttataattaaatgataaggtagaatatttctgcatataaattctggctggcgtggaaatattcttattggtagaaacaactacatcctggtcatcatcctgcctttctctttatggttacaatgatatacactgtttgagatgaggataaaatactctgagtccaaaccgggccc ctctgctaaccatgttcatgccttcttctttttcctacagctcctgggcaacgtgctggttattgtgctgtctcatcattttggcaaagaattgtaatacgactcactatagggcgaattcggatcccccggtagttaattaaaaattagggtgcaagatcatcgataacaagtttgtacaaaaaagcaggcttcgccaccatgggctctaggatcgtgatcaaccgcgagc acctgatgatcgacaggccatacgtgctgctggccgtgctgttcgtgatgtttctgagcctgatcggcctgctggcaatcgcaggaatcaggctgcacagagccgccatctataccgccgagatccacaagagcctgtccaccaacctggacgtgacaaattccatcgagcaccaggtgaaggatgtgctgacaccactgttcaagatcatcggcgacgaa gtgggcctgaggaccccacagcggttcaccgacctggtgaagttcatcagcgataagatcaagtttctgaaccctgatcgggagtacgacttcagagatctgacctggtgcatcaatcccccctgagcggatcaagctggactacgatcagtattgtgccgacgtggccgccgaggagctgatgaacgccctggtgaattccaccctgctggaggccagagc cacaaaccagtttctggccgtgagcaagggcaattgctccggcccaaccacaatccggggccagttctctaacatgtctctgagcctgctggatctgtacctgagcagaggctataatgtgagctccatcgtgaccatgacaagccagggcatgtacggcggcacctatctggtgggcaagcccaacctgtctagcaagggctccgagctgtctcagctgag catgcacagggtgtttgaagtgggcgtgatcaggaacccaggcctgggagcccccgtgtttcacatgaccaattacttcgagcagcccgtgagcaacgatttttctaattgcatggtggccctgggagagctgaagttcgccgccctgtgccacagggaggacagcatcaccatcccatatcaggctctggcaagggcgtgagcttccagctggtgaagct gggcgtgtggaagtcccccaccgacatgagatcctgggtgcctctgtctacagacgatccagtgatcgatcggctgtacctgtcctctcacagaggcgtgatcgccgacaatcaggcaaagtgggcagtgccaaccacaaggaccgacgataagctgagaatggagacatgctttcagcaggcctgtaagggcaagaaccaggccctgtgcgagaatc cagagtgggcacctctgaaggataacagaatccctagctacggcgtgctgagcgtgaatctgtccctgaccgtggagctgaagatcaagatcgccagcggcttcggcccactgatcacacacggctccggcatggacctgtacaagaccaaccacaacaacgtgtactggctgacaatcccacccatgaagaacctggccctgggcgtgatcaatacc ctggagtggatccccaggtttaaggtgtcccctgccctgttcaccgtgccaatcaaggaggccggcgaggattgtcacgccccaacatacctgcccgccgaggtggacggcgatgtgaagctgagctccaacctggtcatcctgcctggccaggacctgcagtacgtgctggccacctatgatacaagccgcgtggagcacgccgtggtgtactacgtgta ctccccctctcggagcttctcctacttttatcctttcagactgccaatcaagggcgtgcctatcgagctgcaggtggagtgctttacctgggataagaagctgtggtgcaggcacttctgcgtgctggcagactctgagagcggaggacacatcacccactccggaatggtgggcatgggcgtgtcttgtaccgtgacacgcgaggacggcacaaacggagga ggaggaagcggaggaggaggatccggcggcggcggctctgccgcccagcctgccatggccgaggtgcagctgctggagtccggaggaggcctggtgcagccaggaggctctctgaggctgagctgcgcagcatccggcttcacctttctagctacgccatgagctgggtgcgccaggcaccaggcaagggcctggagtgggtgagcgccatctac ggatccgacggcgataccacatatgccgattccgtgaagggcaggttcaccatctctcgcgacaacagcaagaatacactgtatctgcagatgaactctctgagggccgaggataccgccgtgtactattgcgcccgcgtgttttacacagccggcttcgactattggggccagggcaccctggtgacagtgtcctctggctccgattctaacgcaggacacg catctgccggaaataccagcgacatcgagctgacacagtccccaagctccctgagcgcctccgtgggcgacagggtgaccatcacatgtcgcgcctctcagagcatctctagctccctgaattggtaccagcagaagcccggcaaggcccctaagctgctgatctatgcagcatctagcctgcagtccggagtgccatctaggttttccggctctggcagcg gcaccgatttcaccctgacaatctcctctctgcagcccgaggactttgccacatactattgtcagcaggcccctgccaagcctccaaccttcggccagggcacaaagctggagatcaagagggcagcagcaagaggcagccaccaccaccaccaccactgagacccagctttcttgtacaaagtggtggctgctagtgaaccaatcacatgatgtcaccc agacatcaggcatacccactagtgtgaaatagacatcagaattaagaaaaacgtagggtccaagtggttccccgttatggactcgctatctgtcaaccagatcttatacctgaagttcacctagatagcccgatagttaccaataagatagtagccatcctggagtatgctcgactgtttaaacctgcaggcatgcaagctgatctttttccctctgccaaaaatta tggggacatcatgaagccccttgagcatctgacttctggctaataaaggaaatttattttcattgcaatagtgtgttggaattttttgtgtctctcactcggaaggacatatgggagggcaaatcatttaaaacatcagaatgagtatttggtttagagtttggcaacatatgcccatatgctggctgccatgaacaaaggttggctataaagaggtcatcagtatatgaaa cagccccctgctgtccattccttattccatagaaaagccttgacttgaggttagattttttttatattttgttttgtgttatttttttctttaacatccctaaaattttccttacatgttttactagccagatttttcctcctctcctgactactcccagtcatagctgtccctcttctcttatggagatccctcgaggagctttttgcaaaagcctaggcctccaaaaaagcctcttcactacttct ggaatagctcagaggccgaggcggcctcggcctctgcataaataaaaaaaattagtcagccatggggcggagaatgggcggaactgggcggagttaggggcgggatgggcggagttaggggcgggactatggttgctgactaattgctgcattaatgaatcggccaacgcgcggggagaggcggtttgcgtattgggcgctcttccgcttcctcgctca ctgactcgctgcgctcggtcgttcggctgcggcgagcggtatcagctcactcaaaggcggtaatacggttatccacagaatcaggggataacgcaggaaagaacatgtgagcaaaaggccagcaaaaggccaggaaccgtaaaaaggccgcgttgctggcgtttttccataggctccgcccccctgacgagcatcacaaaaatcgacgctcaagtca gaggtggcgaaacccgacaggactataaagataccaggcgtttccccctggaagctccctcgtgcgctctcctgttccgaccctgccgcttaccggatacctgtccgcctttctcccttcgggaagcgtggcgctttctcatagctcacgctgtaggtatctcagttcggtgtaggtcgttcgctccaagctgggctgtgtgcacgaaccccccgttcagcccgaccg ctgcgccttatccggtaactatcgtcttgagtccaacccggtaagacacgacttatcgccactggcagcagccactggtaacaggattagcagagcgaggtatgtaggcggtgctacagagttcttgaagtggtggcctaactacggctacactagaagaacagtatttggtatctgcgctctgctgaagccagttaccttcggaaaaagagttggtagctcttga tccggcaaacaaaccaccgctggtagcggtggtttttttgtttgcaagcagcagattacgcgcagaaaaaaaggatctcaagaagatcctttgatcttttctacggggtctgacgctcagtggaacgaaaactcacgttaagggattttggtcatgagattatcaaaaaggatcttcacctagatccttttaaattaaaaatgaagttttaaatcaatctaaagtatata tgagtaaacttggtctgacagttaccaatgcttaatcagtgaggcacctatctcagcgatctgtctatttcgttcatccatagttgcctgactccccgtcgtgtagataactacgatacgggagggcttaccatctggccccagtgctgcaatgataccgcgagacccacgctcaccggctccagatttatcagcaataaaccagccagccggaagggccgagcg cagaagtggtcctgcaactttatccgcctccatccagtctattaattgttgccgggaagctagagtaagtagttcgccagttaatagtttgcgcaacgttgttgccattgctacaggcatcgtggtgtcacgctcgtcgtttggtatggcttcattcagctccggttcccaacgatcaaggcgagttacatgatcccccatgttgtgcaaaaaagcggttagctccttcggtc ctccgatcgttgtcagaagtaagttggccgcagtgttatcactcatggttatggcagcactgcataattctcttactgtcatgccatccgtaagatgcttttctgtgactggtgagtactcaaccaagtcattctgagaatagtgtatgcggcgaccgagttgctcttgcccggcgtcaatacgggataataccgcgccacatagcagaactttaaaagtgctcatcattg gaaaacgttcttcggggcgaaaactctcaaggatcttaccgctgttgagatccagttcgatgtaacccactcgtgcacccaactgatcttcagcatcttttactttcaccagcgtttctgggtgagcaaaaacaggaaggcaaaatgccgcaaaaaagggaataagggcgacacggaaatgttgaatactcatactcttcctttttcaatattattgaagcatttatc agggttattgtctcatgagcggatacatatttgaatgtatttagaaaaataaacaaataggggttccgcgcacatttccccgaaaagtgccacctgacgtctaagaaaccattattatcatgacattaacctataaaaataggcgtatcacgaggccctttcgtctcgcgcgtttcggtgatgacggtgaaaacctctgacacatgcagctcccggagacggtcac agcttgtctgtaagcggatgccgggagcagacaagcccgtcagggcgcgtcagcgggtgttggcgggtgtcggggctggcttaactatgcggcatcagagcagattgtactgagagtgcaccattcgacgctctcccttatgcgactcctgcattaggaagcagcccagtagtaggttgaggccgttgagcaccgccgccgcaaggaatggtctggcttatc gaaattaatacgactcactatagggagaccg |
| pCG-MeV-H-mMHCIIVHH-HA | Expression of pCG-MeV-H with the mouse MHCII targeting VHH clone B and HA tag at the C-terminus | gaattcgagctcgccccgttacataacttacggtaaatggcccgcctggctgaccgcccaacgacccccgcccattgacgtcaataatgacgtatgttcccatagtaacgccaatagggactttccattgacgtcaatgggtggagtatttacggtaaactgcccacttggcagtacatcaagtgtatcatatgccaagtacgccccctattgacgtcaatgacgg taaatggcccgcctggcattatgcccagtacatgaccttatgggactttcctacttggcagtacatctacgtattagtcatcgctattaccatggtgatgcggttttggcagtacatcaatgggcgtggatagcggtttgactcacggggatttccaagtctccaccccattgacgtcaatgggagtttgttttggcaccaaaatcaacgggactttccaaaatgtcgtaac aactccgccccattgacgcaaatgggcggtaggcgtgtacggtgggaggtctatataagcagagctcgtttagtgaaccgtcagatcgcctggagacgccatccacgctgttttgacctccatagaagacaccgggaccgatccagcctccggggggatcgatcccccgatcctgagaacttcagggtgagtttggggacccttgattgttctttctttttcgctat tgtaaaattcatgttatatggagggggcaaagttttcagggtgttgtttagaatgggaagatgtcccttgtatcaccatggaccctcatgataattttgtttctttcactttctactctgttgacaaccattgtctcctcttattttcttttcattttctgtaactttttcgttaaactttagcttgcatttgtaacgaatttttaaattcacttttgtttatttgtcagattgtaagtactttctctaatcac tttttttcaaggcaatcagggtatattatattgtacttcagcacagttttagagaacaattgttataattaaatgataaggtagaatatttctgcatataaattctggctggcgtggaaatattcttattggtagaaacaactacatcctggtcatcatcctgcctttctctttatggttacaatgatatacactgtttgagatgaggataaaatactctgagtccaaaccgggccc ctctgctaaccatgttcatgccttcttctttttcctacagctcctgggcaacgtgctggttattgtgctgtctcatcattttggcaaagaattgtaatacgactcactatagggcgaattcggatcccccggtagttaattaaaaattagggtgcaagatcatcgataacaagtttgtacaaaaaagcaggcttcgccaccatgggctctaggatcgtgatcaaccgcgagc acctgatgatcgacaggccatacgtgctgctggccgtgctgttcgtgatgtttctgagcctgatcggcctgctggcaatcgcaggaatcaggctgcacagagccgccatctataccgccgagatccacaagagcctgtccaccaacctggacgtgacaaattccatcgagcaccaggtgaaggatgtgctgacaccactgttcaagatcatcggcgacgaa gtgggcctgaggaccccacagcggttcaccgacctggtgaagttcatcagcgataagatcaagtttctgaaccctgatcgggagtacgacttcagagatctgacctggtgcatcaatcccccctgagcggatcaagctggactacgatcagtattgtgccgacgtggccgccgaggagctgatgaacgccctggtgaattccaccctgctggaggccagagc cacaaaccagtttctggccgtgagcaagggcaattgctccggcccaaccacaatccggggccagttctctaacatgtctctgagcctgctggatctgtacctgagcagaggctataatgtgagctccatcgtgaccatgacaagccagggcatgtacggcggcacctatctggtgggcaagcccaacctgtctagcaagggctccgagctgtctcagctgag catgcacagggtgtttgaagtgggcgtgatcaggaacccaggcctgggagcccccgtgtttcacatgaccaattacttcgagcagcccgtgagcaacgatttttctaattgcatggtggccctgggagagctgaagttcgccgccctgtgccacagggaggacagcatcaccatcccatatcaggctctggcaagggcgtgagcttccagctggtgaagct gggcgtgtggaagtcccccaccgacatgagatcctgggtgcctctgtctacagacgatccagtgatcgatcggctgtacctgtcctctcacagaggcgtgatcgccgacaatcaggcaaagtgggcagtgccaaccacaaggaccgacgataagctgagaatggagacatgctttcagcaggcctgtaagggcaagaaccaggccctgtgcgagaatc cagagtgggcacctctgaaggataacagaatccctagctacggcgtgctgagcgtgaatctgtccctgaccgtggagctgaagatcaagatcgccagcggcttcggcccactgatcacacacggctccggcatggacctgtacaagaccaaccacaacaacgtgtactggctgacaatcccacccatgaagaacctggccctgggcgtgatcaatacc ctggagtggatccccaggtttaaggtgtcccctgccctgttcaccgtgccaatcaaggaggccggcgaggattgtcacgccccaacatacctgcccgccgaggtggacggcgatgtgaagctgagctccaacctggtcatcctgcctggccaggacctgcagtacgtgctggccacctatgatacaagccgcgtggagcacgccgtggtgtactacgtgta ctccccctctcggagcttctcctacttttatcctttcagactgccaatcaagggcgtgcctatcgagctgcaggtggagtgctttacctgggataagaagctgtggtgcaggcacttctgcgtgctggcagactctgagagcggaggacacatcacccactccggaatggtgggcatgggcgtgtcttgtaccgtgacacgcgaggacggcacaggggaggc ggctctggaggtggggggtccggtggcggcggaagcgcggccgcgcaggttcaactggtgcagagcggaggcggcctggtccagcccggcggaagcctgggcctgagctgtgccgcttctggcaacatcggcagcagagataatatgggctggtatagacaggcccctggaaaacagagagagtgggtggctacaatcagcggctacggcatcg ccacataccgggacagcgtgaagggcagattcaccgtggccaaggacaccgccaagaacatcgtgtccctgcagatgaactacctgaccaccgaagatacagccgtgtactactgctacgcctacgccgtggacagccggaacatcttctggtcccagggcacccaggtgaccgtgtctggcggcggctccggcggcggcagctatccctatgatgtg cccgactatgcgtgaactagtgtgaaatagacatcagaattaagaaaaacgtagggtccaagtggttccccgttatggactcgctatctgtcaaccagatcttatacctgaagttcacctagatagcccgatagttaccaataagatagtagccatcctggagtatgctcgactgtttaaacctgcaggcatgcaagctgatctttttccctctgccaaaaattatgg ggacatcatgaagccccttgagcatctgacttctggctaataaaggaaatttattttcattgcaatagtgtgttggaattttttgtgtctctcactcggaaggacatatgggagggcaaatcatttaaaacatcagaatgagtatttggtttagagtttggcaacatatgcccatatgctggctgccatgaacaaaggttggctataaagaggtcatcagtatatgaaaca gccccctgctgtccattccttattccatagaaaagccttgacttgaggttagattttttttatattttgttttgtgttatttttttctttaacatccctaaaattttccttacatgttttactagccagatttttcctcctctcctgactactcccagtcatagctgtccctcttctcttatggagatccctcgaggagctttttgcaaaagcctaggcctccaaaaaagcctcttcactacttctgg aatagctcagaggccgaggcggcctcggcctctgcataaataaaaaaaattagtcagccatggggcggagaatgggcggaactgggcggagttaggggcgggatgggcggagttaggggcgggactatggttgctgactaattgctgcattaatgaatcggccaacgcgcggggagaggcggtttgcgtattgggcgctcttccgcttcctcgctcactg actcgctgcgctcggtcgttcggctgcggcgagcggtatcagctcactcaaaggcggtaatacggttatccacagaatcaggggataacgcaggaaagaacatgtgagcaaaaggccagcaaaaggccaggaaccgtaaaaaggccgcgttgctggcgtttttccataggctccgcccccctgacgagcatcacaaaaatcgacgctcaagtcagag gtggcgaaacccgacaggactataaagataccaggcgtttccccctggaagctccctcgtgcgctctcctgttccgaccctgccgcttaccggatacctgtccgcctttctcccttcgggaagcgtggcgctttctcatagctcacgctgtaggtatctcagttcggtgtaggtcgttcgctccaagctgggctgtgtgcacgaaccccccgttcagcccgaccgctg cgccttatccggtaactatcgtcttgagtccaacccggtaagacacgacttatcgccactggcagcagccactggtaacaggattagcagagcgaggtatgtaggcggtgctacagagttcttgaagtggtggcctaactacggctacactagaagaacagtatttggtatctgcgctctgctgaagccagttaccttcggaaaaagagttggtagctcttgatcc ggcaaacaaaccaccgctggtagcggtggtttttttgtttgcaagcagcagattacgcgcagaaaaaaaggatctcaagaagatcctttgatcttttctacggggtctgacgctcagtggaacgaaaactcacgttaagggattttggtcatgagattatcaaaaaggatcttcacctagatccttttaaattaaaaatgaagttttaaatcaatctaaagtatatatga gtaaacttggtctgacagttaccaatgcttaatcagtgaggcacctatctcagcgatctgtctatttcgttcatccatagttgcctgactccccgtcgtgtagataactacgatacgggagggcttaccatctggccccagtgctgcaatgataccgcgagacccacgctcaccggctccagatttatcagcaataaaccagccagccggaagggccgagcgcag aagtggtcctgcaactttatccgcctccatccagtctattaattgttgccgggaagctagagtaagtagttcgccagttaatagtttgcgcaacgttgttgccattgctacaggcatcgtggtgtcacgctcgtcgtttggtatggcttcattcagctccggttcccaacgatcaaggcgagttacatgatcccccatgttgtgcaaaaaagcggttagctccttcggtcctc cgatcgttgtcagaagtaagttggccgcagtgttatcactcatggttatggcagcactgcataattctcttactgtcatgccatccgtaagatgcttttctgtgactggtgagtactcaaccaagtcattctgagaatagtgtatgcggcgaccgagttgctcttgcccggcgtcaatacgggataataccgcgccacatagcagaactttaaaagtgctcatcattggaa aacgttcttcggggcgaaaactctcaaggatcttaccgctgttgagatccagttcgatgtaacccactcgtgcacccaactgatcttcagcatcttttactttcaccagcgtttctgggtgagcaaaaacaggaaggcaaaatgccgcaaaaaagggaataagggcgacacggaaatgttgaatactcatactcttcctttttcaatattattgaagcatttatcagg gttattgtctcatgagcggatacatatttgaatgtatttagaaaaataaacaaataggggttccgcgcacatttccccgaaaagtgccacctgacgtctaagaaaccattattatcatgacattaacctataaaaataggcgtatcacgaggccctttcgtctcgcgcgtttcggtgatgacggtgaaaacctctgacacatgcagctcccggagacggtcacagctt gtctgtaagcggatgccgggagcagacaagcccgtcagggcgcgtcagcgggtgttggcgggtgtcggggctggcttaactatgcggcatcagagcagattgtactgagagtgcaccattcgacgctctcccttatgcgactcctgcattaggaagcagcccagtagtaggttgaggccgttgagcaccgccgccgcaaggaatggtctggcttatcgaaa ttaatacgactcactatagggagaccg |

| | | |
|---|---|---|
| pCG-MeV-H-hCD7VHH | Expression of pCG-MeV-H with human CD7 targeting VHH clone VHH10 humanized and HA tag at the C-terminus | gaattcgagctcgccccgttacataacttacggtaaatggcccgcctggctgaccgcccaacgacccccgcccattgacgtcaataatgacgtatgttcccatagtaacgccaatagggactttccattgacgtcaatgggtggagtatttacggtaaactgcccacttggcagtacatcaagtgtatcatatgccaagtacgccccctattgacgtcaatgacgg<br>taaatggcccgcctggcattatgcccagtacatgaccttatgggactttcctacttggcagtacatctacgtattagtcatcgctattaccatggtgatgcggttttggcagtacatcaatgggcgtggatagcggtttgactcacggggatttccaagtctccaccccattgacgtcaatgggagtttgttttggcaccaaaatcaacgggactttccaaaatgtcgtaac<br>aactccgccccattgacgcaaatgggcggtaggcgtgtacggtgggaggtctatataagcagagctcgtttagtgaaccgtcagatcgcctggagacgccatccacgctgttttgacctccatagaagacaccgggaccgatccagcctccgggggatcgatcccccgatcctgagaacttcagggtgagtttggggacccttgattgttctttctttttcgctat<br>tgtaaaattcatgttatatggagggggcaaagttttcagggtgttgtttagaatgggaagatgtcccttgtatcaccatggaccctcatgataattttgtttctttcactttctactctgttgacaaccattgtctcctcttattttcttttcattttctgtaactttttcgttaaactttagcttgcatttgtaacgaatttttaaattcacttttgtttatttgtcagattgtaagtactttctctaatcac<br>ttttttttcaaggcaatcagggtatattatattgtacttcagcacagttttagagaacaattgttataattaaatgataaggtagaatatttctgcatataaattctggctggcgtggaaatattcttattggtagaaacaactacatcctggtcatcatcctgcctttctctttatggttacaatgatatacactgtttgagatgaggataaaatactctgagtccaaaccgggccc<br>ctctgctaaccatgttcatgccttcttctttttcctacagctcctgggcaacgtgctggttattgtgctgtctcatcattttggcaaagaattgtaatacgactcactatagggcgaattcggatcccccggtagttaattaaaaattagggtgcaagatcatcgataacaagtttgtacaaaaaagcaggcttcgccaccatgggctctaggatcgtgatcaaccgcgagc<br>acctgatgatcgacaggccatacgtgctgctggccgtgctgttcgtgatgtttctgagcctgatcggcctgctggcaatcgcaggaatcaggctgcacagagccgccatctataccgccgagatccacaagagcctgtccaccaacctggacgtgacaaattccatcgagcaccaggtgaaggatgtgctgacaccactgttcaagatcatcggcgacgaa<br>gtgggcctgaggaccccacagcggttcaccgacctggtgaagttcatcagcgataagatcaagtttctgaaccctgatcgggagtacgacttcagagatctgacctggtgcatcaatcccccctgagcggatcaagctggactacgatcagtattgtgccgacgtggccgccgaggagctgatgaacgccctggtgaattccaccctgctggaggccagagc<br>cacaaaccagtttctggccgtgagcaagggcaattgctccggcccaaccacaatccggggccagttctctaacatgtctctgagcctgctggatctgtacctgagcagaggctataatgtgagctccatcgtgaccatgacaagccagggcatgtacggcggcacctatctggtgggcaagcccaacctgtctagcaagggctccgagctgtctcagctgag<br>catgcacagggtgtttgaagtgggcgtgatcaggaacccaggcctgggagcccccgtgtttcacatgaccaattacttcgagcagcccgtgagcaacgatttttctaattgcatggtggccctgggagagctgaagttcgccgccctgtgccacagggaggacagcatcaccatcccatatcagggctctggcaagggcgtgagcttccagctggtgaagct<br>gggcgtgtggaagtcccccaccgacatgagatcctgggtgcctctgtctacagacgatccagtgatcgatcggctgtacctgtcctctcacagaggcgtgatcgccgacaatcaggcaaagtgggcagtgccaaccacaaggaccgacgataagctgagaatggagacatgctttcagcaggcctgtaagggcaagaaccaggccctgtgcgagaatc<br>cagagtgggcacctctgaaggataacagaatccctagctacggcgtgctgagcgtgaatctgtccctgaccgtggagctgaagatcaagatcgccagcggcttcggcccactgatcacacacggctccggcatggacctgtacaagaccaaccacaacaacgtgtactggctgacaatcccacccatgaagaacctggccctgggcgtgatcaatacc<br>ctggagtggatccccaggtttaaggtgtcccctgccctgttcaccgtgccaatcaaggaggccggcgaggattgtcacgccccaacatacctgcccgccgaggtggacggcgatgtgaagctgagctccaacctggtcatcctgcctggccaggacctgcagtacgtgctggccacctatgatacaagccgcgtggagcacgccgtggtgtactacgtgta<br>ctccccctctcggagcttctcctacttttatcctttcagactgccaatcaagggcgtgcctatcgagctgcaggtggagtgctttacctgggataagaagctgtggtgcaggcacttctgcgtgctggcagactctgagagcggaggacacatcacccactccggaatggtgggcatgggcgtgtcttgtaccgtgacacgcgaggacggcacagggggaggc<br>ggctctggaggtgggggtccggtggcggcggaagcgcggccgcggaggtgcagctgcaagcttctggcggaggattcgtgcagcccggcggcagcctgagactgtcctgcgccgccagcggctacacccacagctcttattgtatgggatggttccgccaggcccctggcaaggaacgggaattcgtgtctagcatcgacagcgacggcaccacca<br>gctactacgccgatagcgtgaaaggcagattcaccatcagcagagataatagcaagaacaccgtctacctgcagatgaacagcctgcgggccgaggacaccgccacatactactgcgccgctagattcggccctatgggctgcgtggacctgagcacactgtcttttggccactactggggccagggcacacaggtgaccgtgtcctccggcggcggct<br>ccggcggcggcagctatccctatgatgtgcccgactatgcgtgaactagtgtgaaatagacatcagaattaagaaaaacgtagggtccaagtggttccccgttatggactcgctatctgtcaaccagatcttataccctgaagttcacctagatagcccgatagttaccaataagatagtagccatcctggagtatgctcgactgtttaaacctgcaggcatgcaa<br>gctgatctttttccctctgccaaaaattatggggacatcatgaagccccttgagcatctgacttctggctaataaaggaaatttattttcattgcaatagtgtgttggaattttttgtgtctctcactcggaaggacatatgggagggcaaatcatttaaaacatcagaatgagtatttggtttagagtttggcaacatatgcccatatgctggctgccatgaacaaaggttgg<br>ctataaagaggtcatcagtatatgaaacagccccctgctgtccattccttattccatagaaaagccttgacttgaggttagattttttttatattttgttttgtgttatttttttctttaacatccctaaaattttccttacatgttttactagccagatttttcctcctctcctgactactcccagtcatagctgtccctcttctcttatggagatccctcgaggagctttttgcaaaagcctagg<br>cctccaaaaaagcctcttcactacttctggaatagctcagaggccgaggcggcctcggcctctgcataaataaaaaaaattagtcagccatggggcggagaatgggcggaactgggcggagttaggggcgggatgggcggagttaggggcgggactatggttgctgactaattgctgcattaatgaatcggccaacgcgcggggagaggcggtttgcgt<br>attgggcgctcttccgcttcctcgctcactgactcgctgcgctcggtcgttcggctgcggcgagcggtatcagctcactcaaaggcggtaatacggttatccacagaatcaggggataacgcaggaaagaacatgtgagcaaaaggccagcaaaaggccaggaaccgtaaaaaggccgcgttgctggcgtttttccataggctccgcccccctgacgagc<br>atcacaaaaatcgacgctcaagtcagaggtggcgaaacccgacaggactataaagataccaggcgtttccccctggaagctccctcgtgcgctctcctgttccgaccctgccgcttaccggatacctgtccgcctttctcccttcgggaagcgtggcgctttctcatagctcacgctgtaggtatctcagttcggtgtaggtcgttcgctccaagctgggctgtgtgc<br>acgaacccccgttcagcccgaccgctgcgccttatccggtaactatcgtcttgagtccaacccggtaagacacgacttatcgccactggcagcagccactggtaacaggattagcagagcgaggtatgtaggcggtgctacagagttcttgaagtggtggcctaactacggctacactagaagaacagtatttggtatctgcgctctgctgaagccagttacc<br>ttcggaaaaagagttggtagctcttgatccggcaaacaaaccaccgctggtagcggtggtttttttgtttgcaagcagcagattacgcgcagaaaaaaaggatctcaagaagatcctttgatcttttctacggggtctgacgctcagtggaacgaaaactcacgttaagggattttggtcatgagattatcaaaaaggatcttcacctagatccttttaaattaaaaat<br>gaagttttaaatcaatctaaagtatatatgagtaaacttggtctgacagttaccaatgcttaatcagtgaggcacctatctcagcgatctgtctatttcgttcatccatagttgcctgactccccgtcgtgtagataactacgatacgggagggcttaccatctggccccagtgctgcaatgataccgcgagacccacgctcaccggctccagatttatcagcaataaac<br>cagccagccggaagggccgagcgcagaagtggtcctgcaactttatccgcctccatccagtctattaattgttgccgggaagctagagtaagtagttcgccagttaatagtttgcgcaacgttgttgccattgctacaggcatcgtggtgtcacgctcgtcgtttggtatggcttcattcagctccggttcccaacgatcaaggcgagttacatgatcccccatgttgtg<br>caaaaaagcggttagctccttcggtcctccgatcgttgtcagaagtaagttggccgcagtgttatcactcatggttatggcagcactgcataattctcttactgtcatgccatccgtaagatgcttttctgtgactggtgagtactcaaccaagtcattctgagaatagtgtatgcggcgaccgagttgctcttgcccggcgtcaatacgggataataccgcgccacata<br>gcagaactttaaaagtgctcatcattggaaaacgttcttcggggcgaaaactctcaaggatcttaccgctgttgagatccagttcgatgtaacccactcgtgcacccaactgatcttcagcatcttttactttcaccagcgtttctgggtgagcaaaaacaggaaggcaaaatgccgcaaaaaagggaataagggcgacacggaaatgttgaatactcatactct<br>tcctttttcaatattattgaagcatttatcagggttattgtctcatgagcggatacatatttgaatgtatttagaaaaataaacaaataggggttccgcgcacatttccccgaaaagtgccacctgacgtctaagaaaccattattatcatgacattaacctataaaaataggcgtatcacgaggccctttcgtctcgcgcgtttcggtgatgacggtgaaaacctctgaca<br>catgcagctcccggagacggtcacagcttgtctgtaagcggatgccgggagcagacaagcccgtcagggcgcgtcagcgggtgttggcgggtgtcggggctggcttaactatgcggcatcagagcagattgtactgagagtgcaccattcgacgctctcccttatgcgactcctgcattaggaagcagcccagtagtaggttgaggccgttgagcaccgcc<br>gccgcaaggaatggtctggcttatcgaaattaatacgactcactatagggagaccg |
| pCG-MeV/DMV-H-mMHCIIVHH-HA | Expression of chimerized MeV/DMV H protein with the mouse MHCII targeting VHH clone B and HA tag at the C-terminus | gaattcgagctcgccccgttacataacttacggtaaatggcccgcctggctgaccgcccaacgacccccgcccattgacgtcaataatgacgtatgttcccatagtaacgccaatagggactttccattgacgtcaatgggtggagtatttacggtaaactgcccacttggcagtacatcaagtgtatcatatgccaagtacgccccctattgacgtcaatgacgg<br>taaatggcccgcctggcattatgcccagtacatgaccttatgggactttcctacttggcagtacatctacgtattagtcatcgctattaccatggtgatgcggttttggcagtacatcaatgggcgtggatagcggtttgactcacggggatttccaagtctccaccccattgacgtcaatgggagtttgttttggcaccaaaatcaacgggactttccaaaatgtcgtaac<br>aactccgccccattgacgcaaatgggcggtaggcgtgtacggtgggaggtctatataagcagagctcgtttagtgaaccgtcagatcgcctggagacgccatccacgctgttttgacctccatagaagacaccgggaccgatccagcctccgggggatcgatcccccgatcctgagaacttcagggtgagtttggggacccttgattgttctttctttttcgctat<br>tgtaaaattcatgttatatggagggggcaaagttttcagggtgttgtttagaatgggaagatgtcccttgtatcaccatggaccctcatgataattttgtttctttcactttctactctgttgacaaccattgtctcctcttattttcttttcattttctgtaactttttcgttaaactttagcttgcatttgtaacgaatttttaaattcacttttgtttatttgtcagattgtaagtactttctctaatcac<br>ttttttttcaaggcaatcagggtatattatattgtacttcagcacagttttagagaacaattgttataattaaatgataaggtagaatatttctgcatataaattctggctggcgtggaaatattcttattggtagaaacaactacatcctggtcatcatcctgcctttctctttatggttacaatgatatacactgtttgagatgaggataaaatactctgagtccaaaccgggccc<br>ctctgctaaccatgttcatgccttcttctttttcctacagctcctgggcaacgtgctggttattgtgctgtctcatcattttggcaaagaattgtaatacgactcactatagggcgaattcggatcccccggtagttaattaaaaattagggtgcaagatcatcgataacaagtttgtacaaaaaagcaggcttcgccaccatgggctctaggatcgtgatcaaccgcgagc<br>acctgatgatcgacaggccatacgtgctgctggccgtgctgttcgtgatgtttctgagcctgatcggcctgctggcaatcgcaggaatcaggctgcacagagccgccatctataccgccgagatccacaagagcctgtccaccaacctggacgtgacaaattccatcgagcaccaggtgaaggatgtgctgacaccactgttcaagatcatcggcgacgaa<br>gtgggcctgaggaccccacagcggttcaccgacctggtgaagttcatcagcgataagatcaagtttctgaaccctgatcgggagtacgacttcagagatctgacctggtgcatcaatcccccctgagcggatcaagctggactacgatcagtattgtgccgacgtggccgccgaggaactgatcgtgaccaagttcaaagagctgatgaaccacagcctgga<br>catgtctaagggcagaatcttcccccccaagaactgcagcggaagcgtgatcactcggggccagaccatcaaacctggcctgaccctggtgaacatttacaccacaagaaacttcgaggtgtcattcatggtcacagtgatcagtggcggcatgtacggcaagacctacttcctgaaacctccggaacccgatgatcctttcgagttccaggcttttagaatct<br>ttgaggtgggactggtgcgggatgtgggcagcagggaacctgtgctgcagatgaccaacttcatggtgatcgacgaggacgagggcctgaatttttgtctgctgagcgtaggcgagctgagactggccgccgtgtgcgtgcggggtagacctgttgtgacaaaagatatcggcggctacaaggatgagcctttcaaggtggtcacactgggcatcatcggc<br>ggcggactttctaatcaaaaaaccgaaatctaccccaccattgacagcagcatcgaaaagctgtacatcaccagccaccggggcatcatccggaactccaaggccagatggtccgtgcccgctatcagatctgacgacaaggacaagatggaaaagtgcacccaggccctgtgtaaaagcagacctccaccttcctgcaacagctctgactgggagc<br>ctctgacaagcaatagaatccctgcctacgcctatatcgccctggaaatcaaggaagatagcggccttgagctggatatcacctcgaactacggccctctgatcatccacggcgccggcatggacatctatgaaggcccctccagcaaccaggattggctggctattccccccctgtctcagagcgttctgggcgtgatcaacaaagtggatttcaccgccgg<br>attcgacatcaagccccacacactgaccacggccgttgattacgagagcggcaaatgctacgtgcctgtggaactgagcggagccaaggaccaggacctgaagctggaaagcaatctggtggtgttgcctacaaaggattttggctatgtgacagcaacctacgatacatctagaagcgagcacgccatcgtgtactacgtgtacgacaccgctaggtcc<br>agcagctacttcttccctttccggatcaaggccagaggagagccaatctacctgcgcattgagtgcttcccatggagcagacagctgtggtgccaccactactgcatgatcaacagcaccgtgtccaacgaaatcgtggtggtggataacctggtcagcatcaacatgagctgcagcagaggggggaggcggctctggaggtgggggtccggtggcggcg<br>gaagcgcggccgcgcaggttcaactggtgcagagcggaggcggcctggtccagcccggcggaagcctgggcctgagctgtgccgcttctggcaacatcggcagcagagataatatgggctggtatagacaggcccctggaaaacagagagagtgggtggctacaatcagcggctacggcatcgccacataccgggacagcgtgaagggcagatt<br>caccgtggccaaggacaccgccaagaacatcgtgtccctgcagatgaactacctgaccaccgaagatacagccgtgtactactgctacgcctacgccgtggacagccggaacatcttctggtcccagggcacccaggtgaccgtgtctggcggcggctccggcggcggcagctatccctatgatgtgcccgactatgcgtgaactagtgtgaaatagac<br>atcagaattaagaaaaacgtagggtccaagtggttccccgttatggactcgctatctgtcaaccagatcttataccctgaagttcacctagatagcccgatagttaccaataagatagtagccatcctggagtatgctcgactgtttaaacctgcaggcatgcaagctgatctttttccctctgccaaaaattatggggacatcatgaagccccttgagcatctgactt<br>ctggctaataaaggaaatttattttcattgcaatagtgtgttggaattttttgtgtctctcactcggaaggacatatgggagggcaaatcatttaaaacatcagaatgagtatttggtttagagtttggcaacatatgcccatatgctggctgccatgaacaaaggttggctataaagaggtcatcagtatatgaaacagccccctgctgtccattccttattccatagaaaa<br>gccttgacttgaggttagattttttttatattttgttttgtgttatttttttctttaacatccctaaaattttccttacatgttttactagccagatttttcctcctctcctgactactcccagtcatagctgtccctcttctcttatggagatccctcgaggagctttttgcaaaagcctaggcctccaaaaaagcctcttcactacttctggaatagctcagaggccgaggcggcctcggcct<br>ctgcataaataaaaaaaattagtcagccatggggcggagaatgggcggaactgggcggagttaggggcgggatgggcggagttaggggcgggactatggttgctgactaattgctgcattaatgaatcggccaacgcgcggggagaggcggtttgcgtattgggcgctcttccgcttcctcgctcactgactcgctgcgctcggtcgttcggctgcggcgag<br>cggtatcagctcactcaaaggcggtaatacggttatccacagaatcaggggataacgcaggaaagaacatgtgagcaaaaggccagcaaaaggccaggaaccgtaaaaaggccgcgttgctggcgtttttccataggctccgcccccctgacgagcatcacaaaaatcgacgctcaagtcagaggtggcgaaacccgacaggactataaagatac<br>caggcgtttccccctggaagctccctcgtgcgctctcctgttccgaccctgccgcttaccggatacctgtccgcctttctcccttcgggaagcgtggcgctttctcatagctcacgctgtaggtatctcagttcggtgtaggtcgttcgctccaagctgggctgtgtgcacgaaccccccgttcagcccgaccgctgcgccttatccggtaactatcgtcttgagtccaacc<br>cggtaagacacgacttatcgccactggcagcagccactggtaacaggattagcagagcgaggtatgtaggcggtgctacagagttcttgaagtggtggcctaactacggctacactagaagaacagtatttggtatctgcgctctgctgaagccagttaccttcggaaaaagagttggtagctcttgatccggcaaacaaaccaccgctggtagcggtggtttt<br>tttgtttgcaagcagcagattacgcgcagaaaaaaaggatctcaagaagatcctttgatcttttctacggggtctgacgctcagtggaacgaaaactcacgttaagggattttggtcatgagattatcaaaaaggatcttcacctagatccttttaaattaaaaatgaagttttaaatcaatctaaagtatatatgagtaaacttggtctgacagttaccaatgcttaatca<br>gtgaggcacctatctcagcgatctgtctatttcgttcatccatagttgcctgactccccgtcgtgtagataactacgatacgggagggcttaccatctggccccagtgctgcaatgataccgcgagacccacgctcaccggctccagatttatcagcaataaaccagccagccggaagggccgagcgcagaagtggtcctgcaactttatccgcctccatccagt<br>ctattaattgttgccgggaagctagagtaagtagttcgccagttaatagtttgcgcaacgttgttgccattgctacaggcatcgtggtgtcacgctcgtcgtttggtatggcttcattcagctccggttcccaacgatcaaggcgagttacatgatcccccatgttgtgcaaaaaagcggttagctccttcggtcctccgatcgttgtcagaagtaagttggccgcagtgtta<br>tcactcatggttatggcagcactgcataattctcttactgtcatgccatccgtaagatgcttttctgtgactggtgagtactcaaccaagtcattctgagaatagtgtatgcggcgaccgagttgctcttgcccggcgtcaatacgggataataccgcgccacatagcagaactttaaaagtgctcatcattggaaaacgttcttcggggcgaaaactctcaaggatctt<br>accgctgttgagatccagttcgatgtaacccactcgtgcacccaactgatcttcagcatcttttactttcaccagcgtttctgggtgagcaaaaacaggaaggcaaaatgccgcaaaaaagggaataagggcgacacggaaatgttgaatactcatactcttcctttttcaatattattgaagcatttatcagggttattgtctcatgagcggatacatatttgaatgtatt<br>tagaaaaataaacaaataggggttccgcgcacatttccccgaaaagtgccacctgacgtctaagaaaccattattatcatgacattaacctataaaaataggcgtatcacgaggccctttcgtctcgcgcgtttcggtgatgacggtgaaaacctctgacacatgcagctcccggagacggtcacagcttgtctgtaagcggatgccgggagcagacaagccc<br>cgtcagggcgcgtcagcgggtgttggcgggtgtcggggctggcttaactatgcggcatcagagcagattgtactgagagtgcaccattcgacgctctcccttatgcgactcctgcattaggaagcagcccagtagtaggttgaggccgttgagcaccgccgccgcaaggaatggtctggcttatcgaaattaatacgactcactatagggagaccg |

| | | |
|---|---|---|
| pCG-MeV/CDV-H-mMHCIIVHH-HA | Expression of chimerized MeV/CDV H protein with the mouse MHCII targeting VHH clone B and HA tag at the C-terminus | gaattcgagctcgccccgttacataacttacggtaaatggcccgcctggctgaccgcccaacgacccccgcccattgacgtcaataatgacgtatgttcccatagtaacgccaatagggactttccattgacgtcaatgggtggagtatttacggtaaactgcccacttggcagtacatcaagtgtatcatatgccaagtacgccccctattgacgtcaatgacgg<br>taaatggcccgcctggcattatgcccagtacatgaccttatgggactttcctacttggcagtacatctacgtattagtcatcgctattaccatggtgatgcggttttggcagtacatcaatgggcgtggatagcggtttgactcacggggatttccaagtctccaccccattgacgtcaatgggagtttgttttggcaccaaaatcaacgggactttccaaaatgtcgtaac<br>aactccgccccattgacgcaaatgggcggtaggcgtgtacggtgggaggtctatataagcagagctcgtttagtgaaccgtcagatcgcctggagacgccatccacgctgttttgacctccatagaagacaccgggaccgatccagcctccggggggatcgatcccccgatcctgagaacttcagggtgagtttggggacccttgattgttctttctttttcgctat<br>tgtaaaattcatgttatatggagggggcaaagttttcagggtgttgtttagaatgggaagatgtcccttgtatcaccatggaccctcatgataattttgtttctttcactttctactctgttgacaaccattgtctcctcttattttcttttcattttctgtaactttttcgttaaactttagcttgcatttgtaacgaatttttaaattcacttttgtttatttgtcagattgtaagtactttctctaatcac<br>ttttttttcaaggcaatcagggtatattatattgtacttcagcacagttttagagaacaattgttataattaaatgataaggtagaatatttctgcatataaattctggctggcgtggaaatattcttattggtagaaacaactacatcctggtcatcatcctgcctttctctttatggttacaatgatatacactgtttgagatgaggataaaatactctgagtccaaaccgggccc<br>ctctgctaaccatgttcatgccttcttctttttcctacagctcctgggcaacgtgctggttattgtgctgtctcatcattttggcaaagaattgtaatacgactcactatagggcgaattcggatcccccggtagttaattaaaaattagggtgcaagatcatcgataacaagtttgtacaaaaaagcaggcttcgccaccatgggctctaggatcgtgatcaaccgcgagc<br>acctgatgatcgacaggccatacgtgctgctggccgtgctgttcgtgatgtttctgagcctgatcggcctgctggcaatcgcaggaatcaggctgcacagagccgccatctataccgccgagatccacaagagcctgtccaccaacctggacgtgacaaattccatcgagcaccaggtgaaggatgtgctgacaccactgttcaagatcatcggcgacgaa<br>gtgggcctgaggaccccacagcggttcaccgacctggtgaagttcatcagcgataagatcaagtttctgaaccctgatcgggagtacgacttcagagatctgacctggtgcatcaatcccccctgagcggatcaagctggactacgatcagtattgtgccgacgtggccgccagaaaggccatcgccagcgccgccaaccccatcctgctgtctgctctgagt<br>ggaggcagaggcgatatatttcctcctcatagatgcagcggagccacaaccagcgtgggaaaagttttcccactgtccgtcagcctgagcatgagcctgatctccagaaccagcgaggtgatcaacatgctgaccgccattagcgacggcgtgtacggcaaaacctacctgctcgttcctgatgacatcgaaagagagttcgacaccagagagatccgg<br>gtgttcgagatcggcttcattaaaagatggctgaacgacatgcctctgcttcaaacaaccaactacatggtcctgcctaagaacagcaaggccaaggtgtgcaccatcgctgtgggcgagctgacactggcttccctgtgcgtggaagaaagcaccgtgctgctgtaccacgacagcagcggcagccaggacggaattctggtcgtgaccctgggcatctt<br>ctgggccacccctatggatcacatcgaggaagtgattcccgtggcccacccttctatgaagaagatccacatcaccaaccacagaggattcatcaaggacagcatcgccacctggatggtgcctgccctggcttctgagaaacaggaggaacagaaaggctgtctggaaagcgcctgtcagagaaagacatatcctatgtgcaaccaggccagctggg<br>agcctttcggcggaagacagctgcccagctacggccgcctgaccctgcctctggacgccagcgtggacctgcaactgaacatctctttcacctacggtcctgtgattctgaatggcgatggcatggactactacgagtcccctctgctgaacagcggctggctgaccatccccccctaaggacggcaccatcagcggcctgatcaacaaggccggcagagg<br>cgaccagttcaccgtgctgccacatgtgctgacattcgcccctagagaatcctctggcaactgctacctgcctatccagacatctcagatccgggatagagacgtgctgatcgaatctaatatcgtggtgctgccaacacagagcatcagatacgtgatcgccacttacgacatcagccggagcgatcacgccatcgtgtactacgtgtacgaccctatcagga<br>ccatcagctatacacacccccttcagactgaccacgaagggccggcccgatttcctgcggatcgagtgcttcgtgtgggacgacaacctgtggtgtcaccagttctaccggttcgaggccgacatcgccaattctaccacatctgtggaaaacctggtgcggatccggttctcttgtaaccgggggggaggcggctctggaggtgggggggtccggtggcggcg<br>gaagcgcggccgcgcaggttcaactggtgcagagcggaggcggcctggtccagcccggcggaagcctgggcctgagctgtgccgcttctggcaacatcggcagcagagataatatgggctggtatagacaggcccctggaaaacagagagagtgggtggctacaatcagcggctacggcatcgccacataccgggacagcgtgaagggcagatt<br>caccgtggccaaggacaccgccaagaacatcgtgtccctgcagatgaactacctgaccaccgaagatacagccgtgtactactgctacgcctacgccgtggacagccggaacatcttctggtcccagggcacccaggtgaccgtgtctggcggcggctccggcggcggcagctatccctatgatgtgcccgactatgcgtgaactagtgtgaaatagac<br>atcagaattaagaaaaacgtagggtccaagtggttccccgttatggactcgctatctgtcaaccagatcttataccctgaagttcacctagatagcccgatagttaccaataagatagtagccatcctggagtatgctcgactgtttaaacctgcaggcatgcaagctgatctttttccctctgccaaaaattatggggacatcatgaagccccttgagcatctgactt<br>ctggctaataaaggaaatttattttcattgcaatagtgtgttggaattttttgtgtctctcactcggaaggacatatgggagggcaaatcatttaaaacatcagaatgagtatttggtttagagtttggcaacatatgcccatatgctggctgccatgaacaaaggttggctataaagaggtcatcagtatatgaaacagccccctgctgtccattccttattccatagaaaa<br>gccttgacttgaggttagatttttttatatttgttttgtgttatttttttctttaacatccctaaaattttccttacatgttttactagccagatttttcctcctctcctgactactcccagtcatagctgtccctcttctcttatggagatccctcgaggagctttttgcaaaagcctaggcctccaaaaaagcctcttcactacttctggaatagctcagaggccgaggcggcctcggcct<br>ctgcataaataaaaaaaattagtcagccatggggcggagaatgggcggaactgggcggagttaggggcgggatgggcggagttaggggcgggactatggttgctgactaattgctgcattaatgaatcggccaacgcgcggggagaggcggtttgcgtattgggcgctcttccgcttcctcgctcactgactcgctgcgctcggtcgttcggctgcggcgag<br>cggtatcagctcactcaaaggcggtaatacggttatccacagaatcaggggataacgcaggaaagaacatgtgagcaaaaggccagcaaaaggccaggaaccgtaaaaaggccgcgttgctggcgtttttccataggctccgcccccctgacgagcatcacaaaaatcgacgctcaagtcagaggtggcgaaacccgacaggactataaagatac<br>caggcgtttccccctggaagctccctcgtgcgctctcctgttccgaccctgccgcttaccggatacctgtccgcctttctcccttcgggaagcgtggcgctttctcatagctcacgctgtaggtatctcagttcggtgtaggtcgttcgctccaagctgggctgtgtgcacgaaccccccgttcagcccgaccgctgcgccttatccggtaactatcgtcttgagtccaacc<br>cggtaagacacgacttatcgccactggcagcagccactggtaacaggattagcagagcgaggtatgtaggcggtgctacagagttcttgaagtggtggcctaactacggctacactagaagaacagtatttggtatctgcgctctgctgaagccagttaccttcggaaaaagagttggtagctcttgatccggcaaacaaaccaccgctggtagcggtggttttt<br>ttgtttgcaagcagcagattacgcgcagaaaaaaaggatctcaagaagatcctttgatcttttctacggggtctgacgctcagtggaacgaaaactcacgttaagggattttggtcatgagattatcaaaaaggatcttcacctagatccttttaaattaaaaatgaagttttaaatcaatctaaagtatatatgagtaaacttggtctgacagttaccaatgcttaatca<br>gtgaggcacctatctcagcgatctgtctatttcgttcatccatagttgcctgactccccgtcgtgtagataactacgatacgggagggcttaccatctggccccagtgctgcaatgataccgcgagacccacgctcaccggctccagatttatcagcaataaaccagccagccggaagggccgagcgcagaagtggtcctgcaactttatccgcctccatccagt<br>ctattaattgttgccgggaagctagagtaagtagttcgccagttaatagtttgcgcaacgttgttgccattgctacaggcatcgtggtgtcacgctcgtcgtttggtatggcttcattcagctccggttcccaacgatcaaggcgagttacatgatcccccatgttgtgcaaaaaagcggttagctccttcggtcctccgatcgttgtcagaagtaagttggccgcagtgtta<br>tcactcatggttatggcagcactgcataattctcttactgtcatgccatccgtaagatgcttttctgtgactggtgagtactcaaccaagtcattctgagaatagtgtatgcggcgaccgagttgctcttgcccggcgtcaatacgggataataccgcgccacatagcagaactttaaaagtgctcatcattggaaaacgttcttcggggcgaaaactctcaaggatctt<br>accgctgttgagatccagttcgatgtaacccactcgtgcacccaactgatcttcagcatcttttactttcaccagcgtttctgggtgagcaaaaacaggaaggcaaaatgccgcaaaaaagggaataagggcgacacggaaatgttgaatactcatactcttcctttttcaatattattgaagcatttatcagggttattgtctcatgagcggatacatatttgaatgtatt<br>tagaaaaataaacaaataggggttccgcgcacatttccccgaaaagtgccacctgacgtctaagaaaccattattatcatgacattaacctataaaaataggcgtatcacgaggccctttcgtctcgcgcgtttcggtgatgacggtgaaaacctctgacacatgcagctcccggagacggtcacagcttgtctgtaagcggatgccgggagcagacaagcc<br>cgtcagggcgcgtcagcgggtgttggcgggtgtcggggctggcttaactatgcggcatcagagcagattgtactgagagtgcaccattcgacgctctcccttatgcgactcctgcattaggaagcagcccagtagtaggttgaggccgttgagcaccgccgccgcaaggaatggtctggcttatcgaaattaatacgactcactatagggagaccg |

| | | |
|---|---|---|
| pCG-MeV/DMV-F-465 | Expression of chimerized MeV/DMV F protein at DMV-F AA 465 and MeV-F AA 463 | gaattcgagctcgccccgttacataacttacggtaaatggcccgcctggctgaccgcccaacgacccccgcccattgacgtcaataatgacgtatgttcccatagtaacgccaatagggactttccattgacgtcaatgggtggagtatttacggtaaactgcccacttggcagtacatcaagtgtatcatatgccaagtacgccccctattgacgtcaatgacggtaaatggcccgcctggcattatgcccagtacatgaccttatgggactttcctacttggcagtacatctacgtattagtcatcgctattaccatggtgatgcggttttggcagtacatcaatgggcgtggatagcggtttgactcacggggatttccaagtctccaccccattgacgtcaatgggagtttgttttggcaccaaaatcaacgggactttccaaaatgtcgtaacaactccgccccattgacgcaaatgggcggtaggcgtgtacggtgggaggtctatataagcagagctcgtttagtgaaccgtcagatcgcctggagacgccatccacgctgttttgacctccatagaagacaccgggaccgatccagcctccgggggatcgatcccccgatcctgagaacttcagggtgagtttggggacccttgattgttctttctttttcgctattgtaaaattcatgttatatggagggggcaaagttttcagggtgttgtttagaatgggaagatgtcccttgtatcaccatggaccctcatgataattttgtttctttcactttctactctgttgacaaccattgtctcctcttattttcttttcattttctgtaactttttcgttaaactttagcttgcatttgtaacgaatttttaaattcacttttgtttatttgtcagattgtaagtactttctctaatcacttttttttcaaggcaatcagggtatattatattgtacttcagcacagttttagagaacaattgttataattaaatgataaggtagaatatttctgcatataaattctggctggcgtggaaatattcttattggtagaaacaactacatcctggtcatcatcctgcctttctctttatggttacaatgatatacactgtttgagatgaggataaaatactctgagtccaaaccgggcccctctgctaaccatgttcatgccttcttctttttcctacagctcctgggcaacgtgctggttattgtgctgtctcatcattttggcaaagaattgtaatacgactcactatagggcgaattcggatcccccggtagttaattaaaaattagggtgcaagatcatcgataacaagtttgtacaaaaaagcaggcttcgccaccatggccgccagcaacggcggagtgatgtaccagtcattcctgaccatcatcatcctggtgatcatgaccgagggccaaatccattggggcaacctgagcaagatcggcatcgtgggaacaggcagcgctagctataaggtgatgaccagacctaatcaccaatacctggtgatcaaactgatgcctaacgtgaccatgattgacaactgtacacggaccgaggtgaccgagtacagaaagctgctgaaaaccgtgcttgagcctgtgaaaaatgccctgaccgtgatcaccaagaacatcaagcctatccagagcctgtatccgtatgatgtgccggattatgcgaccacaagcagacggagcaagaggttcgccggagtggtgctggccggcgtcgccctcggtgtcgcaaccgccgcccagatcacagccggcgtggctctgcaccagagcatcatgaacagccagagcattgacaacctgagaaccagcctggagaaaagcaaccaggctatcgaggaaatcagacaggccagccaggagacagtgctggctgtgcagggcgtgcaggacttcatcaacaacgagctgatcccaagcatgcaccagctgtcttgtgaaatgctgggccagaagctgggcctgaagctgcttagatactacaccgagatcctgagtatcttcggcccttctctgcgcgaccccgtgtccgccgagataagcatccaggccctcagctacgccctgggaggcgacatcaacaagatcctggaaaagctgggatatagcggcgccgatctgctggccatcctggaatctcggggcatcaaggccaaggtgacccacgtggatctggaaggctacttcatcgtgctgtccatcgcctacccgaccctgtccgaggttaagggcgttattgtgcacaagctggaggccgtgtcctacaaccttggctctcaagagtggtacaccaccctgcctaagtacgtggccaccaatggctatctgatcagcaacttcgacgagagcagctgtgcttttatgtctgaggtgaccatctgcagccagaacgctctgtacccaatgagccctctgctgcagcagtgcctgcggggctccacagcctcttgcgccagatctctggtttccggcacaattggaaatagattcatcctcagcaaaggcaacctgatcgccaactgcgccagcgtgctgtgcaagtgctacagcaccggaacaatcatcagccaggaccccgacaagctgctgaccttcgtggccgctgataagtgtcctctggtcgaagtggacggcatcaccatccaagtgggcagccgagaataccctgacagcgtatacgtgtccagaatcgacctgggtcctgccatctctctggagcgcctggatgtgggcacaaacctgggcaatgccatcgccaagctggaggacgccaaggagctgctggagagctccgatcagatcctgaggagcatgaagggcctgtctagcacctccatcgtgtacatcctgatcgccgtgtgcctgggaggcctgatcggaatccccgccctgatctgctgttgcaggggccgctgagacccagctttcttgtacaaagtggtactagtgtgaaatagacatcagaattaagaaaaacgtagggtccaagtggttccccgttatggactcgctatctgtcaaccagatcttataccctgaagttcacctagatagcccgatagttaccaataagatagtagccatcctggagtatgctcgactgtttaaacctgcaggcatgcaagctgatctttttccctctgccaaaaattatggggacatcatgaagccccttgagcatctgacttctggctaataaaggaaatttattttcattgcaatagtgtgttggaattttttgtgtctctcactcggaaggacatatgggagggcaaatcatttaaaacatcagaatgagtatttggtttagagtttggcaacatatgcccatatgctggctgccatgaacaaaggttggctataaagaggtcatcagtatatgaaacagccccctgctgtccattccttattccatagaaaagccttgacttgaggttagattttttttatattttgttttgtgttattttttttctttaacatccctaaaattttccttacatgttttactagccagatttttcctcctctcctgactactcccagtcatagctgtccctcttctcttatggagatccctcgaggagctttttgcaaaagcctaggcctccaaaaaagcctcttcactacttctggaatagctcagaggccgaggcggcctcggcctctgcataaataaaaaaaattagtcagccatggggcggagaatgggcggaactgggcggagttaggggcgggatgggcggagttaggggcgggactatggttgctgactaattgctgcattaatgaatcggccaacgcgcggggagaggcggtttgcgtattgggcgctcttccgcttcctcgctcactgactcgctgcgctcggtcgttcggctgcggcgagcggtatcagctcactcaaaggcggtaatacggttatccacagaatcaggggataacgcaggaaagaacatgtgagcaaaaggccagcaaaaggccaggaaccgtaaaaaggccgcgttgctggcgtttttccataggctccgcccccctgacgagcatcacaaaaatcgacgctcaagtcagaggtggcgaaacccgacaggactataaagataccaggcgtttccccctggaagctccctcgtgcgctctcctgttccgaccctgccgcttaccggatacctgtccgcctttctcccttcgggaagcgtggcgctttctcatagctcacgctgtaggtatctcagttcggtgtaggtcgttcgctccaagctgggctgtgtgcacgaaccccccgttcagcccgaccgctgcgccttatccggtaactatcgtcttgagtccaacccggtaagacacgacttatcgccactggcagcagccactggtaacaggattagcagagcgaggtatgtaggcggtgctacagagttcttgaagtggtggcctaactacggctacactagaagaacagtatttggtatctgcgctctgctgaagccagttaccttcggaaaaagagttggtagctcttgatccggcaaacaaaccaccgctggtagcggtggtttttttgtttgcaagcagcagattacgcgcagaaaaaaaggatctcaagaagatcctttgatcttttctacggggtctgacgctcagtggaacgaaaactcacgttaagggattttggtcatgagattatcaaaaaggatcttcacctagatccttttaaattaaaaatgaagttttaaatcaatctaaagtatatatgagtaaacttggtctgacagttaccaatgcttaatcagtgaggcacctatctcagcgatctgtctatttcgttcatccatagttgcctgactccccgtcgtgtagataactacgatacgggagggcttaccatctggccccagtgctgcaatgataccgcgagacccacgctcaccggctccagatttatcagcaataaaccagccagccggaagggccgagcgcagaagtggtcctgcaactttatccgcctccatccagtctattaattgttgccgggaagctagagtaagtagttcgccagttaatagtttgcgcaacgttgttgccattgctacaggcatcgtggtgtcacgctcgtcgtttggtatggcttcattcagctccggttcccaacgatcaaggcgagttacatgatcccccatgttgtgcaaaaaagcggttagctccttcggtcctccgatcgttgtcagaagtaagttggccgcagtgttatcactcatggttatggcagcactgcataattctcttactgtcatgccatccgtaagatgcttttctgtgactggtgagtactcaaccaagtcattctgagaatagtgtatgcggcgaccgagttgctcttgcccggcgtcaatacgggataataccgcgccacatagcagaactttaaaagtgctcatcattggaaaacgttcttcggggcgaaaactctcaaggatcttaccgctgttgagatccagttcgatgtaacccactcgtgcacccaactgatcttcagcatcttttactttcaccagcgtttctgggtgagcaaaaacaggaaggcaaaatgccgcaaaaaagggaataagggcgacacggaaatgttgaatactcatactcttcctttttcaatattattgaagcatttatcagggttattgtctcatgagcggatacatatttgaatgtatttagaaaaataaacaaataggggttccgcgcacatttccccgaaaagtgccacctgacgtctaagaaaccattattatcatgacattaacctataaaaataggcgtatcacgaggccctttcgtctcgcgcgtttcggtgatgacggtgaaaacctctgacacatgcagctcccggagacggtcacagcttgtctgtaagcggatgccgggagcagacaagcccgtcagggcgcgtcagcgggtgttggcgggtgtcggggctggcttaactatgcggcatcagagcagattgtactgagagtgcaccattcgacgctctcccttatgcgactcctgcattaggaagcagcccagtagtaggttgaggccgttgagcaccgccgccgcaaggaatggtctggcttatcgaaattaatacgactcactatagggagaccg |

| | | |
|---|---|---|
| pCG-MeV/DMV-F-407 | Expression of chimerized MeV/DMV F protein at DMV-F AA 407 and MeV-F AA 405 | gaattcgagctcgccccgttacataacttacggtaaatggcccgcctggctgaccgcccaacgacccccgcccattgacgtcaataatgacgtatgttcccatagtaacgccaatagggactttccattgacgtcaatgggtggagtatttacggtaaactgcccacttggcagtacatcaagtgtatcatatgccaagtacgccccctattgacgtcaatgacgg<br>taaatggcccgcctggcattatgcccagtacatgaccttatgggactttcctacttggcagtacatctacgtattagtcatcgctattaccatggtgatgcggttttggcagtacatcaatgggcgtggatagcggtttgactcacggggatttccaagtctccaccccattgacgtcaatgggagtttgttttggcaccaaaatcaacgggactttccaaaatgtcgtaac<br>aactccgccccattgacgcaaatgggcggtaggcgtgtacggtgggaggtctatataagcagagctcgtttagtgaaccgtcagatcgcctggagacgccatccacgctgttttgacctccatagaagacaccgggaccgatccagcctccgggggatcgatcccccgatcctgagaacttcagggtgagtttggggacccttgattgttctttctttttcgctat<br>tgtaaaattcatgttatatggagggggcaaagttttcagggtgttgtttagaatgggaagatgtcccttgtatcaccatggaccctcatgataattttgtttctttcactttctactctgttgacaaccattgtctcctcttattttcttttcattttctgtaactttttcgttaaactttagcttgcatttgtaacgaatttttaaattcacttttgtttattgtcagattgtaagtactttctctaatcac<br>tttttttttcaaggcaatcagggtatattatattgtacttcagcacagttttagagaacaattgttataattaaatgataaggtagaatatttctgcatataaattctggctggcgtggaaatattcttattggtagaaacaactacatcctggtcatcatcctgcctttctctttatggttacaatgatatacactgtttgagatgaggataaaatactctgagtccaaaccgggccc<br>ctctgctaaccatgttcatgccttcttctttttcctacagctcctgggcaacgtgctggttattgtgctgtctcatcattttggcaaagaattgtaatacgactcactatagggcgaattcggatcccccggtagttaattaaaaattagggtgcaagatcatcgataacaagtttgtacaaaaaagcaggcttcgccaccatggccgccagcaacggcggagtgatgtac<br>cagtcattcctgaccatcatcatcctggtgatcatgaccgagggccaaatccattggggcaacctgagcaagatcggcatcgtgggaacaggcagcgctagctataaggtgatgaccagacctaatcaccaatacctggtgatcaaactgatgcctaacgtgaccatgattgacaactgtacacggaccgaggtgaccgagtacagaaagctgctgaaa<br>accgtgcttgagcctgtgaaaaatgccctgaccgtgatcaccaagaacatcaagcctatccagagcctgtatccgtatgatgtgccggattatgcgaccacaagcagacggagcaagaggttcgccggagtggtgctggccggcgtcgccctcggtgtcgcaaccgccgcccagatcacagccggcgtggctctgcaccagagcatcatgaacagcca<br>gagcattgacaacctgagaaccagcctggagaaaagcaaccaggctatcgaggaaatcagacaggccagccaggagacagtgctggctgtgcagggcgtgcaggacttcatcaacaacgagctgatcccaagcatgcaccagctgtcttgtgaaatgctgggccagaagctgggcctgaagctgcttagatactacaccgagatcctgagtatcttc<br>ggcccttctctgcgcgaccccgtgtccgccgagataagcatccaggccctcagctacgccctgggaggcgacatcaacaagatcctggaaaagctgggatatagcggcgccgatctgctggccatcctggaatctcggggcatcaaggccaaggtgacccacgtggatctggaaggctacttcatcgtgctgtccatcgcctacccgaccctgtccgaggt<br>taagggcgttattgtgcacaagctggaggccgtgtcctacaaccttggctctcaagagtggtacaccaccctgcctaagtacgtggccaccaatggctatctgatcagcaacttcgacgagagcagctgtgcttttatgtctgaggtgaccatctgcagccagaacgctctgtacccaatgagccctctgctgcagcagtgcctgcggggctccacagcctcttgc<br>gccagatctctggtttccggcacaattggaaatagattcatcctcagcaaaggcaacctgatcgccaactgcgccagcgtgctgtgcaagtgctataccacaggcaccatcatcaaccaggaccccgataagatcctgacatacatcgcagcagaccactgccctgtggtggaagtgaatggcgtgaccatccaagtgggcagccggagatacccagac<br>gccgtgtatctgcacaggatcgatctgggaccacctatctccctggagcgcctggatgtgggcacaaacctgggcaatgccatcgccaagctggaggacgccaaggagctgctggagagctccgatcagatcctgaggagcatgaagggcctgtctagcacctccatcgtgtacatcctgatcgccgtgtgcctgggaggcctgatcggaatccccgccct<br>gatctgctgttgcaggggccgctgagacccagctttcttgtacaaagtggtactagtgtgaaatagacatcagaattaagaaaaacgtagggtccaagtggttccccgttatggactcgctatctgtcaaccagatcttataccctgaagttcacctagatagcccgatagttaccaataagatagtagccatcctggagtatgctcgactgtttaaacctgcaggc<br>atgcaagctgatctttttccctctgccaaaaattatggggacatcatgaagccccttgagcatctgacttctggctaataaaggaaatttattttcattgcaatagtgtgttggaatttttgtgtctctcactcggaaggacatatgggagggcaaatcatttaaaacatcagaatgagtatttggtttagagtttggcaacatatgcccatatgctggctgccatgaacaaa<br>ggttggctataaagaggtcatcagtatatgaaacagccccctgctgtccattccttattccatagaaaagccttgacttgaggttagatttttttatattttgttttgtgttatttttttctttaacatccctaaaattttccttacatgttttactagccagatttttcctcctctcctgactactcccagtcatagctgtccctcttctcttatggagatccctcgaggagctttttgcaaaag<br>cctaggcctccaaaaaagcctcttcactacttctggaatagctcagaggccgaggcggcctcggcctctgcataaataaaaaaaattagtcagccatggggcggagaatgggcggaactgggcggagttaggggcgggatgggcggagttaggggcgggactatggttgctgactaattgctgcattaatgaatcggccaacgcgcggggagaggcg<br>gtttgcgtattgggcgctcttccgcttcctcgctcactgactcgctgcgctcggtcgttcggctgcggcgagcggtatcagctcactcaaaggcggtaatacggttatccacagaatcaggggataacgcaggaaagaacatgtgagcaaaaggccagcaaaaggccaggaaccgtaaaaaggccgcgttgctggcgtttttccataggctccgcccccctg<br>acgagcatcacaaaaatcgacgctcaagtcagaggtggcgaaacccgacaggactataaagataccaggcgtttccccctggaagctccctcgtgcgctctcctgttccgaccctgccgcttaccggatacctgtccgcctttctcccttcgggaagcgtggcgctttctcatagctcacgctgtaggtatctcagttcggtgtaggtcgttcgctccaagctgggc<br>tgtgtgcacgaaccccccgttcagcccgaccgctgcgccttatccggtaactatcgtcttgagtccaacccggtaagacacgacttatcgccactggcagcagccactggtaacaggattagcagagcgaggtatgtaggcggtgctacagagttcttgaagtggtggcctaactacggctacactagaagaacagtatttggtatctgcgctctgctgaagcc<br>agttaccttcggaaaaagagttggtagctcttgatccggcaaacaaaccaccgctggtagcggtggtttttttgtttgcaagcagcagattacgcgcagaaaaaaaggatctcaagaagatcctttgatcttttctacggggtctgacgctcagtggaacgaaaactcacgttaagggattttggtcatgagattatcaaaaaggatcttcacctagatccttttaaatt<br>aaaaatgaagttttaaatcaatctaaagtatatatgagtaaacttggtctgacagttaccaatgcttaatcagtgaggcacctatctcagcgatctgtctatttcgttcatccatagttgcctgactccccgtcgtgtagataactacgatacgggagggcttaccatctggccccagtgctgcaatgataccgcgagacccacgctcaccggctccagatttatcagca<br>ataaaccagccagccggaagggccgagcgcagaagtggtcctgcaactttatccgcctccatccagtctattaattgttgccgggaagctagagtaagtagttcgccagttaatagtttgcgcaacgttgttgccattgctacaggcatcgtggtgtcacgctcgtcgtttggtatggcttcattcagctccggttcccaacgatcaaggcgagttacatgatcccc<br>atgttgtgcaaaaaagcggttagctccttcggtcctccgatcgttgtcagaagtaagttggccgcagtgttatcactcatggttatggcagcactgcataattctcttactgtcatgccatccgtaagatgcttttctgtgactggtgagtactcaaccaagtcattctgagaatagtgtatgcggcgaccgagttgctcttgcccggcgtcaatacgggataataccgcgc<br>cacatagcagaactttaaaagtgctcatcattggaaaacgttcttcggggcgaaaactctcaaggatcttaccgctgttgagatccagttcgatgtaacccactcgtgcacccaactgatcttcagcatcttttactttcaccagcgtttctgggtgagcaaaaacaggaaggcaaaatgccgcaaaaaagggaataagggcgacacggaaatgttgaatactc<br>atactcttcctttttcaatattattgaagcatttatcagggttattgtctcatgagcggatacatatttgaatgtatttagaaaaataaacaaataggggttccgcgcacatttccccgaaaagtgccacctgacgtctaagaaaccattattatcatgacattaacctataaaaataggcgtatcacgaggccctttcgtctcgcgcgtttcggtgatgacggtgaaaacct<br>ctgacacatgcagctcccggagacggtcacagcttgtctgtaagcggatgccgggagcagacaagcccgtcagggcgcgtcagcgggtgttggcgggtgtcggggctggcttaactatgcggcatcagagcagattgtactgagagtgcaccattcgacgctctcccttatgcgactcctgcattaggaagcagcccagtagtaggttgaggccgttgagc<br>accgccgccgcaaggaatggtctggcttatcgaaattaatacgactcactatagggagaccg |

| | | |
|---|---|---|
| pCG-MeV/DMV-F-311 | Expression of chimerized MeV/DMV F protein at DMV-F AA 311 and MeV-F AA 309 | gaattcgagctcgccccgttacataacttacggtaaatggcccgcctggctgaccgcccaacgacccccgcccattgacgtcaataatgacgtatgttcccatagtaacgccaatagggactttccattgacgtcaatgggtggagtatttacggtaaactgcccacttggcagtacatcaagtgtatcatatgccaagtacgccccctattgacgtcaatgacggtaaatggcccgcctggcattatgcccagtacatgaccttatgggactttcctacttggcagtacatctacgtattagtcatcgctattaccatggtgatgcggttttggcagtacatcaatgggcgtggatagcggtttgactcacggggatttccaagtctccaccccattgacgtcaatgggagtttgttttggcaccaaaatcaacgggactttccaaaatgtcgtaacaactccgccccattgacgcaaatgggcggtaggcgtgtacggtgggaggtctatataagcagagctcgtttagtgaaccgtcagatcgcctggagacgccatccacgctgttttgacctccatagaagacaccgggaccgatccagcctccggggggatcgatcccccgatcctgagaacttcagggtgagtttggggacccttgattgttctttctttttcgctattgtaaaattcatgttatatggagggggcaaagttttcagggtgttgtttagaatgggaagatgtcccttgtatcaccatggaccctcatgataattttgtttctttcactttctactctgttgacaaccattgtctcctcttattttcttttcattttctgtaactttttcgttaaactttagcttgcatttgtaacgaatttttaaattcacttttgtttatttgtcagattgtaagtactttctctaatcactttttttttcaaggcaatcagggtatattatattgtacttcagcacagttttagagaacaattgttataattaaatgataaggtagaatatttctgcatataaattctggctggcgtggaaatattcttattggtagaaacaactacatcctggtcatcatcctgcctttctctttatggttacaatgatatacactgtttgagatgaggataaaatactctgagtccaaaccgggcccctctgctaaccatgttcatgccttcttctttttcctacagctcctgggcaacgtgctggttattgtgctgtctcatcattttggcaaagaattgtaatacgactcactatagggcgaattcggatcccccggtagttaattaaaaattagggtgcaagatcatcgataacaagtttgtacaaaaaagcaggcttcgccaccatggccgccagcaacggcggagtgatgtaccagtcattcctgaccatcatcatcctggtgatcatgaccgagggccaaatccattggggcaacctgagcaagatcggcatcgtgggaacaggcagcgctagctataaggtgatgaccagacctaatcaccaatacctggtgatcaaactgatgcctaacgtgaccatgattgacaactgtacacggaccgaggtgaccgagtacagaaagctgctgaaaaccgtgcttgagcctgtgaaaaatgccctgaccgtgatcaccaagaacatcaagcctatccagagcctgtatccgtatgatgtgccggattatgcgaccacaagcagacggagcaagaggttcgccggagtggtgctggccggcgtcgccctcggtgtcgcaaccgccgcccagatcacagccggcgtggctctgcaccagagcatcatgaacagccagagcattgacaacctgagaaccagcctggagaaaagcaaccaggctatcgaggaaatcagacaggccagccaggagacagtgctggctgtgcagggcgtgcaggacttcatcaacaacgagctgatcccaagcatgcaccagctgtcttgtgaaatgctgggccagaagctgggcctgaagctgcttagatactacaccgagatcctgagtatcttcggcccttctctgcgcgaccccgtgtccgccgagataagcatccaggccctcagctacgccctgggaggcgacatcaacaagatcctggaaaagctgggatatagcggcgccgatctgctggccatcctggaatctggggcatcaaggccaaggtgacccacgtggatctggaaggctacttcatcgtgctgtccatcgcctacccgaccctgtccgaggttaagggcgttattgtgcacaagctggagggcgtgagctataatatcggctcccaggagtggtacaccacagtgcctaagtatgtggccacccagggctacctgatctctaacttcgacgagtcctcttgtacctttatgccagagggcacagtgtgctcccagaatgccctgtacccaatgtctcccctgctgcaggagtgtctgcggggctctaccaagagctgcgccagaacactggtgagcggctccttcggcaatcggttcatcctgtcccagggcaacctgatcgccaattgtgcctctatcctgtgcaagtgctataccacaggcaccatcatcaaccaggaccccgataagatcctgacatacatcgcagcagaccactgccctgtggtggaagtgaatggcgtgaccatccaagtgggcagccggagatacccagacgccgtgtatctgcacaggatcgatctgggaccacctatctccctggagcgcctggatgtgggcacaaacctgggcaatgccatcgccaagctggaggacgccaaggagctgctggagagctccgatcagatcctgaggagcatgaagggcctgtctagcacctccatcgtgtacatcctgatcgccgtgtgcctgggaggcctgatcggaatccccgccctgatctgctgttgcaggggccgctgagacccagctttcttgtacaaagtggtactagtgtgaaatagacatcagaattaagaaaaacgtagggtccaagtggttccccgttatggactcgctatctgtcaaccagatcttataccctgaagttcacctagatagcccgatagttaccaataagatagtagccatcctggagtatgctcgactgtttaaacctgcaggcatgcaagctgatctttttccctctgccaaaaattatggggacatcatgaagccccttgagcatctgacttctggctaataaaggaaatttattttcattgcaatagtgtgttggaattttttgtgtctctcactcggaaggacatatgggagggcaaatcatttaaaacatcagaatgagtatttggtttagagtttggcaacatatgcccatatgctggctgccatgaacaaaggttggctataaagaggtcatcagtatatgaaacagccccctgctgtccattccttattccatagaaaagccttgacttgaggttagattttttttatattttgttttgtgttatttttttctttaacatccctaaaattttccttacatgttttactagccagatttttcctcctctcctgactactcccagtcatagctgtccctcttctcttatggagatccctcgaggagctttttgcaaaagcctaggcctccaaaaaagcctcttcactacttctggaatagctcagaggccgaggcggcctcggcctctgcataaataaaaaaaattagtcagccatggggcggagaatgggcggaactgggcggagttaggggcgggatgggcggagttaggggcgggactatggttgctgactaattgctgcattaatgaatcggccaacgcgcggggagaggcggtttgcgtattgggcgctcttccgcttcctcgctcactgactcgctgcgctcggtcgttcggctgcggcgagcggtatcagctcactcaaaggcggtaatacggttatccacagaatcaggggataacgcaggaaagaacatgtgagcaaaaggccagcaaaaggccaggaaccgtaaaaaggccgcgttgctggcgtttttccataggctccgcccccctgacgagcatcacaaaaatcgacgctcaagtcagaggtggcgaaacccgacaggactataaagataccaggcgtttccccctggaagctccctcgtgcgctctcctgttccgaccctgccgcttaccggatacctgtccgcctttctcccttcgggaagcgtggcgctttctcatagctcacgctgtaggtatctcagttcggtgtaggtcgttcgctccaagctgggctgtgtgcacgaaccccccgttcagcccgaccgctgcgccttatccggtaactatcgtcttgagtccaacccggtaagacacgacttatcgccactggcagcagccactggtaacaggattagcagagcgaggtatgtaggcggtgctacagagttcttgaagtggtggcctaactacggctacactagaagaacagtatttggtatctgcgctctgctgaagccagttaccttcggaaaaagagttggtagctcttgatccggcaaacaaaccaccgctggtagcggtggtttttttgtttgcaagcagcagattacgcgcagaaaaaaaggatctcaagaagatcctttgatcttttctacggggtctgacgctcagtggaacgaaaactcacgttaagggattttggtcatgagattatcaaaaaggatcttcacctagatccttttaaattaaaaatgaagttttaaatcaatctaaagtatatatgagtaaacttggtctgacagttaccaatgcttaatcagtgaggcacctatctcagcgatctgtctatttcgttcatccatagttgcctgactccccgtcgtgtagataactacgatacgggagggcttaccatctggccccagtgctgcaatgataccgcgagacccacgctcaccggctccagatttatcagcaataaaccagccagccggaagggccgagcgcagaagtggtcctgcaactttatccgcctccatccagtctattaattgttgccgggaagctagagtaagtagttcgccagttaatagtttgcgcaacgttgttgccattgctacaggcatcgtggtgtcacgctcgtcgtttggtatggcttcattcagctccggttcccaacgatcaaggcgagttacatgatcccccatgttgtgcaaaaaagcggttagctccttcggtcctccgatcgttgtcagaagtaagttggccgcagtgttatcactcatggttatggcagcactgcataattctcttactgtcatgccatccgtaagatgcttttctgtgactggtgagtactcaaccaagtcattctgagaatagtgtatgcggcgaccgagttgctcttgcccggcgtcaatacgggataataccgcgccacatagcagaactttaaaagtgctcatcattggaaaacgttcttcggggcgaaaactctcaaggatcttaccgctgttgagatccagttcgatgtaacccactcgtgcacccaactgatcttcagcatcttttactttcaccagcgtttctgggtgagcaaaaacaggaaggcaaaatgccgcaaaaaagggaataagggcgacacggaaatgttgaatactcatactcttcctttttcaatattattgaagcatttatcagggttattgtctcatgagcggatacatatttgaatgtatttagaaaaataaacaaataggggttccgcgcacatttccccgaaaagtgccacctgacgtctaagaaaccattattatcatgacattaacctataaaaataggcgtatcacgaggccctttcgtctcgcgcgtttcggtgatgacggtgaaaacctctgacacatgcagctcccggagacggtcacagcttgtctgtaagcggatgccgggagcagacaagcccgtcagggcgcgtcagcgggtgttggcgggtgtcggggctggcttaactatgcggcatcagagcagattgtactgagagtgcaccattcgacgctctcccttatgcgactcctgcattaggaagcagcccagtagtaggttgaggccgttgagcaccgccgccgcaaggaatggtctggcttatcgaaattaatacgactcactatagggagaccg |
| pCG-MeV/DMV-F-229 | Expression of chimerized MeV/DMV F protein at DMV-F AA 229 and MeV-F AA 228 | gaattcgagctcgccccgttacataacttacggtaaatggcccgcctggctgaccgcccaacgacccccgcccattgacgtcaataatgacgtatgttcccatagtaacgccaatagggactttccattgacgtcaatgggtggagtatttacggtaaactgcccacttggcagtacatcaagtgtatcatatgccaagtacgccccctattgacgtcaatgacggtaaatggcccgcctggcattatgcccagtacatgaccttatgggactttcctacttggcagtacatctacgtattagtcatcgctattaccatggtgatgcggttttggcagtacatcaatgggcgtggatagcggtttgactcacggggatttccaagtctccaccccattgacgtcaatgggagtttgttttggcaccaaaatcaacgggactttccaaaatgtcgtaacaactccgccccattgacgcaaatgggcggtaggcgtgtacggtgggaggtctatataagcagagctcgtttagtgaaccgtcagatcgcctggagacgccatccacgctgttttgacctccatagaagacaccgggaccgatccagcctccggggggatcgatcccccgatcctgagaacttcagggtgagtttggggacccttgattgttctttctttttcgctattgtaaaattcatgttatatggagggggcaaagttttcagggtgttgtttagaatgggaagatgtcccttgtatcaccatggaccctcatgataattttgtttctttcactttctactctgttgacaaccattgtctcctcttattttcttttcattttctgtaactttttcgttaaactttagcttgcatttgtaacgaatttttaaattcacttttgtttatttgtcagattgtaagtactttctctaatcactttttttttcaaggcaatcagggtatattatattgtacttcagcacagttttagagaacaattgttataattaaatgataaggtagaatatttctgcatataaattctggctggcgtggaaatattcttattggtagaaacaactacatcctggtcatcatcctgcctttctctttatggttacaatgatatacactgtttgagatgaggataaaatactctgagtccaaaccgggcccctctgctaaccatgttcatgccttcttctttttcctacagctcctgggcaacgtgctggttattgtgctgtctcatcattttggcaaagaattgtaatacgactcactatagggcgaattcggatcccccggtagttaattaaaaattagggtgcaagatcatcgataacaagtttgtacaaaaaagcaggcttcgccaccatggccgccagcaacggcggagtgatgtaccagtcattcctgaccatcatcatcctggtgatcatgaccgagggccaaatccattggggcaacctgagcaagatcggcatcgtgggaacaggcagcgctagctataaggtgatgaccagacctaatcaccaatacctggtgatcaaactgatgcctaacgtgaccatgattgacaactgtacacggaccgaggtgaccgagtacagaaagctgctgaaaaccgtgcttgagcctgtgaaaaatgccctgaccgtgatcaccaagaacatcaagcctatccagagcctgtatccgtatgatgtgccggattatgcgaccacaagcagacggagcaagaggttcgccggagtggtgctggccggcgtcgccctcggtgtcgcaaccgccgcccagatcacagccggcgtggctctgcaccagagcatcatgaacagccagagcattgacaacctgagaaccagcctggagaaaagcaaccaggctatcgaggaaatcagacaggccagccaggagacagtgctggctgtgcagggcgtgcaggacttcatcaacaacgagctgatcccaagcatgcaccagctgtcttgtgaaatgctgggccagaagctgggcctgaagctgcttagatactacaccgagatcctgagtatcttcggcccttctctgcgggaccccatcagcgccgagatctccatccaggccctgtcctatgcactgggaggcgacatcaacaaggtgctggagaagctgggctactctggcggcgatctgctgggcatcctggagagcggggcatcaaggccagaatcacccacgtggatacagagtcttacttcatcgtgctgagcatcgcctaccccacactgtccgagatcaagggcgtgatcgtgcacaggctggagggcgtgagctataatatcggctcccaggagtggtacaccacagtgcctaagtatgtggccacccagggctacctgatctctaacttcgacgagtcctcttgtacctttatgccagagggcacagtgtgctcccagaatgccctgtacccaatgtctcccctgctgcaggagtgtctgcggggctctaccaagagctgcgccagaacactggtgagcggctccttcggcaatcggttcatcctgtcccagggcaacctgatcgccaattgtgcctctatcctgtgcaagtgctataccacaggcaccatcatcaaccaggaccccgataagatcctgacatacatcgcagcagaccactgccctgtggtggaagtgaatggcgtgaccatccaagtgggcagccggagatacccagacgccgtgtatctgcacaggatcgatctgggaccacctatctccctggagcgcctggatgtgggcacaaacctgggcaatgccatcgccaagctggaggacgccaaggagctgctggagagctccgatcagatcctgaggagcatgaagggcctgtctagcacctccatcgtgtacatcctgatcgccgtgtgcctgggaggcctgatcggaatccccgccctgatctgctgttgcaggggccgctgagacccagctttcttgtacaaagtggtactagtgtgaaatagacatcagaattaagaaaaacgtagggtccaagtggttccccgttatggactcgctatctgtcaaccagatcttataccctgaagttcacctagatagcccgatagttaccaataagatagtagccatcctggagtatgctcgactgtttaaacctgcaggcatgcaagctgatctttttccctctgccaaaaattatggggacatcatgaagccccttgagcatctgacttctggctaataaaggaaatttattttcattgcaatagtgtgttggaattttttgtgtctctcactcggaaggacatatgggagggcaaatcatttaaaacatcagaatgagtatttggtttagagtttggcaacatatgcccatatgctggctgccatgaacaaaggttggctataaagaggtcatcagtatatgaaacagccccctgctgtccattccttattccatagaaaagccttgacttgaggttagattttttttatattttgttttgtgttatttttttctttaacatccctaaaattttccttacatgttttactagccagatttttcctcctctcctgactactcccagtcatagctgtccctcttctcttatggagatccctcgaggagctttttgcaaaagcctaggcctccaaaaaagcctcttcactacttctggaatagctcagaggccgaggcggcctcggcctctgcataaataaaaaaaattagtcagccatggggcggagaatgggcggaactgggcggagttaggggcgggatgggcggagttaggggcgggactatggttgctgactaattgctgcattaatgaatcggccaacgcgcggggagaggcggtttgcgtattgggcgctcttccgcttcctcgctcactgactcgctgcgctcggtcgttcggctgcggcgagcggtatcagctcactcaaaggcggtaatacggttatccacagaatcaggggataacgcaggaaagaacatgtgagcaaaaggccagcaaaaggccaggaaccgtaaaaaggccgcgttgctggcgtttttccataggctccgcccccctgacgagcatcacaaaaatcgacgctcaagtcagaggtggcgaaacccgacaggactataaagataccaggcgtttccccctggaagctccctcgtgcgctctcctgttccgaccctgccgcttaccggatacctgtccgcctttctcccttcgggaagcgtggcgctttctcatagctcacgctgtaggtatctcagttcggtgtaggtcgttcgctccaagctgggctgtgtgcacgaaccccccgttcagcccgaccgctgcgccttatccggtaactatcgtcttgagtccaacccggtaagacacgacttatcgccactggcagcagccactggtaacaggattagcagagcgaggtatgtaggcggtgctacagagttcttgaagtggtggcctaactacggctacactagaagaacagtatttggtatctgcgctctgctgaagccagttaccttcggaaaaagagttggtagctcttgatccggcaaacaaaccaccgctggtagcggtggtttttttgtttgcaagcagcagattacgcgcagaaaaaaaggatctcaagaagatcctttgatcttttctacggggtctgacgctcagtggaacgaaaactcacgttaagggattttggtcatgagattatcaaaaaggatcttcacctagatccttttaaattaaaaatgaagttttaaatcaatctaaagtatatatgagtaaacttggtctgacagttaccaatgcttaatcagtgaggcacctatctcagcgatctgtctatttcgttcatccatagttgcctgactccccgtcgtgtagataactacgatacgggagggcttaccatctggccccagtgctgcaatgataccgcgagacccacgctcaccggctccagatttatcagcaataaaccagccagccggaagggccgagcgcagaagtggtcctgcaactttatccgcctccatccagtctattaattgttgccgggaagctagagtaagtagttcgccagttaatagtttgcgcaacgttgttgccattgctacaggcatcgtggtgtcacgctcgtcgtttggtatggcttcattcagctccggttcccaacgatcaaggcgagttacatgatcccccatgttgtgcaaaaaagcggttagctccttcggtcctccgatcgttgtcagaagtaagttggccgcagtgttatcactcatggttatggcagcactgcataattctcttactgtcatgccatccgtaagatgcttttctgtgactggtgagtactcaaccaagtcattctgagaatagtgtatgcggcgaccgagttgctcttgcccggcgtcaatacgggataataccgcgccacatagcagaactttaaaagtgctcatcattggaaaacgttcttcggggcgaaaactctcaaggatcttaccgctgttgagatccagttcgatgtaacccactcgtgcacccaactgatcttcagcatcttttactttcaccagcgtttctgggtgagcaaaaacaggaaggcaaaatgccgcaaaaaagggaataagggcgacacggaaatgttgaatactcatactcttcctttttcaatattattgaagcatttatcagggttattgtctcatgagcggatacatatttgaatgtatttagaaaaataaacaaataggggttccgcgcacatttccccgaaaagtgccacctgacgtctaagaaaccattattatcatgacattaacctataaaaataggcgtatcacgaggccctttcgtctcgcgcgtttcggtgatgacggtgaaaacctctgacacatgcagctcccggagacggtcacagcttgtctgtaagcggatgccgggagcagacaagcccgtcagggcgcgtcagcgggtgttggcgggtgtcggggctggcttaactatgcggcatcagagcagattgtactgagagtgcaccattcgacgctctcccttatgcgactcctgcattaggaagcagcccagtagtaggttgaggccgttgagcaccgccgccgcaaggaatggtctggcttatcgaaattaatacgactcactatagggagaccg |

| | | |
|---|---|---|
| pCG-MeV/DMV-F-123 | Expression of chimerized MeV/DMV F protein at DMV-F AA 123 and MeV-F AA 122 | gaattcgagctcgccccgttacataacttacggtaaatggcccgcctggctgaccgcccaacgacccccgcccattgacgtcaataatgacgtatgttcccatagtaacgccaatagggactttccattgacgtcaatgggtggagtatttacggtaaactgcccacttggcagtacatcaagtgtatcatatgccaagtacgccccctattgacgtcaatgacgg<br>taaatggcccgcctggcattatgcccagtacatgaccttatgggactttcctacttggcagtacatctacgtattagtcatcgctattaccatggtgatgcggttttggcagtacatcaatgggcgtggatagcggtttgactcacggggatttccaagtctccaccccattgacgtcaatgggagtttgttttggcaccaaaatcaacgggactttccaaaatgtcgtaac<br>aactccgccccattgacgcaaatgggcggtaggcgtgtacggtgggaggtctatataagcagagctcgtttagtgaaccgtcagatcgcctggagacgccatccacgctgttttgacctccatagaagacaccgggaccgatccagcctccgggggatcgatcccccgatcctgagaacttcagggtgagtttggggacccttgattgttctttctttttcgctat<br>tgtaaaattcatgttatatggagggggcaaagttttcagggtgttgtttagaatgggaagatgtcccttgtatcaccatggaccctcatgataattttgtttctttcactttctactctgttgacaaccattgtctcctcttattttcttttcattttctgtaactttttcgttaaactttagcttgcatttgtaacgaattttaaattcacttttgtttatttgtcagattgtaagtactttctctaatcac<br>ttttttttcaaggcaatcagggtatattatattgtacttcagcacagttttagagaacaattgttataattaaatgataaggtagaatatttctgcatataaattctggctggcgtggaaatattcttattggtagaaacaactacatcctggtcatcatcctgcctttctctttatggttacaatgatatacactgtttgagatgaggataaaatactctgagtccaaaccgggccc<br>ctctgctaaccatgttcatgccttcttctttttcctacagctcctgggcaacgtgctggttattgtgctgtctcatcattttggcaaagaattgtaatacgactcactatagggcgaattcggatcccccggtagttaattaaaaattagggtgcaagatcatcgataacaagtttgtacaaaaaagcaggcttcgccaccatggccgccagcaacggcggagtgatgtac<br>cagtcattcctgaccatcatcatcctggtgatcatgaccgagggccaaatccattggggcaacctgagcaagatcggcatcgtgggaacaggcagcgctagctataaggtgatgaccagacctaatcaccaatacctggtgatcaaactgatgcctaacgtgaccatgattgacaactgtacacggaccgaggtgaccgagtacagaaagctgctgaaa<br>accgtgcttgagcctgtgaaaaatgccctgaccgtgatcaccaagaacatcaagcctatccagagcctgtatccgtatgatgtgccggattatgcgaccacaagcagacggagcaagaggtttgccggcgtggtgctggcaggagccgccctgggagtggcaaccgccgcccagatcacagccggcatcgccctgcaccagtccatgctgaactctcag<br>gccatcgataatctgagggcctccctggagacaacaaaccaggcaatcgaggcaatcaggcaggcaggacaggagatgatcctggccgtgcagggcgtgcaggactatatcaacaatgagctgatcccttctatgaatcagctgagctgcgatctgatcggccagaagctgggcctgaagctgctgcggtactataccgagatcctgtccctgttcggac<br>cttctctgcgggaccccatcagcgccgagatctccatccaggccctgtcctatgcactgggaggcgacatcaacaaggtgctggagaagctgggctactctggcggcgatctgctgggcatcctggagagccggggcatcaaggccagaatcacccacgtggatacagagtcttacttcatcgtgctgagcatcgcctaccccacactgtccgagatcaag<br>ggcgtgatcgtgcacaggctggagggcgtgagctataatatcggctcccaggagtggtacaccacagtgcctaagtatgtggccacccagggctacctgatctctaacttcgacgagtcctcttgtacctttatgccagagggcacagtgtgctcccagaatgccctgtacccaatgtctcccctgctgcaggagtgtctgcggggctctaccaagagctgcgc<br>cagaacactggtgagcggctccttcggcaatcggttcatcctgtcccagggcaacctgatcgccaattgtgcctctatcctgtgcaagtgctataccacaggcaccatcatcaaccaggaccccgataagatcctgacatacatcgcagcagaccactgccctgtggtggaagtgaatggcgtgaccatccaagtgggcagccggagatacccagacgcc<br>gtgtatctgcacaggatcgatctgggaccacctatctccctggagcgcctggatgtgggcacaaacctgggcaatgccatcgccaagctggaggacgccaaggagctgctggagagctccgatcagatcctgaggagcatgaagggcctgtctagcacctccatcgtgtacatcctgatcgccgtgtgcctgggaggcctgatcggaatccccgccctgat<br>ctgctgttgcaggggccgctgagacccagctttcttgtacaaagtggtactagtgtgaaatagacatcagaattaagaaaaacgtagggtccaagtggttccccgttatggactcgctatctgtcaaccagatcttataccctgaagttcacctagatagcccgatagttaccaataagatagtagccatcctggagtatgctcgactgtttaaacctgcaggcatgc<br>aagctgatctttttccctctgccaaaaattatggggacatcatgaagccccttgagcatctgacttctggctaataaaggaaatttattttcattgcaatagtgtgttggaattttttgtgtctctcactcggaaggacatatgggagggcaaatcatttaaaacatcagaatgagtatttggtttagagtttggcaacatatgcccatatgctggctgccatgaacaaaggtt<br>ggctataaagaggtcatcagtatatgaaacagccccctgctgtccattccttattccatagaaaagccttgacttgaggttagattttttttatattttgttttgtgttatttttttctttaacatccctaaaattttccttacatgttttactagccagatttttcctcctctcctgactactcccagtcatagctgtccctcttctcttatggagatccctcgaggagctttttgcaaaagccta<br>ggcctccaaaaaagcctcttcactacttctggaatagctcagaggccgaggcggcctcggcctctgcataaataaaaaaaattagtcagccatggggcggagaatgggcggaactgggcggagttaggggcgggatgggcggagttaggggcgggactatggttgctgactaattgctgcattaatgaatcggccaacgcgcggggagaggcggtttg<br>cgtattgggcgctcttccgcttcctcgctcactgactcgctgcgctcggtcgttcggctgcggcgagcggtatcagctcactcaaaggcggtaatacggttatccacagaatcaggggataacgcaggaaagaacatgtgagcaaaaggccagcaaaaggccaggaaccgtaaaaaggccgcgttgctggcgtttttccataggctccgcccccctgacga<br>gcatcacaaaaatcgacgctcaagtcagaggtggcgaaacccgacaggactataaagataccaggcgtttccccctggaagctccctcgtgcgctctcctgttccgaccctgccgcttaccggatacctgtccgcctttctcccttcgggaagcgtggcgctttctcatagctcacgctgtaggtatctcagttcggtgtaggtcgttcgctccaagctgggctgtgt<br>gcacgaaccccccgttcagcccgaccgctgcgccttatccggtaactatcgtcttgagtccaacccggtaagacacgacttatcgccactggcagcagccactggtaacaggattagcagagcgaggtatgtaggcggtgctacagagttcttgaagtggtggcctaactacggctacactagaagaacagtatttggtatctgcgctctgctgaagccagtt<br>accttcggaaaaagagttggtagctcttgatccggcaaacaaaccaccgctggtagcggtggtttttttgtttgcaagcagcagattacgcgcagaaaaaaaggatctcaagaagatcctttgatcttttctacggggtctgacgctcagtggaacgaaaactcacgttaagggattttggtcatgagattatcaaaaaggatcttcacctagatccttttaaattaaa<br>aatgaagttttaaatcaatctaaagtatatatgagtaaacttggtctgacagttaccaatgcttaatcagtgaggcacctatctcagcgatctgtctatttcgttcatccatagttgcctgactccccgtcgtgtagataactacgatacgggagggcttaccatctggccccagtgctgcaatgataccgcgagacccacgctcaccggctccagatttatcagcaata<br>aaccagccagccggaagggccgagcgcagaagtggtcctgcaactttatccgcctccatccagtctattaattgttgccgggaagctagagtaagtagttcgccagttaatagtttgcgcaacgttgttgccattgctacaggcatcgtggtgtcacgctcgtcgtttggtatggcttcattcagctccggttcccaacgatcaaggcgagttacatgatcccccatgt<br>tgtgcaaaaaagcggttagctccttcggtcctccgatcgttgtcagaagtaagttggccgcagtgttatcactcatggttatggcagcactgcataattctcttactgtcatgccatccgtaagatgcttttctgtgactggtgagtactcaaccaagtcattctgagaatagtgtatgcggcgaccgagttgctcttgcccggcgtcaatacgggataataccgcgccac<br>atagcagaactttaaaagtgctcatcattggaaaacgttcttcggggcgaaaactctcaaggatcttaccgctgttgagatccagttcgatgtaacccactcgtgcacccaactgatcttcagcatcttttactttcaccagcgtttctgggtgagcaaaaacaggaaggcaaaatgccgcaaaaaagggaataagggcgacacggaaatgttgaatactcata<br>ctcttcctttttcaatattattgaagcatttatcagggttattgtctcatgagcggatacatatttgaatgtatttagaaaaataaacaaataggggttccgcgcacatttccccgaaaagtgccacctgacgtctaagaaaccattattatcatgacattaacctataaaaataggcgtatcacgaggccctttcgtctcgcgcgtttcggtgatgacggtgaaaacctctg<br>acacatgcagctcccggagacggtcacagcttgtctgtaagcggatgccgggagcagacaagcccgtcagggcgcgtcagcgggtgttggcgggtgtcggggctggcttaactatgcggcatcagagcagattgtactgagagtgcaccattcgacgctctcccttatgcgactcctgcattaggaagcagcccagtagtaggttgaggccgttgagcacc<br>gccgccgcaaggaatggtctggcttatcgaaattaatacgactcactatagggagaccg |

| | | |
|---|---|---|
| pLX311-ahCD3/hCD80 | Lentiviral transfer vector coding for an anti-human CD3 antibody and human CD80 to provide costimulation on the surface of HEK293 producer cells | ctcgaggcctgcaggtgcaaagatggataaagttttaaacagagaggaatctttgcagctaatggaccttctaggtcttgaaaggagtgggaattggctccggtgcccgtcagtgggcagagcgcacatcgcccacagtccccgagaagttggggggaggggtcggcaattgaaccggtgcctagagaaggtggcgcggggtaaactgggaaagtgat<br>gtcgtgtactggctccgcctttttcccgagggtgggggagaaccgtatataagtgcagtagtcgccgtgaacgttctttttcgcaacgggtttgccgccagaacacaggtaagtgccgtgtgtggttcccgcgggcctggcctctttacgggttatggcccttgcgtgccttgaattacttccacctggctgcagtacgtgattcttgatcccgagcttcgggttggaagtg<br>ggtgggagagttcgaggccttgcgcttaaggagccccttcgcctcgtgcttgagttgaggcctggcctgggcgctggggccgccgcgtgcgaatctggtggcaccttcgcgcctgtctcgctgctttcgataagtctctagccatttaaaatttttgatgacctgctgcgacgctttttttctggcaagatagtcttgtaaatgcgggccaagatctgcacactggtatttc<br>ggtttttggggccgcgggcggcgacggggcccgtgcgtcccagcgcacatgttcggcgaggcggggcctgcgagcgcggccaccgagaatcggacgggggtagtctcaagctggccggcctgctctggtgcctggcctcgcgccgccgtgtatcgccccgccctgggcggcaaggctggcccggtcggcaccagttgcgtgagcggaaagatggcc<br>gcttcccggccctgctgcagggagctcaaaatggaggacgcggcgctcgggagagcgggcgggtgagtcacccacacaaaggaaaagggcctttccgtcctcagccgtcgcttcatgtgactccacggagtaccgggcgccgtccaggcacctcgattagttctcgagcttttggagtacgtcgtctttaggttggggggaggggttttatgcgatggagttt<br>ccccacactgagtgggtggagactgaagttaggccagcttggcacttgatgtaattctccttggaatttgccctttttgagtttggatcttggttcattctcaagcctcagacagtggttcaaagtttttttcttccatttcaggtgtcgtgaggctagcatcgattgatcaacaagtttgtacaaaaaagcaggcttcgccaccatggccttaccagtgaccgccttgctcctg<br>ccgctggccttgctgctccacgccgccaggccgatgggatggtcatgtatcatactgttcctggtggcgacagctaccggcgtgcatagccaagtgcagctccagcagtccggcgccgagcttgctcgcccgggcgcaagtgtaaagatgtcctgcaaggctagcggctacactttcacccgttataccatgcactgggtgaaacagcgacctggtcagggtt<br>tagagtggatcggctacataaaccccagccggggtacactaattataatcaaaaatttaaagacaaggccacactgacgacagataagtcctcatctaccgcctatatgcagcttagctctcttacctccgaagacagcgctgtgtattactgcgcaaggtactatgatgaccattactgtctagactactgggggcaaggtaccacactgacagtgagctct<br>ggaggcgggggggagtggaggtgggggcagcggcggtggcggaagtgatatcgttctgacccagagcccagctattatgtctgcctctcccggcgagaaggtgactatgacgtgttccgcatcatccagcgttagttacatgaattggtatcagcaaaagagtgggacctctcctaagcggtggatctacgatacgtcaaagctcgcctccggcgtaccagc<br>gcacttcagagggtcgggttccgggacaagctatagtctgactatttcagggatggaagccgaagatgccgcaacctattactgccagcagtggtcctcaaacccatttactttcggaagcggaacaaaactcgaaatcaacaggggatccgagcagaaactgatttctgaggaagacctgaacgctgtcggccaagacactcaggaggtcatagttgtcc<br>cccactctctcccttttaaagtggtggtcatttcggccatcctggcactggtcgttttgactattattagtctgatcattttgatcatgttgtggcagaagaaaccaagaggatcaatccccaatcctttactcggactagatggatccgcaacaaacttctctctgctgaaacaagccggagatgtcgaagagaatcctggaccggctagcatgggccacacgcggcg<br>gcagggcaccagcccttccaagtgtccatatttgaacttcttccagcttttagtgctcgctggcttgtcccatttctgcagtggcgttattcatgtgaccaaagaggtcaaagaggtcgccacccttagttgcggacataatgtctccgtggaggaactggcgcagaccaggatttactggcagaaggagaaaaagatggtgcttacaatgatgtccggggacatga<br>atatttggcctgaatataaaaaccgaaccatttcgacatcactaataatctcagcatcgtgattctggctctaaggccctcagacgagggtacatacgagtgtgtagtcctgaaatatgaaaaggatgcattcaagcgtgagcacctggcagaggtgactctatcagttaaggccgactttcccacgcccagtatctctgatttcgagatccctacttccaacatta<br>gacgcatcatctgtagcactagcggggggtttccggaaccccatctgtcttggctcgaaaatggagaagagttgaatgccataaacacaacagtgagccaagatccggagaccgagctgtacgccgtatcttctaaactggatttcaacatgacgacaaaccactcattcatgtgcttaatcaagtacggtcacctgcgggtcaaccagacatttaattggaa<br>cactactaagcaagaacactttccagacaatctgctgccatcatgggcaataaccctcattagcgtgaacggaatctttgttatatgttgcctgacctattgttttgctccacggtgcagagaacgcaggagaaacgaaaggctcagacgagagtcggtgcgccctgtttaagacccagctttcttgtacaaagtggttgatatcggtaagcctatccctaaccctct<br>cctcggtctcgattctacgtagtaatgaactagtaccggttaagtcgacaatcaacgcgttaagtcgacaatcaacctctggattacaaaatttgtgaaagattgactggtattcttaactatgttgctccttttacgctatgtggatacgctgctttaatgcctttgtatcatgctattgcttcccgtatggctttcattttctcctccttgtataaatcctggttgctgtctctttatgag<br>gagttgtggcccgttgtcaggcaacgtggcgtggtgtgcactgtgtttgctgacgcaacccccactggttggggcattgccaccacctgtcagctcctttccgggactttcgctttccccctccctattgccacggcggaactcatcgccgcctgccttgcccgctgctggacaggggctcggctgttgggcactgacaattccgtggtgttgtcggggaaatcatcgtc<br>ctttccttggctgctcgcctgtgttgccacctggattctgcgcgggacgtccttctgctacgtcccttcggccctcaatccagcggaccttccttcccgcggcctgctgccggctctgcggcctcttccgcgtcttcgccttcgccctcagacgagtcggatctccctttgggccgcctccccgcgtcgactttaagaccaatgacttacaaggcagctgtagatcttagcc<br>actttttaaaagaaaaggggggactggaagggctaattcactcccaacgaagacaagatctgctttttgcttgtactgggtctctctggttagaccagatctgagcctgggagctctctggctaactagggaacccactgcttaagcctcaataaagcttgccttgagtgcttcaagtagtgtgtgcccgtctgttgtgtgactctggtaactagagatccctcagaccc<br>ttttagtcagtgtggaaaatctctagcagtacgtatagtagttcatgtcatcttattattcagtatttataacttgcaaagaaatgaatatcagagagtgagaggaacttgtttattgcagcttataatggttacaaataaagcaatagcatcacaaatttcacaaataaagcatttttttcactgcattctagttgtggtttgtccaaactcatcaatgtatcttatcatgtctggct<br>ctagctatcccgcccctaactccgcccatcccgcccctaactccgcccagttccgcccattctccgccccatggctgactaattttttttatttatgcagaggccgaggccgcctcggcctctgagctattccagaagtagtgaggaggcttttttggaggcctagggacgtacccaattcgccctatagtgagtcgtattacgcgcgctcactggccgtcgttttacaac<br>gtcgtgactgggaaaaccctggcgttacccaacttaatcgccttgcagcacatccccctttcgccagctggcgtaatagcgaagaggcccgcaccgatcgcccttcccaacagttgcgcagcctgaatggcgaatgggacgcgccctgtagcggcgcattaagcgcggcgggtgtggtggttacgcgcagcgtgaccgctacacttgccagcgccctagc<br>gcccgctcctttcgctttcttcccttcctttctcgccacgttcgccggctttccccgtcaagctctaaatcgggggctccctttagggttccgatttagtgctttacggcacctcgaccccaaaaaacttgattagggtgatggttcacgtagtgggccatcgccctgatagacggtttttcgccctttgacgttggagtccacgttctttaatagtggactcttgttccaaactgg<br>aacaacactcaaccctatctcggtctattcttttgatttataagggattttgccgatttcggcctattggttaaaaaatgagctgatttaacaaaaatttaacgcgaattttaacaaaatattaacgcttacaatttaggtggcacttttcggggaaatgtgcgcggaacccctatttgtttatttttctaaatacattcaaatatgtatccgctcatgagacaataaccctgataa<br>atgcttcaataatattgaaaaaggaagagtatgagtattcaacatttccgtgtcgcccttattcccttttttgcggcattttgccttcctgtttttgctcacccagaaacgctggtgaaagtaaaagatgctgaagatcagttgggtgcacgagtgggttacatcgaactggatctcaacagcggtaagatccttgagagttttcgccccgaagaacgttttccaatgatga<br>gcacttttaaagttctgctatgtggcgcggtattatcccgtattgacgccgggcaagagcaactcggtcgccgcatacactattctcagaatgacttggttgagtactcaccagtcacagaaaagcatcttacggatggcatgacagtaagagaattatgcagtgctgccataaccatgagtgataacactgcggccaacttacttctgacaacgatcggaggac<br>cgaaggagctaaccgcttttttgcacaacatgggggatcatgtaactcgccttgatcgttgggaaccggagctgaatgaagccataccaaacgacgagcgtgacaccacgatgcctgtagcaatggcaacaacgttgcgcaaactattaactggcgaactacttactctagcttcccggcaacaattaatagactggatggaggcggataaagttgcagga<br>ccacttctgcgctcggcccttccggctggctggtttattgctgataaatctggagccggtgagcgtgggtctcgcggtatcattgcagcactggggccagatggtaagccctcccgtatcgtagttatctacacgacggggagtcaggcaactatggatgaacgaaatagacagatcgctgagataggtgcctcactgattaagcattggtaactgtcagaccaa<br>gtttactcatatatactttagattgatttaaaacttcatttttaatttaaaaggatctaggtgaagatcctttttgataatctcatgaccaaaatcccttaacgtgagttttcgttccactgagcgtcagaccccgtagaaaagatcaaaggatcttcttgagatcctttttttctgcgcgtaatctgctgcttgcaaacaaaaaaaccaccgctaccagcggtggtttgtttgccgg<br>atcaagagctaccaactctttttccgaaggtaactggcttcagcagagcgcagataccaaatactgttcttctagtgtagccgtagttaggccaccacttcaagaactctgtagcaccgcctacatacctcgctctgctaatcctgttaccagtggctgctgccagtggcgataagtcgtgtcttaccgggttggactcaagacgatagttaccggataaggcgcag<br>cggtcgggctgaacggggggttcgtgcacacagcccagcttggagcgaacgacctacaccgaactgagatacctacagcgtgagctatgagaaagcgccacgcttcccgaagggagaaaggcggacaggtatccggtaagcggcagggtcggaacaggagagcgcacgagggagcttccagggggaaacgcctggtatctttatagtcctgtcg<br>ggtttcgccacctctgacttgagcgtcgatttttgtgatgctcgtcaggggggcggagcctatggaaaaacgccagcaacgcggcctttttacggttcctggccttttgctggccttttgctcacatgttctttcctgcgttatcccctgattctgtggataaccgtattaccgcctttgagtgagctgataccgctcgccgcagccgaacgaccgagcgcagcgagtcagt<br>gagcgaggaagcggaagagcgcccaatacgcaaaccgcctctccccgcgcgttggccgattcattaatgcagctggcacgacaggtttcccgactggaaagcgggcagtgagcgcaacgcaattaatgtgagttagctcactcattaggcaccccaggctttacactttatgcttccggctcgtatgttgtgtggaattgtgagcggataacaatttcacaca<br>ggaaacagctatgaccatgattacgccaagcgcgcaattaaccctcactaaagggaacaaaagctggagctgcaagcttaatgtagtcttatgcaatactcttgtagtcttgcaacatggtaacgatgagttagcaacatgccttacaaggagagaaaaagcaccgtgcatgccgattggtggaagtaaggtggtacgatcgtgccttattaggaaggcaac<br>agacgggtctgacatggattggacgaaccactgaattgccgcattgcagagatattgtatttaagtgcctagctcgatacataaacgggtctctctggttagaccagatctgagcctgggagctctctggctaactagggaacccactgcttaagcctcaataaagcttgccttgagtgcttcaagtagtgtgtgcccgtctgttgtgtgactctggtaactagagatc<br>cctcagacccttttagtcagtgtggaaaatctctagcagtggcgcccgaacagggacttgaaagcgaaagggaaaccagaggagctctctcgacgcaggactcggcttgctgaagcgcgcacggcaagaggcgaggggcggcgactggtgagtacgccaaaaattttgactagcggaggctagaaggagagagatgggtgcgagagcgtcagtat<br>taagcgggggagaattagatcgcgatgggaaaaaattcggttaaggccagggggaaagaaaaaatataaattaaaacatatagtatgggcaagcagggagctagaacgattcgcagttaatcctggcctgttagaaacatcagaaggctgtagacaaatactgggacagctacaaccatcccttcagacaggatcagaagaacttagatcattatata<br>atacagtagcaaccctctattgtgtgcatcaaaggatagagataaaagacaccaaggaagctttagacaagatagaggaagagcaaaacaaaagtaagaccaccgcacagcaagcggccgctgatcttcagacctggaggaggagatatgagggacaattggagaagtgaattatataaatataaagtagtaaaaattgaaccattaggagtagca<br>cccaccaaggcaaagagaagagtggtgcagagagaaaaaagagcagtgggaataggagctttgttccttgggttcttgggagcagcaggaagcactatgggcgcagcgtcaatgacgctgacggtacaggccagacaattattgtctggtatagtgcagcagcagaacaatttgctgagggctattgaggcgcaacagcatctgttgcaactcacagtc<br>tggggcatcaagcagctccaggcaagaatcctggctgtggaaagatacctaaaggatcaacagctcctggggatttggggttgctctggaaaactcatttgcaccactgctgtgccttggaatgctagttggagtaataaatctctggaacagatttggaatcacacgacctggatggagtgggacagagaaattaacaattacacaagcttaatacactcctt<br>aattgaagaatcgcaaaaccagcaagaaaagaatgaacaagaattattggaattagataaatgggcaagtttgtggaattggtttaacataacaaattggctgtggtatataaaattattcataatgatagtaggaggcttggtaggtttaagaatagtttttgctgtactttctatagtgaatagagttaggcagggatattcaccattatcgtttcagacccacctccc<br>aaccccgaggggacccatgcattgcatctcaattagtcagcaaccaggtgtggaaagtccccaggctccccagcaggcagaagtatgcaaagcatgcgtctcaattagtcagcaaccatagtcccgcccctaactccgcccatcccgcccctaactccgcccagttccgcccattctccgccccatggctgactaattttttttatttatgcagaggccgaggc<br>cgcctcggcctctgagctattccagaagtagtgaggaggcttttttggaggcctaggcttttgcaaaaagctttctagaggtaccaccatggccaagcctttgtctcaagaagaatccaccctcattgaaagagcaacggctacaatcaacagcatccccatctctgaagactacagcgtcgccagcgcagctctctctagcgacggccgcatcttcactggtgt<br>caatgtatatcattttactgggggaccttgtgcagaactcgtggtgctgggcactgctgctgctgcggcagctggcaacctgacttgtatcgtcgcgatcggaaatgagaacaggggcatcttgagcccctgcggacggtgccgacaggtgcttctcgatctgcatcctgggatcaaagccatagtgaaggacagtgatggacagccgacggcagttgggattc<br>gtgaattgctgccctctggttatgtgtgggagggcctgcagctgcagtagtaaggcgcgccgttaacgaattctagatcttgagacaaatggcagtattcatccacaattttaaaagaaaaggggggattggggggtacagtgcaggggaaagaatagtagacataatagcaacagacatacaaactaaagaattacaaaaacaaattacaaaaattcaa<br>aattttcgggtttattacagggacagcagagatccactttggcgccgg |
